# Viral disease outcomes are indistinguishable between experimentally infected bats and rodents

**DOI:** 10.1101/2025.08.28.672855

**Authors:** Maxwell J. Farrell, Samantha K. Tucker, Nardus Mollentze, Daniel G. Streicker

## Abstract

A common explanation for bats being a conspicuous source of zoonotic viruses is their purported ability to coexist with viruses without suffering overt disease. This belief has catalyzed the discovery of unique features of bat immune systems which may have translational value as broad-spectrum antivirals, particularly if they evolved as a byproduct of bats’ unique life history rather than through conventional co-evolutionary processes. Surprisingly, whether bats compared to other host groups suffer less disease from co-evolved viruses or from viruses generally has not been formally assessed. Here, synthesizing eighty-six years of experimental infections, involving 54 viruses, 85 host species, and over 5,600 animals, we document disease in bats following inoculation by taxonomically diverse viruses, including ones that are relatively benign in humans. The occurrence of overt disease, the likelihood of mortality, and the severity of disease in bats were indistinguishable from those experienced by rodents, another group associated with many zoonotic viruses. These patterns were consistent when considering only bat-associated viruses inoculated into bats and rodent-associated viruses inoculated into rodents and among inoculations which lacked shared co-evolutionary history. Instead, disease outcomes following infection were shaped by experimental design, viral host range, and evolutionary context. Unexceptional disease in bats from novel or co-evolved infections suggests that order-level host life history traits such as flight have inconsistent consequences for antiviral immunity and highlights the need to evaluate the functional properties of putatively unique features of bat immunity in vivo. These results do not exclude the possibility of developing broader-acting or more potent antivirals from bat immune systems nor do they diminish the potential value of mechanistic insights into bat immunity. However, they do not support the premise underlying the idea that bats will be an unusually rich source of future biomedical breakthroughs.

---

The purported rarity of overt disease in bats infected with viruses that are pathogenic in other mammals has fueled the widespread perception that bats have enhanced capabilities to limit the severity of active viral infections (Baker et al., 2013; Brook and Dobson, 2015; Calisher et al., 2006; O’Shea et al., 2014; Shen et al., 2010; Wang et al., 2011; Wong et al., 2007). Efforts to explain this phenomenon have identified numerous features of bat immune systems that appear to differ from other mammals (Banerjee et al., 2020; Irving et al., 2021; Morales et al., 2025). Most enticingly, some aspects of bat immune systems such as their capacity to limit inflammation are hypothesized to have evolved as a byproduct of the energetic demands of flight which are unique to bats among mammals. If so, these features might protect against diverse viruses, potentially representing candidates for breakthrough, broad-spectrum antivirals (Baid et al., 2024; Dolgin, 2023; O’Shea et al., 2014; Shen et al., 2010). Supporting this possibility, bat derived ASC2 (apoptosis-associated speck-like protein containing a CARD) appears to dampen inflammation caused by both coronaviruses and orthomyxoviruses, and reduced inflammatory cytokines caused by orthoreovirus and flaviviruses (Ahn et al., 2023). Alternatively, apparent limitation of virus-induced disease in bats may largely reflect conventional adaptations to the specific viral groups that bats co-evolved with. Indeed, neither taxon-specific immune features nor sub-clinical infections from viruses that cause disease in other species are unique to bats (Shaw et al., 2017; Farrell and Davies, 2019). In this case, interactions between bat life history and their co-evolved viruses might still lead bats to have more potent disease control mechanisms against these particular viruses when compared to other host groups, but these defenses would be unlikely to offer protection against a broad spectrum of viruses.

Distinguishing whether bats exhibit more potent or taxonomically comprehensive suppression of viral disease compared to other mammals underpins research into the therapeutic potential of bat immune systems and the nature of bats as special zoonotic reservoirs, but remains untested. Observations of infection outcomes in wild bats are largely anecdotal; for example, viruses detected following mortality events with unknown etiology, or viruses detected in apparently healthy bats whose ultimate fates are unobservable (Kemenesi et al., 2018; Kohl et al., 2020). Field observations are further complicated by the small body sizes and cryptic life histories of bats, which limit detections of virus-induced morbidity and mortality to the most conspicuous events. In contrast, experimental inoculations of captive animals can confidently monitor infection outcomes following known exposure events. Such infections have revealed limited disease during infection by several viruses that can cause high mortality in humans (e.g., Marburg virus, Nipah virus, Hendra virus) (Calisher et al., 2006; Baker et al., 2013; Guth et al., 2022). Determining whether these observations represent the exception or the rule requires synthesizing outcomes across a broader range of viruses. Until now, such analyses have been precluded by the absence of systematic datasets on the disease outcomes of experimental infections.

We examine the current state of evidence that bats experience dampened virus-induced disease by extracting infection outcome data from eighty-six years of experimental viral infection studies. The severity of disease outcomes may be influenced by both tolerance (i.e., the ability to limit disease at a given viral load) and resistance (i.e., the capacity to inhibit infection upon exposure). Both phenomena likely occur in bats and may select for increased viral pathogenicity. Tolerance has been of particular interest since it has been hypothesized to be heightened in bats relative to other mammals and to promote zoonotic transmission by enabling bat survival and continued viral shedding despite high viral loads (Brook et al., 2023; Guth et al., 2022; Irving et al., 2021; Mandl et al., 2018). We note that despite the informal use of the term “tolerance” in the bat virus literature when referring to the outcomes of experimental and natural viral infections, formally measuring tolerance would require non-invasive quantification of viral loads in physiologically relevant tissues and prohibitively expensive long-term monitoring of fitness costs in infected hosts. It is therefore unclear that tolerance has or can be measured from experimental or natural infections (Medzhitov et al., 2012). We therefore frame our analysis around the occurrence and severity of outcomes following viral infection based on the extent of author-reported disease (including clinical and sub-clinical pathologies). Because we are interested in disease outcomes in compatible host-virus pairs, we restrict our analysis to inoculations in which the original authors deemed hosts to be susceptible (See S1 for detailed methods).

We contrast bats with rodents, another species-rich mammalian order (Upham et al., 2019) that hosts a large number of zoonotic viruses (some asymptomatically; Pereira et al., 2023). However, on average rodents have shorter life expectancies and higher reproductive rates than bats, which some theory has suggested may make them less likely to evolve mechanisms to suppress to viral disease, though other models have suggested that these relationships may be complex (Albery and Becker, 2021; Hayman, 2019; Han et al., 2015; Luis et al., 2013; Boots et al., 2013; Miller et al., 2007). Importantly, our dataset included viruses that are presumed to have co-evolved with bats and rodents (homologous inoculations), and those that are currently unlinked to either mammalian order (heterologous inoculations). This allowed us to define and test two main hypotheses on how disease outcomes in bats differ from other groups: that non-specific antiviral mechanisms enable bats to reduce disease severity irrespective of co-evolutionary relationships (generalized response) and that bats suppress disease from their co-evolved viruses better than other host groups due to the combined selection pressures of bat life history constraints and infection by specific viral groups (heightened co-evolutionary response).

Beyond intrinsic host properties, variation in disease outcomes may reflect eco-evolutionary characteristics of viruses (Galen et al., 2022; Leggett et al., 2013; Agueldo Romero and Elena, 2008). For example, viruses that infect a broader diversity of host species may replicate poorly in any single species, making them less pathogenic than host-specific viruses (Antonovics et al., 2013; Leggett et al., 2013). Viruses may also cause increased pathogenicity in novel hosts which lack co-evolved defenses, or alternatively disease severity may be reduced in novel hosts due to poor adaptation and reduced replication (Antonovics et al., 2013; Leggett et al., 2013). Therefore, in our models of disease outcomes we included measures of viral host range and proxies for the extent of co-evolution with inoculated host groups. Experimental design choices such as the dose and route of inoculation, the method and extent of virus passage prior to inoculation, and the number of individuals included in each study may also influence the likelihood that an inoculation leads to detectable disease. We therefore adjusted for these experimental details in our models where possible, or carried out independent analyses to ensure they were unlikely to explain our main findings. Finally, we investigated the relationship between human case fatality rates and disease outcomes in experimentally inoculated bats and rodents to assess whether either host group was systematically inoculated with viruses that are more pathogenic in a third host group, and to explore the suggestion that bats consistently experience reduced disease severity from viruses that are highly pathogenic to humans.

## Diverse viruses cause overt disease in bats and rodents

We identified 112 studies published between 1936 – 2022 that reported disease outcomes following experimental inoculations (S3) of 54 viruses from fifteen viral families (53 RNA viruses; 1 DNA virus) into 85 susceptible host species (Figs. 1, S8; Table S3). These represented 149 host-virus pairs and a total of 5,622 inoculated animals. We excluded laboratory-bred rodents as they may lack the full immune competency of their wild counterparts (Abolins et al., 2017). Across host-virus combinations, bat-virus pairs included a larger number of viruses than rodent-virus pairs, but fewer bat species were subjected to experimental inoculation, indicating frequent use of a smaller number of model bat species (Fig. S9). Nevertheless, while some host clades were not subjected to experimental inoculations, our dataset comprised a phylogenetically representative sample of both bats and rodents (Fig. S8), capturing variation in host life history traits hypothesized to influence disease outcomes (Hayman, 2019; Fig. S10). For each host-virus pair per study, we recorded author-reported signs of disease and the occurrence of virus-induced mortality. We further converted signs of disease into a five point severity scale which ranged from no overt disease to death. We therefore consider three distinct disease outcome metrics: disease presence, mortality, and severity (S1.2). Across this dataset, disease occurred in 48% (40/83) of rodent-virus pairs and 73% (48/66) of bat-virus pairs (Fig. 1). Among all host-virus pairs, viruses from fifteen families caused disease (Table S5), with severe disease observed across the host phylogeny rather than being clustered within select clades (Fig. 1). Filoviruses and paramyxoviruses are commonly cited examples of highly pathogenic viruses that rarely cause clinical disease in bats. Consistent with this idea, all tested filoviruses caused at most subclinical disease in bats and most paramyxoviruses caused no disease or subclinical disease in bats. However, two out of six bat-paramyxovirus pairs (specifically Sosuga pararubulavirus in *Rousettus aegyptiacus* and Newcastle disease virus in *Myotis lucifugus*) caused severe clinical disease and death, respectively. Other notable virus-bat pairs that resulted in severe or lethal disease included Foot and mouth disease virus inoculated into *Desmodus rotundus*, Cocal vesiculovirus inoculated into *Myotis lucifugus*, Kaeng Khoi virus inoculated into *Chaerephon plicatus*, Tacaribe virus inoculated into *Carollia perspicillata* and *Artibeus jamaicensis*, Kyasanur Forest Disease Virus inoculated into *Rousettus leschenaultii* and *Cynopterus sphinx*, Tick-borne encephalitis virus inoculated into *Barbastella barbastellus* and *Plecotus auritus*, Dengue virus inoculated into *Artibeus literatus,* West Nile Virus inoculated into *Eptesicus fuscus* and all 9 included lyssavirus species inoculated into a variety of bat species (Table S5).

## Disease presence, mortality, and severity are indistinguishable between bats and rodents

Although raw data suggested similar disease outcomes between host orders (Fig. 1), this comparison did not control for differences in experimental design (e.g., dose, inoculation route, number of animals inoculated, etc.), host specificity, host-virus co-evolution, or the non-independence among taxa due to shared evolutionary histories. Indeed, the number of individual bats per study tended to be higher and more variable than the number of rodents (bats: median = 14.5, range 1 – 493; rodents: median = 6, range = 1 – 61), which, if unaccounted for, could inflate observations of clinical disease in bats. We therefore used hierarchical Bayesian models to examine host taxonomic effects alongside eco-evolutionary metrics reflecting the potential degree of virus adaptation to inoculated hosts, and the details of experimental design. At the host species level (i.e., the occurrence or maximum severity of disease among all individuals inoculated with a given virus within a given study; n=193 unique host-virus-study combinations, see S1.4), disease presence, mortality, and severity following viral inoculation remained indistinguishable between bats and rodents (the 80% credible interval for the effect of host order included zero; Fig. 2). The absence of host order effects was robust to removing lyssaviruses (S2.2.4), which represented 23% (45/193) of host-virus-study combinations in our dataset and, due to causing mortality, are sometimes considered the “exception to the rule” as viruses that cause severe disease in bats (Hayman, 2019). In contrast, hierarchical effects describing non-independence among hosts and viruses indicated that variation in disease outcomes was partially driven by variation among hosts and viruses that was not captured by other predictors in the model (S2.4).

**Figure 1:**
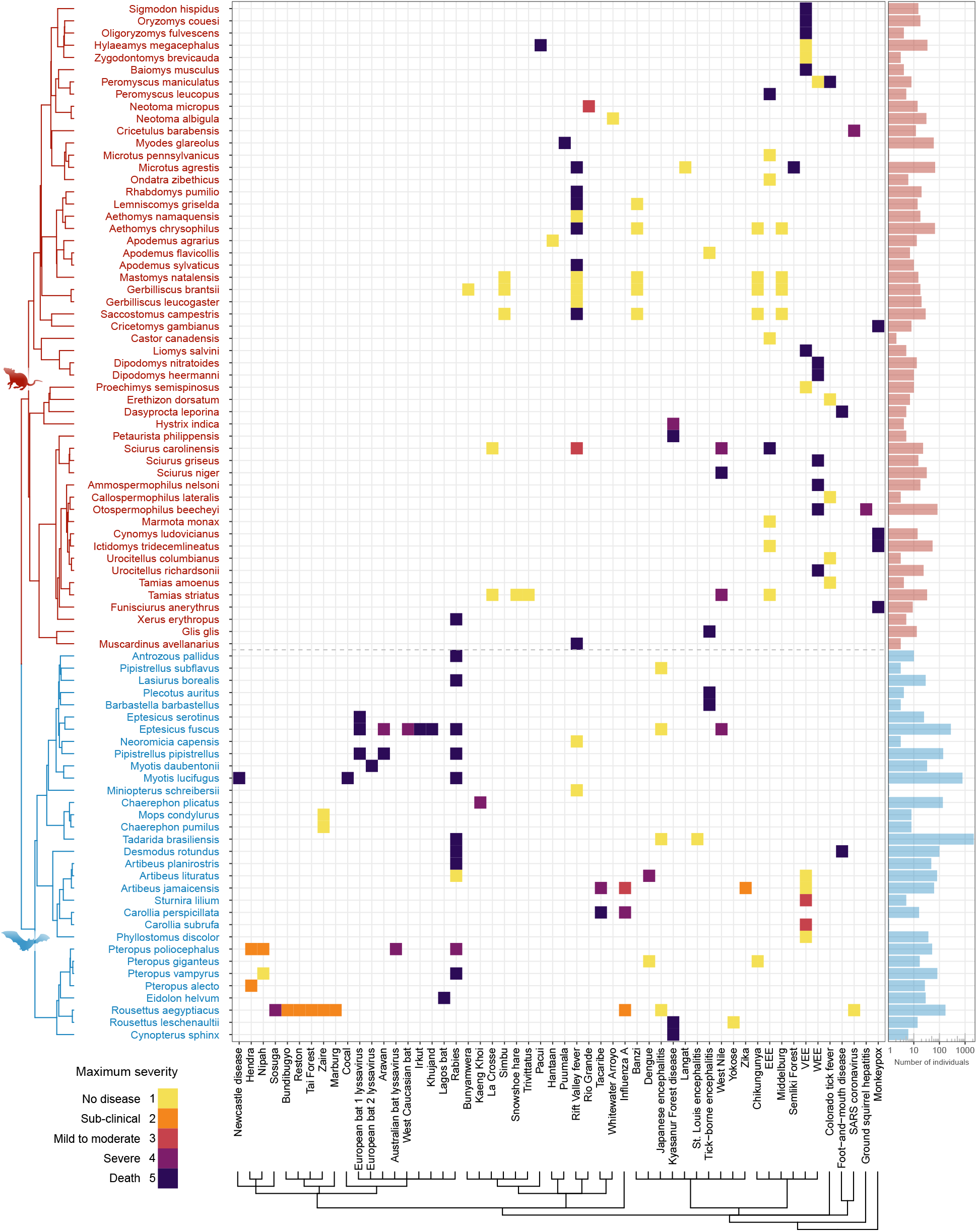
Maximum disease severity observed in experimental infections for each host-virus combination. The five point severity scale is calculated based on author reports of infection-induced disease and death (see S3 and S1.2.6 for details). Blank spaces indicate untested interactions. Hosts (rows) are grouped according to the Upham et al. (2019) mammal supertree. Viruses (columns) are grouped following the ICTV taxonomy. The barchart on the right shows the total number of host individuals per species included in experimental infections. For an alternative version depicting mean observed severity see Fig. S13. Raw data and scripts to produce the figure are available in the Data & Code supplement.

**Figure 2:**
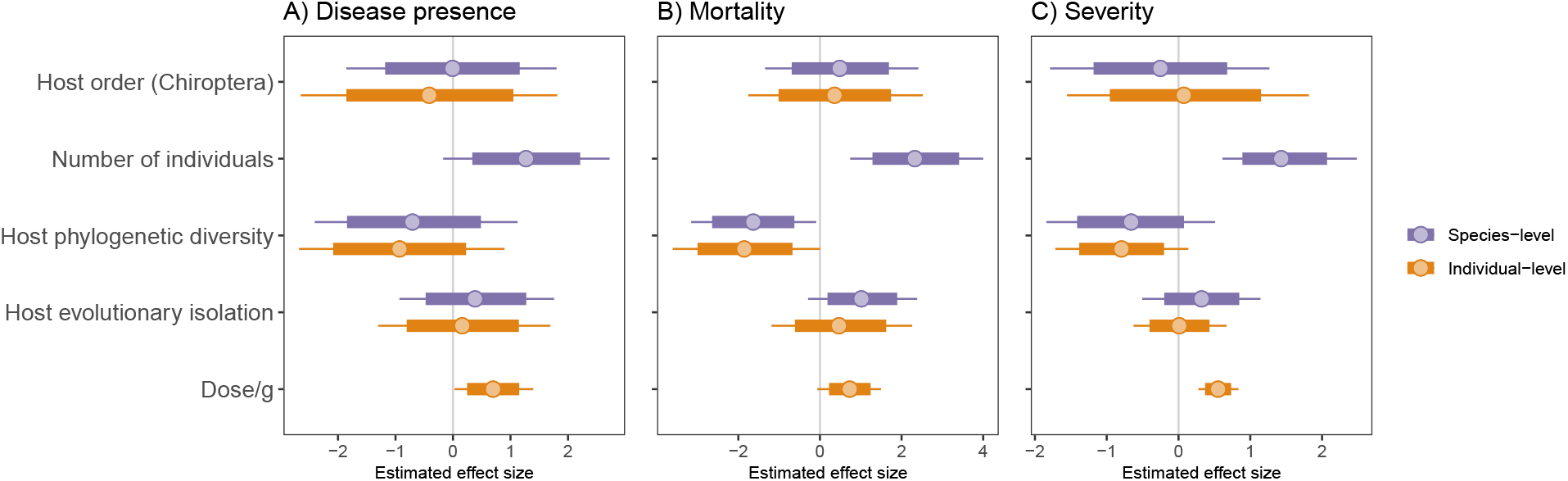
Estimated effect sizes for predictors in species-level and individual-level models of A) disease presence, B) mortality, and C) disease severity. Points represent posterior mean estimates, thick bars represent 80% credible intervals, thin bars represent 95% credible intervals. Predictors in both species- and individual-level models include host order (effect of Chiroptera with Rodentia as reference group), the standardized effect size of phylogenetic distance among all documented hosts per virus (“Host phylogenetic diversity”), the mean evolutionary distance from all documented host species to the inoculated host species (“Host evolutionary isolation”), and phylogenetic and non-phylogenetic hierarchical effects for host and virus species (S2.4). Species-level models additionally included the number of individuals per study (“Number of individuals”) and a hierarchical effect of article. Individual-level models additionally included inoculation dose adjusted for species body mass (“Dose/g”; see S2.2.5 for alternative approaches to model inoculation doses) and hierarchical effects for inoculation route (e.g., direct inoculation into the brain, oronasal inoculation, etc.) and the dose measurement units (e.g., Mouse LD50, TCID50, PFU, etc.). Positive host order effects would indicate more severe disease in bats compared to rodents.

Modeling host-virus interactions at the species-level maximized the diversity of hosts and viruses in our analyses by including studies that did not report data for individual animals. However, this approach disguised individual animal-level variation in disease outcomes and required omitting experimental conditions that varied within studies (e.g., dose and inoculation route). We therefore analyzed infection outcomes for 1,620 individual animals, spanning 44 viruses, 75 host species, and 128 host-virus pairs from 84 studies which reported sufficiently resolved data. Using a similar structure to our species-level models, we fit individual-level models for the three measures of disease outcome, but replaced the study-level continuous predictor (number of inoculated host individuals) with body mass-corrected dose and inoculation route (see S1.2). These models again found no difference between bats and rodents in any disease metric (Fig. 2).

## Contexts of infection shape disease outcomes

While our data and models do not support a host order effect, we did find evidence for several other factors theoretically predicted to influence the outcomes of infection. Among eco-evolutionary traits of viruses, infection by phylogenetic generalists (i.e., viruses that infect evolutionarily diverse host species) was less likely to result in disease, mortality, and severe disease (Fig. 2), consistent with the hypothesis that costs of generalism lead specialist viruses to cause more severe disease (Leggett et al., 2013).

To test whether either host order dependably experienced reduced disease when exposed to potentially novel viruses, we included an effect of host evolutionary isolation (HEI; Farrell and Davies, 2019), calculated as the mean phylogenetic distance from all known hosts to the inoculated host (Carlson et al. (2022); see S1.2.4 for details). As pathogens commonly infect closely related host species, this metric assumes that the phylogenetic centroid of known susceptible species reflects the most likely position of hosts to which a virus is best adapted, and that the distance from an inoculated host to this centroid approximates the extent of co-adaptation (Antonovics et al., 2013; Davies and Pedersen, 2008; Park et al., 2018; Farrell and Davies, 2019). Although estimated effect sizes were uncertain, mean effects of HEI were positively associated with disease outcomes, with greater certainty in species-level models of mortality (80% CIs excluded zero; S2.2; Fig. 2, Fig. S27). Although tentative, these results suggest that virus inoculations into distant relatives of currently documented susceptible species may more often result in more severe disease, a pattern previously reported for infection-induced mortality in humans and domesticated animals (Guth et al., 2019; Farrell and Davies, 2019; Leggett et al., 2013; Longdon et al., 2015).

As HEI does not differentiate whether documented host species are reservoirs to which the virus may be adapted, incidental hosts, or a combination of the two, it does not directly test whether heterologous inoculations caused more disease. We therefore constructed a smaller dataset of 36 viruses for which the reservoir host group was known at least to the level of taxonomic order, and introduced a binary effect (“reservoir match”) using an existing dataset of virus-reservoir associations to assign the host provenance of viruses (Mollentze and Streicker, 2020; S2.2.2). Across models, we found little compelling evidence for elevated disease outcomes in either homologous or heterologous infections (Fig. S20). A single exception was our individual-level model of disease presence, where inoculations into homologous hosts (e.g., bats inoculated with bat-reservoired viruses and rodents inoculated into rodent-reservoired viruses) increased the probability of disease (80%CI excluded zero; S20). This effect held in a sensitivity analysis that removed the HEI variable, since inoculations associated with low HEI tended to be homologous (Fig. S14; Fig. S21). Whether including or excluding HEI, models with the reservoir match effect found no effects of host order (Fig. S21, S20). Fitting these models with a reservoir match by host order interaction term similarly provided no evidence that bats and rodents differed in their ability to mitigate homologous versus heterologous infections (Figs. S22, S23 & S24).

Although a weak positive effect of reservoir match appears at odds with positive effects of HEI, it may be that for some host-virus interactions, adaptive virulence in homologous infections may outweigh maladaptive virulence in heterologous infections (Cressler et al., 2016). Alternatively, because our reservoir match metric was defined at the order and not species level, it may fail to capture the extent to which the inoculated host species experienced selection to reduce disease caused by infection with that particular virus. As a final sensitivity analysis and to separately test the hypotheses that bats have unusually potent disease suppression with either bat-associated viruses or evolutionarily novel viruses, we refit our disease outcome models to homologous and heterologous infections separately. Trends in eco-evolutionary variables and experimental design effects were similar to those observed in our main models, detecting no clear difference between bats and rodents for any measure of disease outcome (S2.2.3). This suggests that whether the virus involved is one for which bats are unlikely to have co-evolved defenses or one where virus-specific antiviral defenses may have putatively co-evolved, the frequency or severity of disease in bats cannot be demonstrated to be unusual using existing data from experimental infections.

We also detected influences of experimental design in both our individual- and species-level models. In the species-level models, all three disease outcomes were positively associated with the number of inoculated individuals (Fig. 2), pointing to the possibility that experimental infection studies are often underpowered to determine whether a specific host-virus association causes disease. In our individual-level models, larger inoculation doses unsurprisingly increased the probability of disease and led to more severe disease (Fig. 2). Inoculation doses ranged over six orders of magnitude in both groups (Figs. S4, S5), but were not significantly different between bats and rodents (Fig. S6). While high doses are often used in experimental inoculations to increase the likelihood of disease (raising separate questions about how well these experiments recapitulate natural infection), this does not explain the lack of a host order effect in our models. Similarly, as inoculations directly into the brain or cranium are associated with increased severity, and are more commonly performed in bats compared to rodents (Fig. S7) we re-fit the individual-level full model removing inoculations via direct injection into the brain or crania. These models show no host order effects, indicating that the tendency for these inoculations to be associated with bats does not impact conclusions drawn from our main models (Fig. S30). Thus, our failure to detect decreased disease in bats cannot be explained by systematic differences in dose or inoculation route between bats and rodents (Figs. 2, S4, S7, S6).

Taken together, our findings support the conclusion that specialist viruses cause more severe disease and highlight ways that experimental design choices may bias current perceptions of generalised disease suppression in prominent reservoirs of viral zoonoses. Impacts of host-virus co-evolution appear to be more complex. Overt disease may be more likely for viruses infecting evolutionarily distant hosts, perhaps reflecting maladaptive virulence (Farrell and Davies, 2019). At the same time, viruses infecting species closely related to their co-adapted reservoir hosts may be more likely to successfully infect and cause disease, in this case reflecting adaptive virulence (Cressler et al., 2016) which may be exacerbated when viruses infect susceptible hosts with little to no evolved defenses.

## Comparing disease severity among bats, rodents, and humans

In principle, inoculation of bats with disproportionately pathogenic viruses could have disguised true differences in disease severity relative to rodents. We explored this idea using human case fatality rates (CFR; Guth et al., 2022) as a comparable and broadly-available measure of pathogenicity. As expected from the generally elevated CFRs of bat-associated zoonotic viruses (Guth et al., 2022), experiments inoculating bats more often used viruses with higher CFRs compared to experiments inoculating rodents (Fig. S32A). However, this effect was not robust after controlling for virus-level hierarchical effects (Fig. S35). We also failed to find a difference in the CFRs of viruses used in bats versus rodents when restricting the analysis to heterologous inoculations or when excluding bat-associated viruses (Fig. S35). As such, if a genuine reduction of disease in bats was undetectable in our dataset due to biases in viruses selected for experiments, it would primarily reflect bats experiencing less-than-expected disease from bat-associated viruses – consistent with the expectation that evolutionary association leads to virus-specific adaptations.

**Figure 3:**
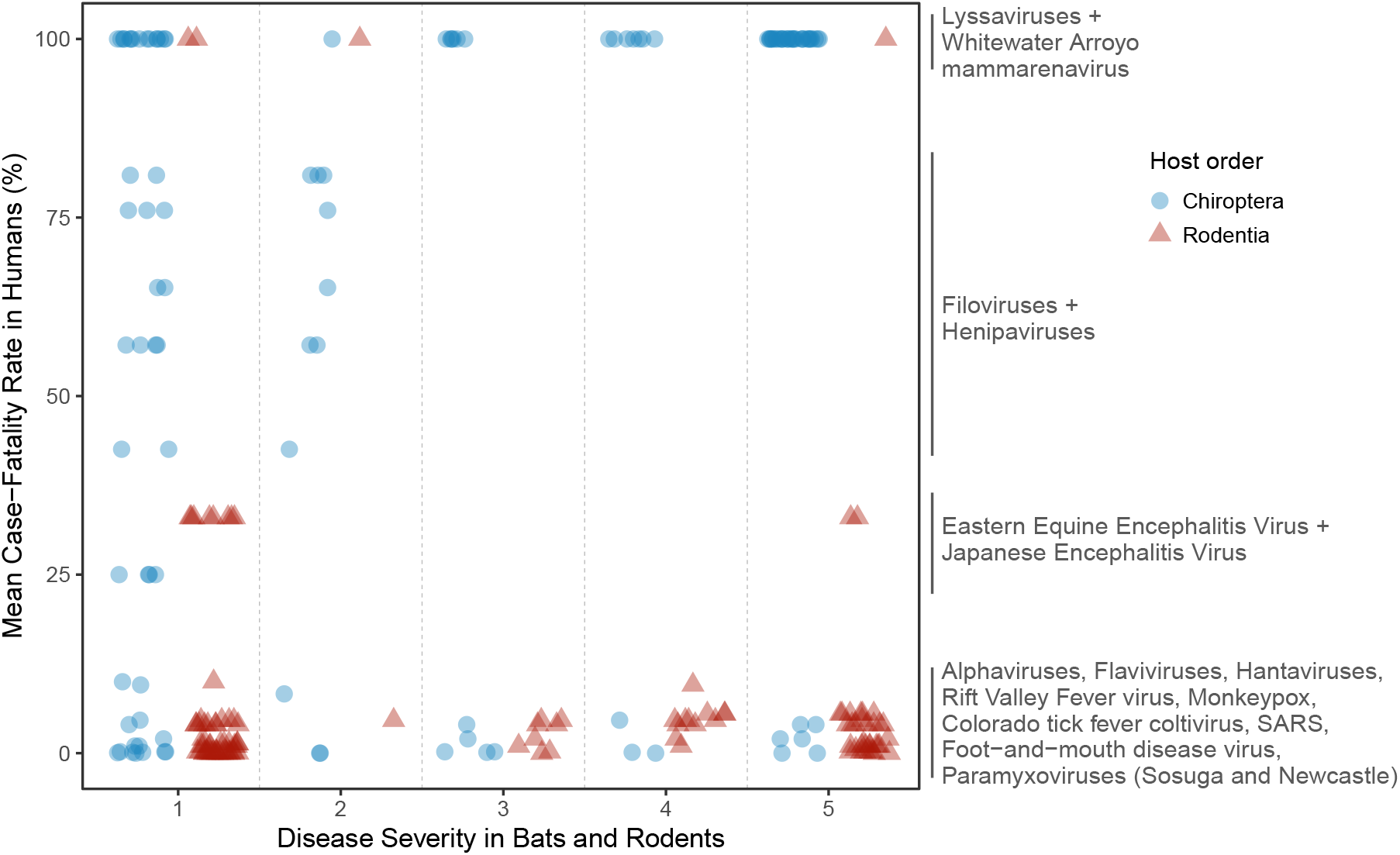
Mean human case fatality rates per virus versus maximum observed disease severity per experimental infection study. Colors and shapes indicate the taxonomic order (Chiroptera / Rodentia) of host species in the experimental infection data. See Fig. S34 for an alternate version indicating the taxonomic group of known reservoir hosts. Points are jittered horizontally to improve visibility of overlapping points.

We next related CFRs with disease outcomes in experimentally inoculated animals to examine whether bats consistently have reduced disease for viruses that cause severe disease in humans (Figs. 3 & S33). Supporting this notion, we identified a putative region where viruses cause high mortality rates in humans but mild disease in bats. However, this region included only bat-borne filoviruses (which at most caused sub-clinical disease in bats, i.e. a severity score of 2; Fig. 1) and henipaviruses (Fig. S34). More surprising, non-bat-associated viruses which cause relatively low human mortality (i.e., flaviviruses, alphaviruses, and hantaviruses; CFRs *<*20%), sometimes caused severe disease in bats, including mortality (severity scores of 5; Fig. 3) and high infection-fatality rates (Fig. S33). Disease outcomes in bats are therefore compatible with conventional expectations that vertebrate immune systems are tuned to suppress disease caused by co-evolved viruses, but that these defenses fail to reliably protect against evolutionarily novel viruses (Farrell and Davies, 2019; Poulin and Combes, 1999). In both bats and rodents, inoculations of moderate to high human CFR viruses from heterologous hosts were virtually absent (Fig. S34), highlighting a major data gap in current understanding of viral disease.

## Discussion

The belief that bats have evolved mechanisms to suppress viral disease more effectively than other animals prompted discoveries of features of bat immune systems which are proposed to have translational medical value (Baid et al., 2024; Banerjee et al., 2020; Irving et al., 2021; Ahn et al., 2023). Synthesizing eighty-six years of experimental infections, we show that a wide diversity of viruses cause severe disease in bats and find no evidence for systematic differences in disease occurrence or severity between bats and rodents, whether considering all viruses, viruses that putatively co-evolved with bats, or viruses that are presumably novel to bats. Instead, disease outcomes are shaped by factors which would be expected in any vertebrate, including aspects of experimental design, the host specificity of viruses used, and to a less certain extent the degree of co-evolutionary association with the inoculated host. These findings support the conclusion that some bat species evolved defenses which limit disease from specific co-evolved viruses, but the current literature provides no evidence that the frequency, potency, or cross-virus generalizability of these defenses in bats should be expected to be unusual relative to other mammals.

Our results, constituting the first quantitative test of whether bats experience comparatively low rates of virus-induced disease, support the idea that bats are unlikely to possess a ‘one size fits all’ solution where life-history driven selection resulted in generalized mechanisms to reduce the frequency or severity of viral disease (Pei et al., 2024). If unique traits of bats (increased longevity given body size, flight) led to the evolution of unique antiviral mechanisms, our results suggest that these are either insufficient to protect bats at the organismal level (versus effects observed in vitro) or only protect against certain viruses. For example, we report a variety of non-bat associated viruses which caused severe disease in bats and show that the co-evolutionary context of infection influences disease outcomes in bats, patterns which would not be expected if bats had near-universal protection against viruses. Equally surprising, disease outcomes remained unexceptional in bats even when considering putatively co-evolved infections. This suggests that the evolutionary pressures resulting from the unique interaction of bat life history (longevity, flight) with the viruses they maintain have not necessarily resulted in mechanisms that allow bats to survive infections by co-evolved viruses better than other host groups. We speculate that beliefs that bats have exceptional abilities to coexist with viruses without overt signs of disease became embedded due to the association of bats with viruses that are highly pathogenic in humans (Guth et al., 2022), the challenges of observing virus-induced mortality in most bat species under natural conditions due to their small body sizes and reclusive behaviors, and an extrapolation of theory founded on a small set of well-studied systems such as henipaviruses and filoviruses (Calisher et al., 2006; Baker et al., 2013; Williamson et al., 1998; Guito et al., 2021).

Synthesizing published experimental infections introduced several challenges for our analyses. While we adjusted for variation in experimental design choices such as number of inoculated individuals, dose, and inoculation route in our models of disease outcomes, we found large heterogeneity across study designs and other features of experimental design, such as the degree of prior viral passage may have altered infection outcomes. Although the degree and mechanisms of pathogen evolution following serial passage depend on many factors (Tsugawa et al., 2014; Elena et al., 2025; Peña et al., 2016), serial passage often increases virulence in passaged hosts and attenuates virulence in original hosts (Kubinak et al., 2015; Lezcano et al., 2023; Ebert, 1998). The viruses in our study were most often passaged in either primate or rodent sources, rather than bats (Fig. S11), which might be expected to decrease, not increase the severity of disease outcomes in inoculated bats. Where available, the mean number of prior passages also did not differ for bat and rodent inoculations (Fig. S12), suggesting that our inability to recover effects of host order on disease outcomes is unlikely attributable to systematic biases in passaging practices. Nevertheless, future studies should examine how passage number, host origin, and passaging mechanisms affect disease outcomes. Our study was also limited to bats and rodents. While these mammalian groups are commonly used in experimental viral infections and – as the two largest mammalian orders – host the majority of zoonotic viruses (Mollentze and Streicker, 2020), the omission of other host clades opens the possibility that both bats and rodents experience unusually mild disease. However, given that the life history strategy of rodents prioritizes reproduction over longevity (Albery and Becker, 2021; Hayman, 2019; Han et al., 2015; Luis et al., 2013), and mechanisms that limit the severity of viral infections in bats are suggested to be byproducts of adaptations to powered flight (Baker et al., 2013; Brook and Dobson, 2015; Calisher et al., 2006; O’Shea et al., 2014; Shen et al., 2010; Wang et al., 2011; Wong et al., 2007), this outcome seems unlikely. Although our results should be re-visited as inoculations into other host orders accumulate, we suggest that bats and rodents are well characterized groups that represent broader phenomena of taxon-specific immunological traits (Shaw et al., 2017) and context-dependent outcomes of host-virus interactions. It is also conceivable that certain bat clades have heightened tolerance or susceptibility to viruses, akin to results observed in experimental infections of invertebrates (Longdon et al., 2015; Imrie et al., 2024). While the hierarchical effects of bat species in our models of disease outcome do not obviously support this conclusion (S2.4), we caution that sparse data for some bat clades makes these effects difficult to interpret. Another implicit challenge of using experimental infection data is that experiments are typically carried out over short timescales that preclude quantification of long term fitness impacts, and they cannot quantify viral load in infected tissues without sacrificing individuals or using highly invasive sampling (both of which could affect host fitness). As these unmeasured outcomes define viral tolerance, both our results and outcomes of individual experiments may reflect either resistance, tolerance, or a combination of both. Irrespective of the mechanism, we find no evidence for unusual control of infection outcomes in bats. Another limitation of our analysis is that comparatively few viruses (8 out of 54) were inoculated into both rodents and bats. However, when subset to include only these viruses, our models still found no consistent difference between bats and rodents (S2.2.7), reinforcing the current limited evidence for enhanced tolerance to viral infection in either clade.

### Implications for future research

The belief that bats evolved exceptionally generalized or unusually potent mechanisms to suppress virus-induced disease largely underpins the aspiration that bats will be an unusually rich source of broad-acting or potent antivirals. The incompatibility of the currently available data with this premise questions whether the translational value of bat immunity should be expected to be disproportionately high relative to other host groups. Nevertheless, our results do not diminish the potential biomedical value of uncovering the mechanisms by which animals evade severe disease, and useful antivirals may be developed from bat immune systems in the future. Indeed bat ASC2, though in early stages of development, may represent one such biomedical application of bat immunity (Ahn et al., 2023). Further, the mechanisms that underpin putatively elevated tolerance observed in co-evolved host virus-relationships (e.g., filoviruses and henipaviruses) may yield useful therapeutics targeted to particular viral clades. However, by countering the current narrative that such biomedical breakthroughs will arise *disproportionately* from studying bats, our study points to the potential value of antiviral discovery in immune systems of other organisms.

Looking forward, it will be useful to identify how specific immunological traits protect across related viruses and, if some host taxa show heightened ability to reduce viral disease, to determine which viruses their immune features may suppress. To this end, we identify major evidence gaps in the current body of experimental inoculations: few bat-associated viruses have been inoculated into rodents, rodent-associated viruses with moderate human CFR have not been inoculated into bats or wild rodents, and relatively few human-associated viruses have been inoculated into either bats or rodents (Fig. S34). Future comparisons of disease outcomes across host groups should inoculate representative species from each group with the same sets of viruses, comprising those that are both homologous and heterologous with a paired design involving the same doses and inoculation routes. Although it would incur major logistical challenges and costs, such experiments would ideally be carried out using non-invasive proxies for viral load (e.g. using bioluminescent viruses (Stading et al., 2016)) and would observe animals over sufficient timescales to quantify multiple impacts of infection on host fitness. When known, studies focused on comparing natural immune responses should also aim to use realistic doses and inoculation routes. Our finding that inoculation dose and study size influence observed disease outcomes also suggests that some studies showing limited disease may be artifacts of insufficient powering of experiments for disease detection. This emphasizes the need for larger experiments to detect disease outcomes, and the value of synthesizing data across multiple experiments to detect biases which would be missed by single studies and to make generalizable inferences despite them, as we have done here.

The recurrent emergence of high impact zoonoses from bats, including ebolaviruses, henipaviruses, and coronaviruses, makes the need to study the relationship between bats and viruses unquestionable. The inconsistency between our results suggesting that disease outcomes in bats follows conventional evolutionary expectations, and the pervasive notion of exceptional viral tolerance in bats emphasizes the need to enhance communication between virologists, immunologists, ecologists, and evolutionary biologists. In particular, we urge greater consideration of context dependence in bat-virus interactions through using analogous baselines (e.g., comparing bats and other wildlife rather than bats to humans or laboratory animals), explicit measurement of tolerance versus resistance mechanisms within host individuals, and incorporating robust quantitative consideration of host phylogeny, virus phylogeny, and co-evolutionary associations to differentiate specific outcomes from generalizable conclusions.

## Acknowledgements

We thank members of the Streicker group, the MRC-University of Glasgow Centre for Virus Research, the University of Glasgow School of Biodiversity, One Health & Veterinary Medicine, and Nicole Mideo for valuable feedback. This work was supported by a Philip Leverhulme Prize in Biological Sciences (PLP-2020-362), a Wellcome Trust Senior Research Fellowship (217221/Z/19/Z), the NSF/BBSRC Ecology and Evolution of Infectious Diseases Program (DEB 2011069, BB/V003798/1) awarded to D.S., and an NSERC PDF awarded to M.J.F. S.T. was funded by the Wellcome Trust Integrative Infection Biology PhD Programme at the University of Glasgow (218518/Z/19/Z). Additional funding was provided by the UK Medical Research Council (MRC) through core support to the MRC-University of Glasgow Centre for Virus Research (Virus Cross Species Transmission Programme: MC UU 00034/3).

## Data & Code Accessibility

Data and scripts to reproduce analyses are available at https://github.com/maxfarrell/bat rodent inoculations and archived at Zenodo (Farrell et al., 2026).

## Author Contributions

Conceptualization: D.S. and M.J.F.; Methodology: M.J.F, N.M, and D.S.; Data compilation: M.J.F. and S. T.; Data analysis: M.J.F.; Visualization: M.J.F.; Funding acquisition: D.S. and M.J.F.; Supervision: D.S.; Writing – original draft: M.J.F.; Writing – review and editing: M.J.F., D.S., and N.M.

