## Supplementary Materials for "Viral disease outcomes are indistinguishable between experimentally infected bats and rodents"

Maxwell J. Farrell*[1,2], Samantha K. Tucker[3], Nardus Mollentze[1,2], Daniel G. Streicker[1,2]

[1]School of Biodiversity, One Health & Veterinary Medicine, University of Glasgow
[2]MRC-University of Glasgow Centre for Virus Research
[3]School of Infection & Immunity, University of Glasgow

August 19, 2026

**S3 Articles reporting disease outcomes for susceptible host species** **55**

### S1 Materials & Methods

#### S1.1 Literature identification

##### S1.1.1 Search strategy

To identify primary scientific articles describing experimental infections of viruses in bats and rodents, we first conducted a literature search via Web of Science (WOS) with Query S1 in the title, abstract, or author keywords. Terms were selected to include publications mentioning common names for groups of bats and rodents (rodent names based on the NCBI Taxonomy), viruses in general, and terms related to experimental infection. The search conducted on April 14th 2022 via the University of Glasgow returned 2,955 articles.

```
1  (   (bat OR bats OR *chiropter* OR megabat* OR microbat* OR flying fox)
2    OR (rodent OR mouse OR mice OR rat OR agouti OR anomalure OR
3        beaver OR cavy OR casiragua OR chinchilla OR chipmunk OR degu OR
4        dormouse OR gerbil OR gopher OR guiara OR guinea pig OR gundi OR
5        hamster OR hocicudo OR hutia OR jerboa OR marmot OR mole-rat OR
6        muskrat OR lemming OR nutria OR paca OR pacarana OR pectinator OR
7        porcupine OR prairie dog OR punare OR springhare OR squirrel OR
8        tree-rat OR tuco-tuco OR viscacha OR vole OR woodrat)
9  )
10 AND (virus  OR  viral  OR  "virus isolate")
11 AND ("experimental infection"  OR "experimental infections" OR
12      "experimentally infect" OR "experimentally infected" OR
13      "experimentally inoculate" OR "experimental inoculation" OR
14      "induce infection" OR "induced infection" OR "we infect" OR
15      "we infected" OR "animal challenge" OR "artificially infected")
```

Query S1: Primary Web of Science query, selecting publications involving either bats or rodents (lines 1–9), and viruses (line 10) as well as experimental infection (lines 11–15).

To supplement the WOS search results, we added publications describing virus associations with bats and rodents reported in the CLOVER database (Gibb et al., 2021), which harmonizes data on documented host species from the Global Mammal Parasite Database 2.0 (Stephens et al., 2017), the ENHanCEd Infectious Diseases Database (EID2; Wardeh et al. (2015), the Host-Parasite Phylogeny Project dataset (Olival et al., 2017), and from Shaw et al. (2020). To merge duplicated articles and limit results to those reporting experimental infections, we gathered titles and abstracts for CLOVER articles via PubMed (rentrez R package v1.2.3; Winter, 2017). We filtered the CLOVER articles to include those with the same experimental infection terms (Query S1, lines 11–15) in the title or abstract, before merging them with the WOS articles. The combined set yielded 3,495 unique articles.

##### S1.1.2 Article filtering

Because we sought to identify drivers of viral tolerance that were the natural product of ecological and life history differences among species, we limited our dataset to animals expected to be immunocompetent with a suite of immune features as indistinguishable from wild individuals as possible. Specifically, artificial selection and/or multiple generations of inbreeding may alter natural responses to infection. Artificial breeding contexts are likely to reduce exposure to pathogens and alter pressures to maintain immune responses. We therefore excluded studies conducted on individuals bought from farms or commercial breeders. We also excluded studies of common laboratory animals (mice, rats, hamster, gerbil, guinea pigs) if the origin of the animals was not indicated. For rodents from non-model species that were born in captivity, we restricted inclusion to first generation animals (i.e. offspring of wild-caught individuals). We also encountered studies that used bats which were born in captivity, but decided to retain these since bats have longer generation times than rodents, making these individuals less likely to be inbred over the time course that they have been held captive.

To remove studies on common laboratory bred animals, we searched article titles and abstracts for the same common names used in the original WOS search, but excluded the terms "mouse", "mice", "rat", and "guinea pig" to help omit common laboratory rodents. As some wild rodent common names include the terms "mouse" or "rat" (e.g., "Western jumping mouse"), we also gathered a list of Latin binomials, common names, and synonyms for all recognized bats and rodents from the Mammal Diversity Database v1.9 (Mammal Diversity Database, 2022), excluding names associated with common laboratory rodents (*Mus musculus*, *Rattus norvegicus*, *Rattus rattus*, *Cavia porcellus*, *Mesocricetus auratus*). This resulted in a list of 11,809 names used to identify articles mentioning wild bats or rodents in the title or abstract. With this approach, we identified 834 publications likely describing experimental infections in wild bats and rodents. To verify the 2661 excluded articles did not include experimental infections of wild animals, we randomly sampled 300 articles (11.27%) from this set for manual verification. Of the three hundred, we identified only one experimental infection of a wild animal, however its omission from our pipeline was due to a typographical error in the abstract of the original article, with the host referred to as a "grey quirrell" instead of "grey squirrel". This article by Timoney (1971) could not be included because we could not locate the full article text. Considering the low frequency of this kind of major typo (0.33%) – one likely caused by digitization (optical character recognition) error due to scanning of old articles – we assumed the other 2361 articles could be safely excluded.

##### S1.1.3 Identifying core publications

Of the 834 potential publications identified, we conducted a deeper manual review to exclude publications describing cell line experiments with no in vivo experiments, studies of only laboratory animals commonly available from commercial breeders (unless there was explicit mention of the animals being wild caught), studies using non-viral pathogens, and those where relevant host or virus names were mentioned for background or methodological contexts, but were not the focus of the study. We also excluded studies that did not conduct experimental infections of bats or rodents (e.g., experimental infections of non-bat and non-rodent hosts which passed earlier filters because bats or rodents were mentioned in abstracts, studies of naturally infected hosts, serological surveys, and studies of arthropod vector competence). We included experimental transmission studies (cases where a susceptible individual was infected by transmission from an infected individual in captivity), and vaccine studies if the abstract indicated there was also an experimental infection challenge, as the control groups may include useful information. At this stage, review publications were flagged for later screening of references. To avoid redundancy, datasets with individual DOIs separate from their primary paper were excluded. When primary publications for a given dataset were not returned in our search, we manually included them when relevant. The primary screen resulted in 240 core publications for inclusion.

##### S1.1.4 Expansion of core publications

We were able to gather full text articles for 238 out of 240 core publications. The remaining two articles were published in Russian in 1940 and 1941, one of which the University of Glasgow Library confirmed as a duplicate of an already included paper with slightly different title translation. A single study was excluded as the available digital copy was illegible. Of those remaining, 30 non-English articles were excluded (11 Russian, 6 French, 3 Czech, 2 German, 2 Spanish, 1 Bosnian, 1 Italian, 1 Japanese, 1 Polish, 1 Portuguese, and 1 Slovak).

Among the remaining set we identified 33 review papers. Since reviews are assumed to have conducted detailed literature searches and/or are guided by expert knowledge of specific fields, we used them to expand on the set of core publications. First, for each review we identified any cited references to experimental infections and included them in our core set, if not already present. Second, we conducted a WOS search for literature citing these review papers to identify any recent publications that met our study criteria (WOS accessed July 8th 2022 via University of Glasgow). This search returned 1,247 citations, representing 1,200 unique citing articles, 1,179 of which were available via WOS. These publications were then filtered by searching for experimental infection terms (Query S1, lines 11–15) in the Topic field, yielding 115 publications which were screened in the same manner as the core publications. Note that within WOS "Topic" includes the Title, Abstract, Author Keywords and "Keywords Plus" which is a proprietary approach from Clarivate (owners of WOS) to generate a set of keywords derived from words or phrases that frequently appear in the titles of an article's references, but do not appear in the title of the article itself. Additional review papers identified in this step were also screened to identify any cited references to experimental infections.

In total, the expansion resulted in identification of an additional 95 publications, bringing the total set to 336. After excluding review papers (n = 50), articles without readily available full texts, articles in languages other than English, symposium abstracts, theses, and datasets unlinked to a paper, our literature search yielded 246 primary research articles which were screened for disease outcome data.

###### S1.1.5 Screening of studies for disease outcome data

The primary research articles were screened to identify publications in which authors reported the hosts as susceptible to infection (by any method of testing), and reported some aspect of disease outcome (clinical disease, mortality, histopathology, behavioral change, symptoms of infection). This screen yielded 137 articles reporting disease outcomes for susceptible host-virus combinations which were read in detail to create structured data. 105 of these contained individual-level host data, while an additional 32 articles reported data only at the host species level (see Section S3 for references).

##### S1.2 Data extraction and harmonization

###### S1.2.1 Extraction of individual-level data

For each individual, the host species, virus species, virus strain, virus passage history, inoculation route, and inoculation dose were recorded. When available the sex of each individual was also recorded. To document the outcome of infection, we recorded whether each individual was deemed susceptible to infection, was reported to show signs of clinical disease, the severity of disease as described by the authors of the study, whether there was any change in body temperature associated with infection, the degree of histopathology, individual organs or tissues that showed signs of damage, whether individuals died of infection, changes in behavior, other notable symptoms, and the amount of time individuals were observed post-inoculation.

###### S1.2.2 Extraction of species-level data

For the 32 studies where individual-level data were not reported in original articles, we extracted signs of disease mentioned across the entire study population, reporting the maximum of each disease category observed (for example, if infection resulted in mortality of any individuals, we set mortality to "True" for that host-virus-study combination). For articles that provided individual-level data, we collapsed disease outcomes across individuals to identify any instances of disease, mortality, and the maximum observed severity score (see below) for species-level models described below, but retained variation in disease outcome for individual-level models.

##### S1.2.3 Harmonization of species names, known hosts, reservoirs, and host traits

Virus names were harmonized to ICTV accepted names via NCBI. Viruses without accepted ICTV names were excluded from further analyses. These included four novel viruses (TRVL 34053-1 virus, Turkmenia-1-74, Lyssavirus Eptesicus isolate, Lyssavirus Rousettus isolate), three unique reassortments of La Crosse and Snowshoe Hare viruses, one virus not identifiable to a contemporary accepted species (Korean Hemorrhagic Fever virus), and two viruses reported only at the genus level (a *Lyssavirus* and a *Poxvirus*). Host names were harmonized to the Upham mammal supertree (Upham et al. 2019) via the Mammal Diversity Database v1.9 (Mammal Diversity Database 2022). Information on virus host ranges were taken from VIRION (Carlson et al. 2022) and used to augment information on susceptible hosts from our experimental infection data. Information on the taxonomic orders of known reservoir hosts was taken from Mollentze & Streicker (2020).

##### S1.2.4 Calculation of host specificity metrics

To investigate effects of the host specificity of the viruses used in experimental inoculations, we gathered lists of the known mammalian hosts of each virus from the VIRION database (Carlson et al., 2022) including the susceptible experimentally inoculated hosts identified in our database. Host names were harmonized to match the Upham et al. (2019) mammal supertree. As a measure of phylogenetic generalism per virus, we used the standardized effect size of phylogenetic diversity ("ses.pd"). This was calculated as the sum of phylogenetic branch lengths connecting all known hosts and adjusted using a null model of expected diversity based on the observed host richness using the *pd_query* function in the R package PhyloMeasures v2.1 (Tsirogiannis and Sandel, 2016), using the "uniform" null model with 1000 replicates each sampling with equal probability among all possible tips. As a measure of phylogenetic generalism at the host-virus level, we calculated a measure of host evolutionary isolation (Farrell and Davies, 2019), which is the mean phylogenetic distance from the experimentally inoculated host species to all other documented hosts.

##### S1.2.5 Harmonization of signs of infection

We recorded whether the authors reported the presence of "clinical disease", representing a binary subjective assessment of whether individuals showed overt signs of infection. We also recorded whether or not individuals died due to infection. Individuals that were culled as part of experimental design, died from anesthesia, or died of unknown causes not attributable to infection were recorded as unknown (NA) for death due to infection. We also recorded any reported behavioral or physical changes following infection, which we grouped into 11 broad categories (Table S1).

| *Sign of infection category* | *Changes included* |
|---|---|
| Aggression | aggression, aggressive, agitation, biting, vicious, irritable, furious |
| Vocalizations | vocalization, odd vocalization, dysphasia, hissing sounds |
| Increased sensitivity | excitement, flying violently, irritability to light |
| Impaired movement | paralysis, tremor, spasms, ataxia, weakness, paresis, nystagmus, seizure, muscle weakness, trembling, coma, unable to fly, unable to move, unable to stand, altered reflexes, incoordination, hunching, staggering, head tilt |
| Lethargy | lethargy, depression, prostration, listless, sleepy, reluctance to move |
| Loss of appetite | inappetence, ate very little, refused food, dehydration |
| Weight loss | weight loss, anorexia |
| Pneumonia | pneumonia, respiratory disease |
| Eye / nose infections | rhinitis, mild conjunctivitis, nasal discharge, ocular discharge, crusty nose, nasal crust, bloody nose |
| Skin rashes / lesions | skin lesions, cutaneous lesion, hypopigmentation, tongue lesion, lip lesion, lesions near eyes, corneal lesion, epithelial hyperplasia, multiple dermal foci, lesion on gum, skin pustules, petechial rash, vesicular rash, pustular rash |
| Internal organ damage | splenitis, cardiomyocyte necrosis, haemorrhage in lungs, lesions on brain, lesions on testes, lesions on salivary gland, hepatocellular degeneration, ulceration of villus tips, multifocally fused villi, degeneration of villus cells, or any mention of damage to internal organs or tissues (e.g., components of the respiratory tract, digestive tract, central nervous system, reproductive system, kidneys, lymph nodes, etc.) |

Table S1: Eleven sign of infection categories with examples of associated behavioral or physical changes.

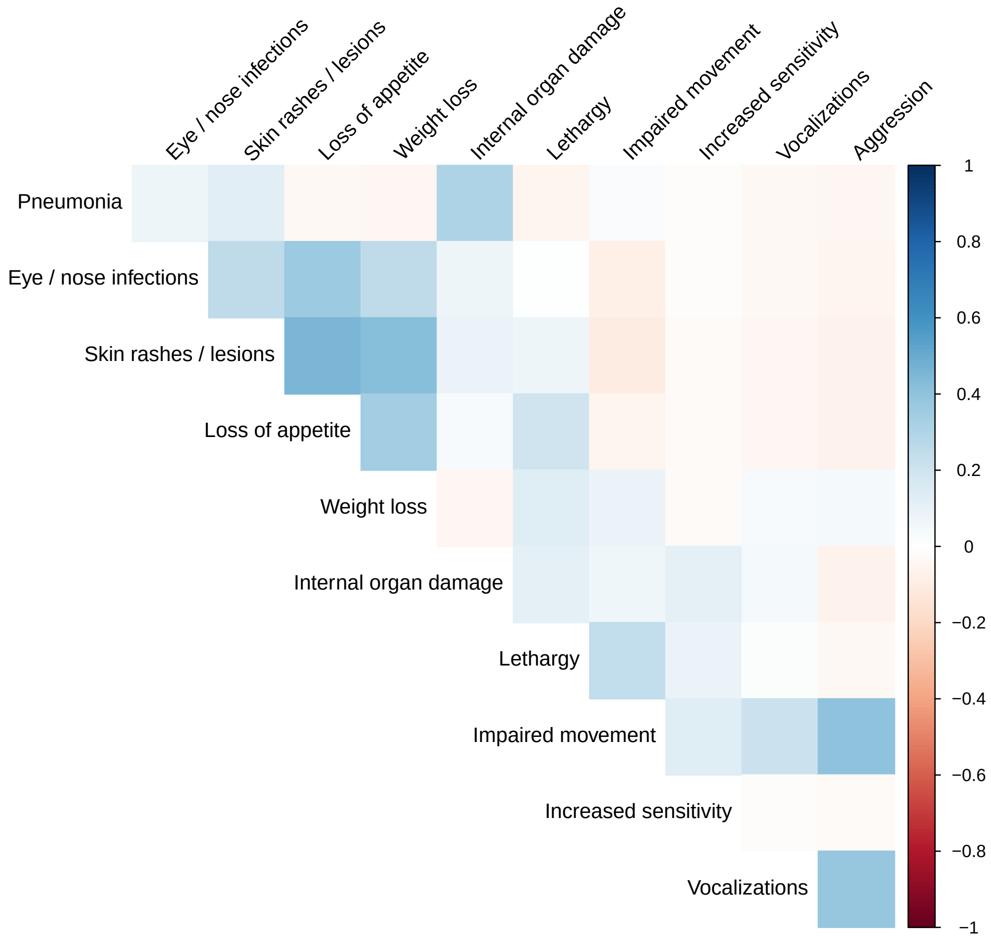

Figure S1: Pairwise correlations between 11 categories of observed signs of disease (physical and behavioral changes). Correlations are based on the presence/absence of each category per host-virus pair (e.g., if impaired movement and aggression were often observed in the same host species infected by the same virus, this is reflected in a positive correlation). Rows and columns of the correlation matrix are calculated using the angular order of the eigenvectors approach implemented in the corrplot R package (Wei and Simko, 2021).

Some studies clearly described which aspects of disease manifestation were looked for and documented. For example, Hutson et al. (2013) write "Post-inoculation, individual animals were observed daily for signs of morbidity, or malaise (inappetence, decreased activity, recumbence with reluctance to move, etc.) and clinical lesions or rash for up to 34-41 days depending on the progression of disease". However, most studies did not clearly define which signs of disease were systematically monitored. As such, signs of disease may be regarded as akin to presence-only data since we do not always have information on when clinical signs were looked for but not found. It is also important to note that by documenting reported signs of disease, we found some individuals reported as showing no "clinical disease" by the authors, but displaying one or more of the signs of disease described above. This raises the issue of subjective assessments when reporting the presence of "clinical disease". Therefore in our analyses we construct a binary response variable "presence of disease" which is equal to 1 if there is author-reported clinical disease, disease-induced mortality, or any sign of disease reported.

##### S1.2.6 Creation of disease severity measure

To investigate disease severity, we created an ordinal rank from 1 to 5 based on a combination of author-reported "clinical disease", mentions of severity, signs of disease, animal culling for ethical reasons, and death due to infection. We assign severity scores considering the potential impact of each criteria under natural conditions, assigning lower scores to outcomes that are likely to be survivable and higher scores to clinical signs (e.g., impaired movement, internal organ damage) that are likely to lead to death in natural settings. When signs of infection spanned multiple criteria (e.g., loss of appetite and death), we assigned the higher severity score.

| *Severity rank* | *Criteria* |
|---|---|
| 1 - No disease | Authors state "no clinical disease" or do not mention clinical disease, AND no other symptoms or signs of illness are reported. No mortality due to infection. |
| 2 - Subclinical disease | Authors state "no clinical disease" but report some symptoms. No mortality due to infection. |
| 3 - Mild to moderate disease | Authors report "clinical disease" but do not report specific symptoms (therefore we assume these are mild), OR authors report clinical disease or do not explicitly mention it AND report "mild", "minimal", or "mild to moderate" disease, OR report "mild", "moderate," or "medium" severity of histopathology, OR report symptoms including eye/nose infection, lethargy, skin infection, loss of appetite, weight loss, aggression, altered vocalizations, increased sensitivity, or pneumonia. No mortality due to infection. |
| 4 - Severe disease | Authors report "severe" disease, OR report "severe" histopathology, OR report evidence of internal organ damage OR report impaired movement OR report that disease was so severe that the animal was culled for ethical reasons. No mortality due to infection. |
| 5 - Death | Authors report death due to infection |

Table S2: Criteria used to assign ordinal severity scores to individuals. Ranking ranges from 1 (low) to 5 (high).

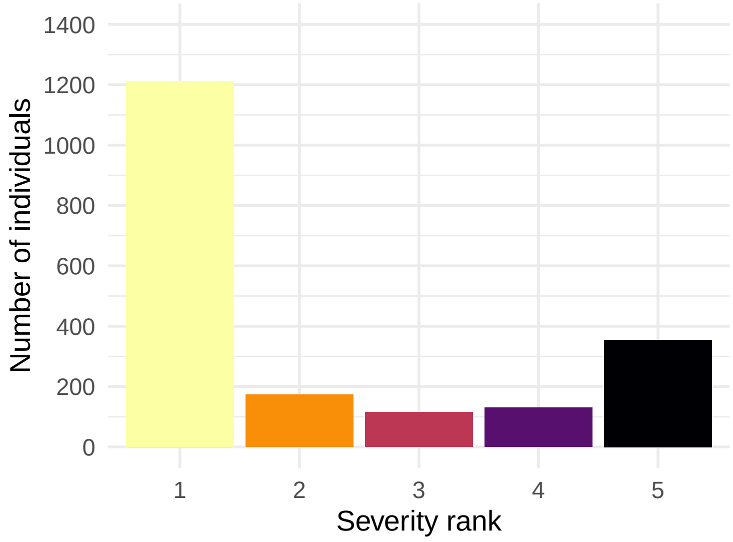

Figure S2: Distribution of ranked disease severity for susceptible individuals (n=2324). See Table S2 for details on how ranks were attributed.

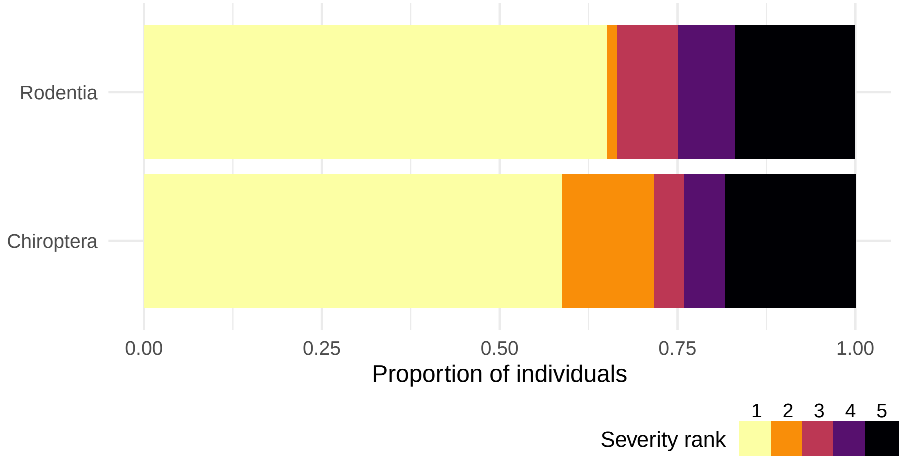

Figure S3: The proportion of individual bats and rodents assigned to each severity rank (n=2324). See Table S2 for details on how ranks were attributed.

##### S1.2.7 Harmonization of dose and inoculation route

Infectious dose and route of inoculation per individual were recorded. As dose information was reported in various units, we grouped dose units into four main categories: Mouse LD50 (predominantly Mouse Intracerebral LD50, but including various mouse ages, mean and median measures, and unreported inoculation sites), Plaque Forming Units (PFU), Tissue Culture Infectious Dose 50 (TCID50, including median, mean, and different tissue cultures as this was rarely reported), and Focal Forming Units (FFU). Mouse LD50 was most commonly reported, followed by PFU, TCID50, then FFU. For each individual, dose per body weight was calculated using interpolated mean adult body weights per species from the COMBINE mammal trait database (Soria et al., 2021). We normalize the inoculation dose by mean species body mass in order to allow for more meaningful comparisons across taxa as host body size has been hypothesized to affect viral infections through differences in metabolic rate, immune system responses, and host physiological factors such as organ size, blood volume, number of cells which influence disease progression following inoculation (Cable et al., 2007; Althaus, 2015; Banerjee et al., 2017). In a comparative study, the same number of viral particles in a given inoculation would result in a relatively lower inoculation dose for larger bodied animals – a minimum dose may lead to a productive infection in two differently sized species, but a larger dose may be needed to cause severe disease in a larger animal. As such, inoculation dose should be normalized to a host specific measure such as dose per gram of body weight. While there is debate about the exact shape of the allometric relationship to use when modelling inoculation-disease scaling (Cable et al., 2007; Althaus, 2015; Banerjee et al., 2017), we argue that use of a simple dose-per-weight metric is necessary to adjust for proportional differences in physiological capacity among host species and the viral load per unit of tissue, offering a fairer comparison of disease severity than the raw estimate of viral particles per inoculation. To determine the sensitivity of our results to this normalization approach, we fit alternative individual-level models using dose uncorrected for body mass (Fig. S28) and adding a separate body mass effect (Fig. S29), both of which yielded qualitatively similar results to our main models dose divided by average body mass.

While some studies have conducted experiments to determine the relationship between units, such as Hardy et al. (1974) finding that PFU and MICLD50 were roughly equivalent for Western equine encephalitis virus strains, exact relationships between dose units were unavailable for most viruses, making it impossible to harmonize dosages into a single comparable scale. Instead, we standardized to units of standard deviation per dose category and include a hierarchical effect of dose category in models adjusting for relative dose (see section S1.4). We also recorded inoculation routes when reported and grouped these into six major types: injection into the the brain (i.e., intracerebral and intracranial), intramuscular, intraperitoneal, under the skin (subcutaneous, subdermal, or intradermal), oronasal inoculation (including oronasal, oral, nasal, or intranasal), and simulated "natural" infections (e.g., transplacental, contact, arthropod vectors, biting, or aerosols). Dose quantities were not reported for natural infections and treated and could not be included in individual-level models adjusting for relative dose.

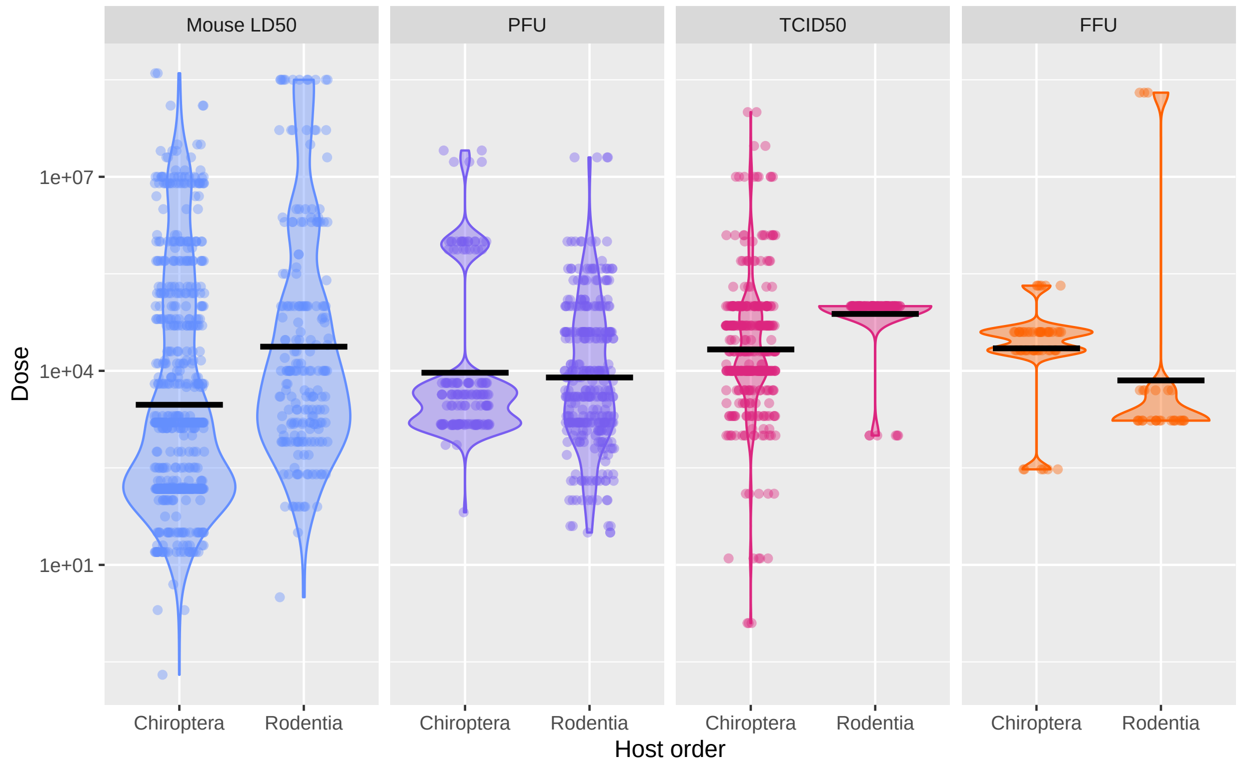

Figure S4: Reported doses per individual shows no systematic differences between bats and rodents. Dosage is adjusted for body mass using the average adult female mass per species in grams. Doses are subset to the four most commonly reported metrics: Mouse Lethal Dose 50 (LD50), Plaque Forming Units (PFU), Tissue Culture Infective Dose 50 (TCID50), and Focal Forming Units (FFU). Means per dose unit and host order are indicated by black bars.

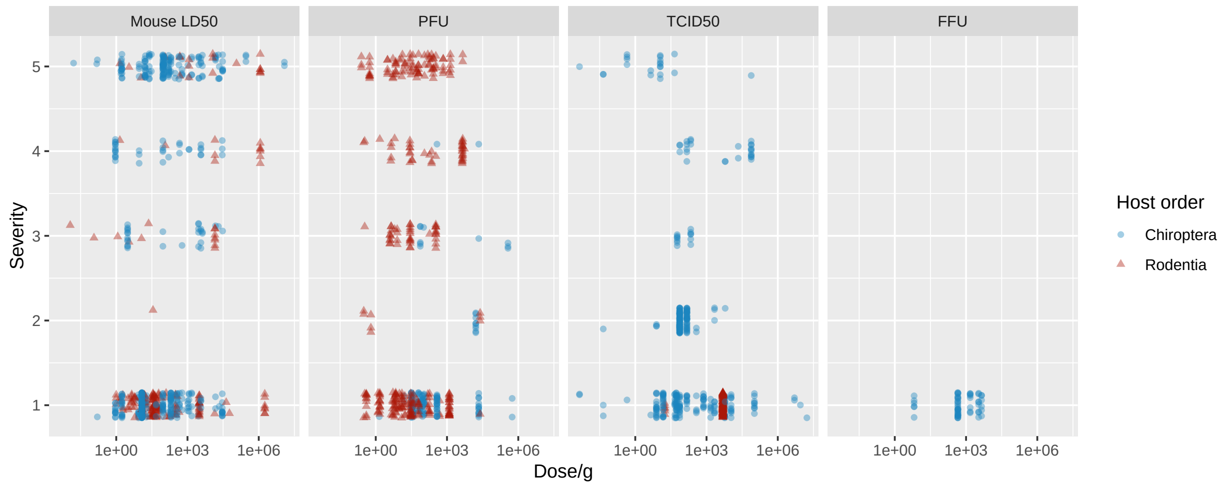

Figure S5: Severity does not systematically differ between bats and rodents across dose units. Plots show severity per dose adjusted for the average body weight per species and are faceted by the unit dose was measured in, with colors representing host taxonomic order.

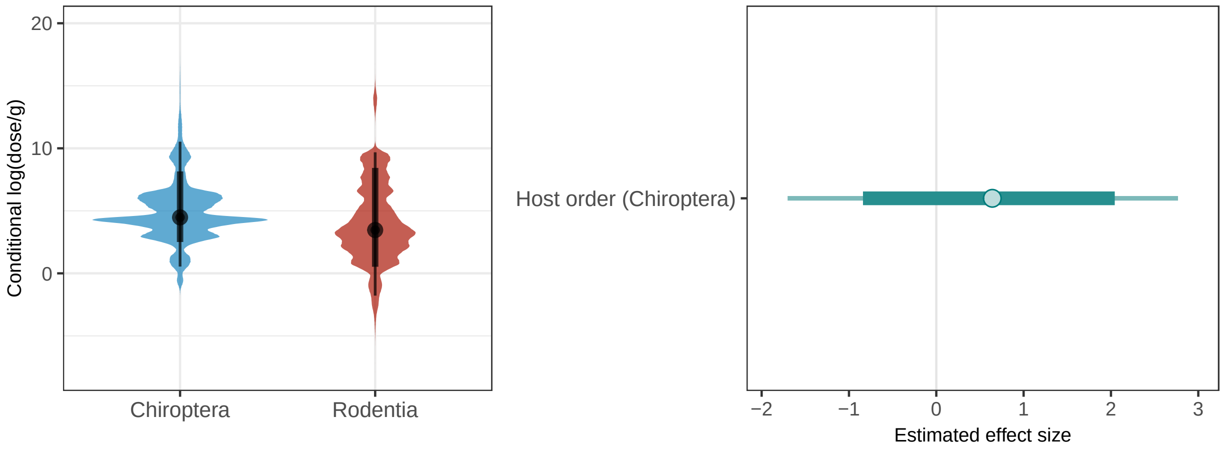

Figure S6: Absence of systematic differences in dose for experiments inoculating viruses into bats versus rodents. A) Conditional effect of host order on log-transformed dose adjusted for the average body weight per species. B) Estimated effect size of host order on log-transformed dose adjusted for the average body weight per species. Models include hierarchical effects for dose unit (Mouse LD50, PFU, etc.), inoculation route, and both phylogenetic/taxonomic and non-phylogenetic/non-taxonomic hierarchical effects for hosts and viruses.

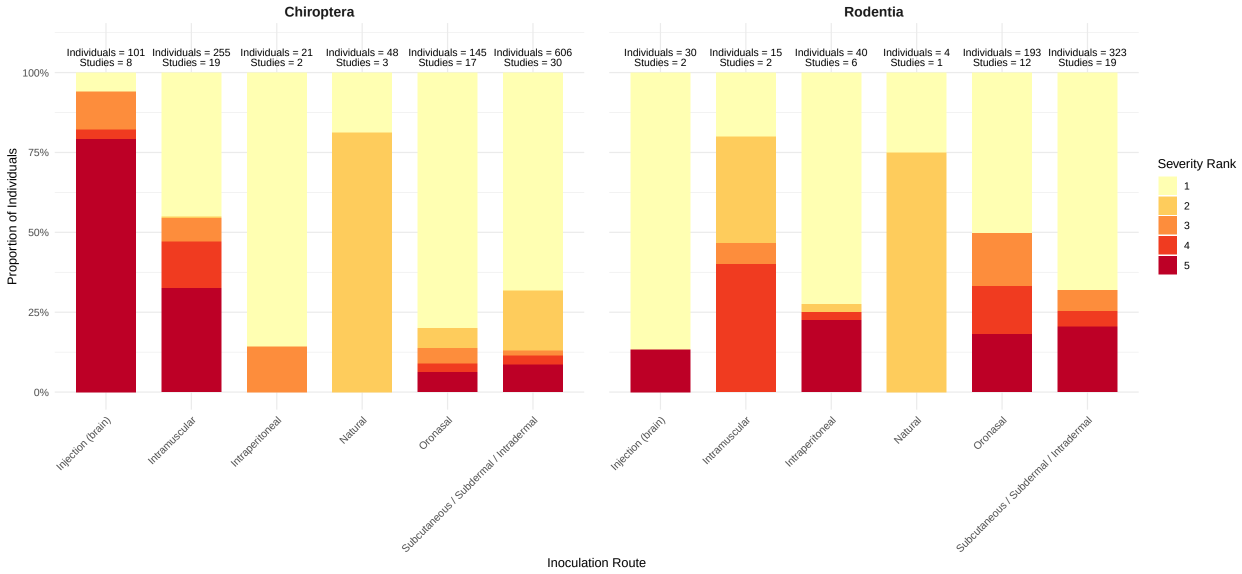

Figure S7: Severity across inoculation routes and host orders. As inoculations directly into the brain (i.e., intracerebral and intracranial injections) are often lethal and more likely to be performed on bats compared to rodents, we fit additional sensitivity models removing this inoculation route (See S2.2.6).

#### S1.3 Data descriptions and summaries

##### S1.3.1 Phylogenetic distributions of inoculated hosts

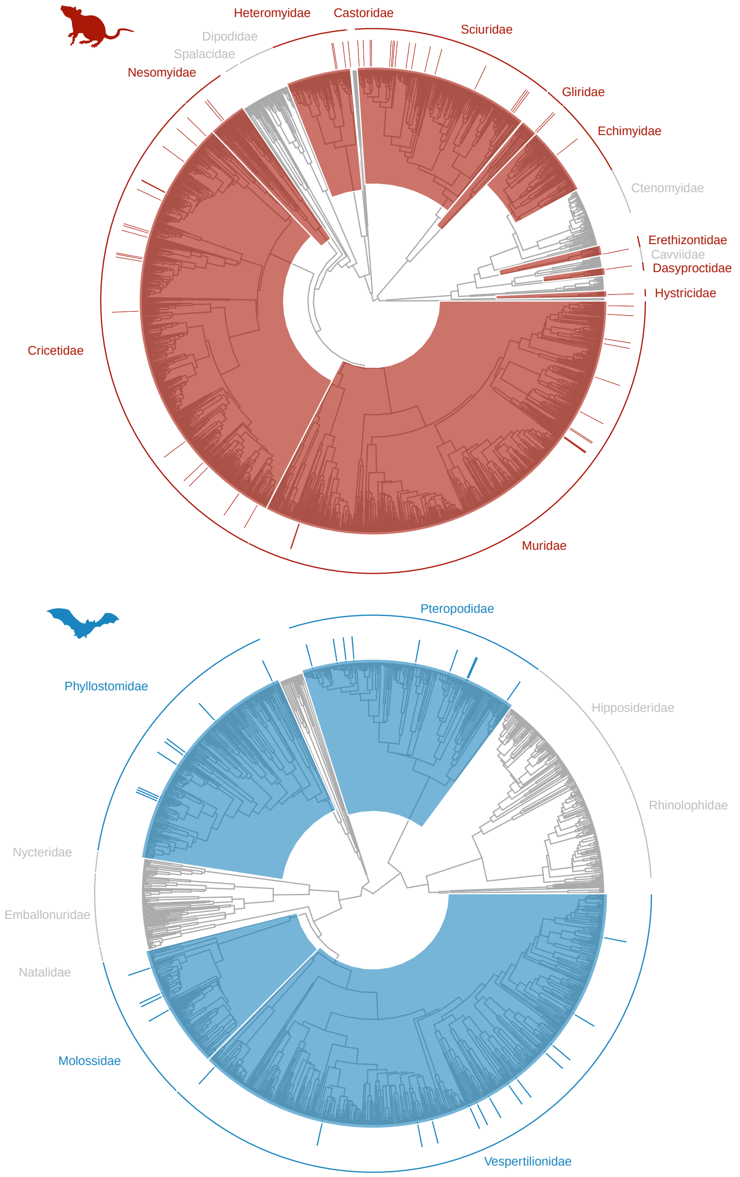

Figure S8: Phylogenetic distributions of rodent (top; n=53) and bat (bottom; n=32) host taxa included in our experimental infection dataset. Species included in experimental infections are indicated by bars. Included host families are colored according to host order; host families which were not used in experimental infections are in gray. We label unrepresented bat families with more than ten species, rodent families with more than twenty species. Among bats, the four most species-rich of the eighteen recognized families were represented: Vespertilionidae, Pteropodidae, Phyllostomidae, Molossidae. Among rodents, eleven of thirty four families were represented: Sciuridae, Cricetidae, Muridae, Heteromyidae, Nesomyidae, Castoridae, Dasyproctidae, Echimyidae, Erethizontidae, Gliridae, Hystricidae (see Table S3 for counts of species per family).

##### S1.3.2 Sampling effort across host families

| Order | Family | N hosts | N viruses | N individuals | N articles |
|---|---|---|---|---|---|
| Chiroptera | Vespertilionidae | 13 | 15 | 390 | 27 |
| Chiroptera | Pteropodidae | 9 | 29 | 461 | 29 |
| Chiroptera | Phyllostomidae | 8 | 8 | 479 | 19 |
| Chiroptera | Molossidae | 4 | 11 | 669 | 13 |
| Rodentia | Cricetidae | 18 | 16 | 234 | 17 |
| Rodentia | Sciuridae | 17 | 15 | 337 | 31 |
| Rodentia | Muridae | 12 | 11 | 196 | 10 |
| Rodentia | Heteromyidae | 3 | 2 | 38 | 2 |
| Rodentia | Gliridae | 2 | 3 | 21 | 4 |
| Rodentia | Nesomyidae | 2 | 6 | 23 | 3 |
| Rodentia | Echimyidae | 1 | 1 | 10 | 1 |
| Rodentia | Erethizontidae | 1 | 1 | 7 | 1 |
| Rodentia | Hystricidae | 1 | 1 | 7 | 1 |
| Rodentia | Dasyproctidae | 1 | 1 | 5 | 1 |
| Rodentia | Castoridae | 1 | 1 | 2 | 1 |

Table S3: The numbers of experimental host species per host order and family, including numbers of viral species tested, numbers of individuals with individual-level data reported, and numbers of articles.

##### S1.3.3 Sampling effort across viral families

| Family | N viruses | N hosts | N individuals | N articles |
|---|---|---|---|---|
| *Flaviviridae* | 14 | 29 | 895 | 28 |
| *Rhabdoviridae* | 10 | 15 | 3803 | 36 |
| *Peribunyaviridae* | 9 | 13 | 272 | 9 |
| *Togaviridae* | 8 | 39 | 471 | 12 |
| *Filoviridae* | 6 | 4 | 189 | 7 |
| *Paramyxoviridae* | 4 | 5 | 140 | 10 |
| *Hantaviridae* | 4 | 6 | 155 | 5 |
| *Arenaviridae* | 2 | 3 | 54 | 3 |
| *Phenuiviridae* | 2 | 15 | 467 | 6 |
| *Coronaviridae* | 2 | 4 | 48 | 4 |
| *Poxviridae* | 1 | 4 | 132 | 11 |
| *Spinareoviridae* | 1 | 5 | 20 | 2 |
| *Picornaviridae* | 1 | 2 | 25 | 2 |
| *Orthomyxoviridae* | 1 | 4 | 136 | 4 |
| *Orthoherpesviridae* | 1 | 1 | 18 | 1 |
| *Hepadnaviridae* | 1 | 2 | 104 | 4 |

Table S4: The numbers of experimental virus species, host species tested, individuals with individual-level data reported, and papers per viral family.

##### S1.3.4 Accumulation plots for hosts, viruses, and host-virus pairs

To investigate potential changes in study bias we plotted the accumulation of unique hosts, viruses, and host-virus interactions included in our experimental dataset through time, separated by host order (Fig. S9). The plots indicate that a greater diversity of rodent-virus pairs are included in experimental inoculations, however this is driven by the inclusion of a larger diversity of rodents, while bat-virus pairs comprise an increasing number of viruses inoculated into a smaller number of bat hosts.

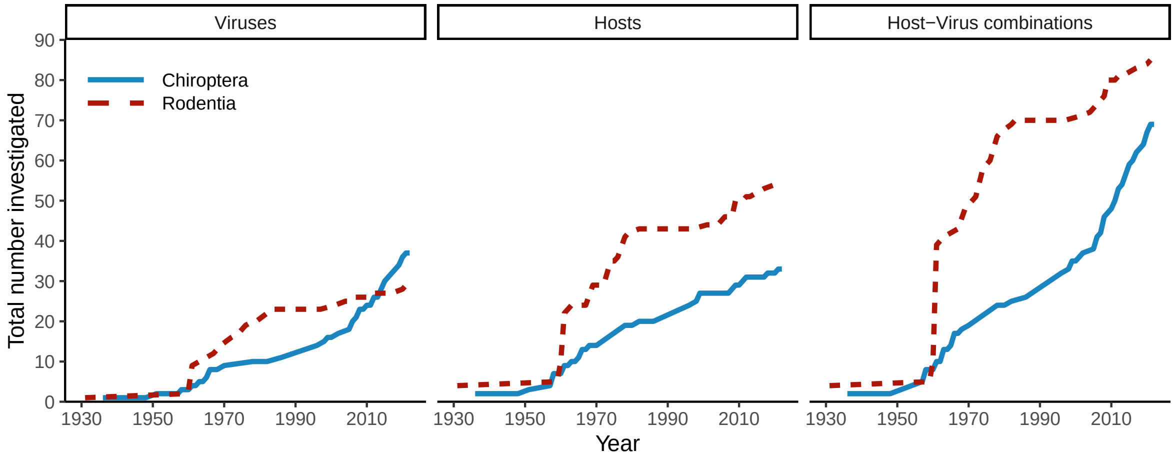

Figure S9: Accumulation curves for the total numbers of unique virus species, host species, and host-virus combinations included in our dataset of experimental inoculations through time. Colors and line types represent host taxonomic order.

| Virus family | Any disease | | Clinical | | Sub-clinical | |
|---|---|---|---|---|---|---|
| | Bats | Rodents | Bats | Rodents | Bats | Rodents |
| *Arenaviridae* | 2 / 2 | 0 / 1 | 2 / 2 | 0 / 1 | 0 / 2 | 0 / 1 |
| *Coronaviridae* | 0 / 1 | 1 / 1 | 0 / 1 | 1 / 1 | 0 / 1 | 0 / 1 |
| *Filoviridae* | 5 / 7 | 0 / 0 | 0 / 7 | 0 / 0 | 5 / 7 | 0 / 0 |
| *Flaviviridae* | 7 / 14 | 6 / 13 | 3 / 14 | 5 / 13 | 1 / 14 | 0 / 13 |
| *Hantaviridae* | 0 / 0 | 1 / 2 | 0 / 0 | 0 / 2 | 0 / 0 | 0 / 2 |
| *Hepadnaviridae* | 0 / 0 | 1 / 1 | 0 / 0 | 1 / 1 | 0 / 0 | 0 / 1 |
| *Orthomyxoviridae* | 3 / 3 | 0 / 0 | 2 / 3 | 0 / 0 | 1 / 3 | 0 / 0 |
| *Paramyxoviridae* | 5 / 6 | 0 / 0 | 2 / 6 | 0 / 0 | 3 / 6 | 0 / 0 |
| *Peribunyaviridae* | 1 / 1 | 1 / 9 | 1 / 1 | 1 / 9 | 0 / 1 | 0 / 9 |
| *Phenuiviridae* | 0 / 2 | 9 / 13 | 0 / 2 | 9 / 13 | 0 / 2 | 0 / 13 |
| *Picornaviridae* | 1 / 1 | 1 / 1 | 1 / 1 | 1 / 1 | 0 / 1 | 0 / 1 |
| *Poxviridae* | 0 / 0 | 4 / 4 | 0 / 0 | 4 / 4 | 0 / 0 | 0 / 4 |
| *Rhabdoviridae* | 22 / 23 | 1 / 1 | 22 / 23 | 0 / 1 | 0 / 23 | 0 / 1 |
| *Spinareoviridae* | 0 / 0 | 1 / 5 | 0 / 0 | 0 / 5 | 0 / 0 | 0 / 5 |
| *Togaviridae* | 2 / 6 | 14 / 32 | 2 / 6 | 10 / 32 | 0 / 6 | 0 / 32 |

Table S5: Host-virus pairs showing any sign of disease, author reported “clinical disease”, and only sub-clinical disease (maximum severity rank of 2: authors state “no clinical disease” but report some disease signs). For each virus family–host order combination, the denominator indicates the number of virus species–host species pairs with available data, while the numerator indicates those displaying the signs of disease at the given level. Note that clinical disease, whether present or absent, was not reported in all studies.

##### S1.3.5 Distributions of host traits for represented taxa

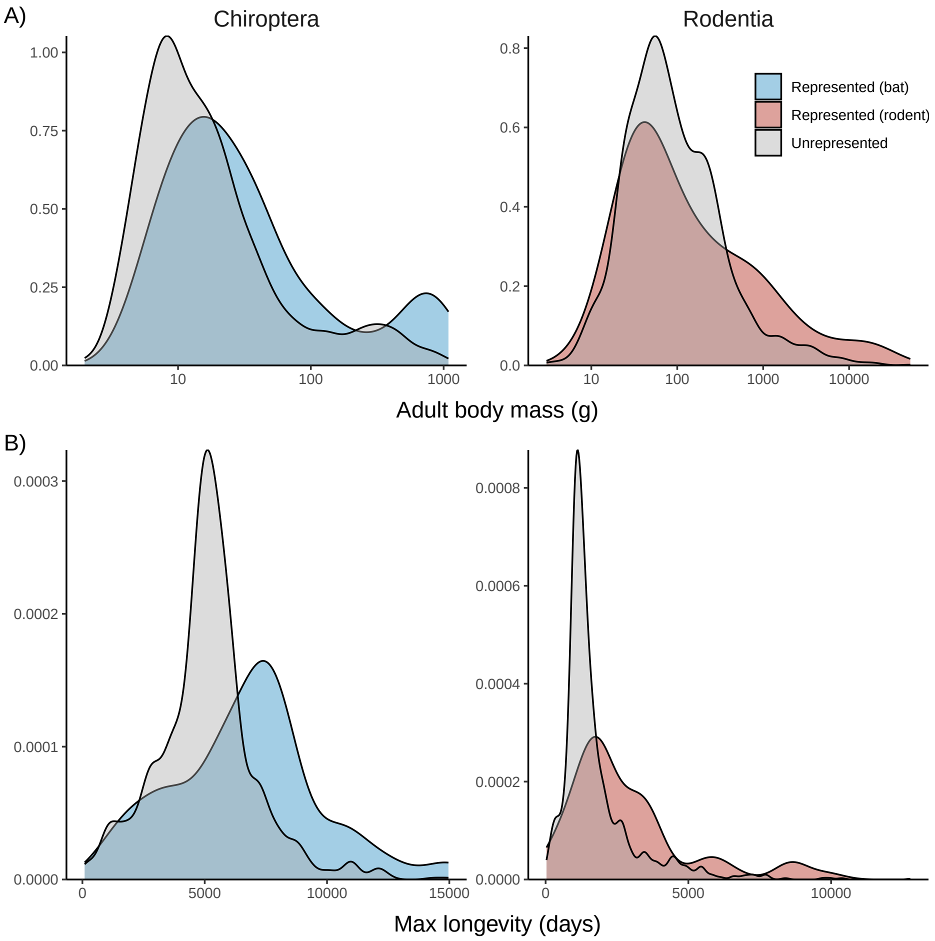

Figure S10: Distributions of life-history traits: A) adult body mass (g), and B) maximum longevity (days) for bats and rodents. Colours indicate distributions for species represented and unrepresented in our dataset. For both bats and rodents, inoculated species in our dataset spanned the full distribution of life history traits, though for both orders, there was a tendency to use relatively long-lived species in experiments. Trait data from Soria et al. (2021).

##### S1.3.6 Degree and type of viral passage prior to inoculation

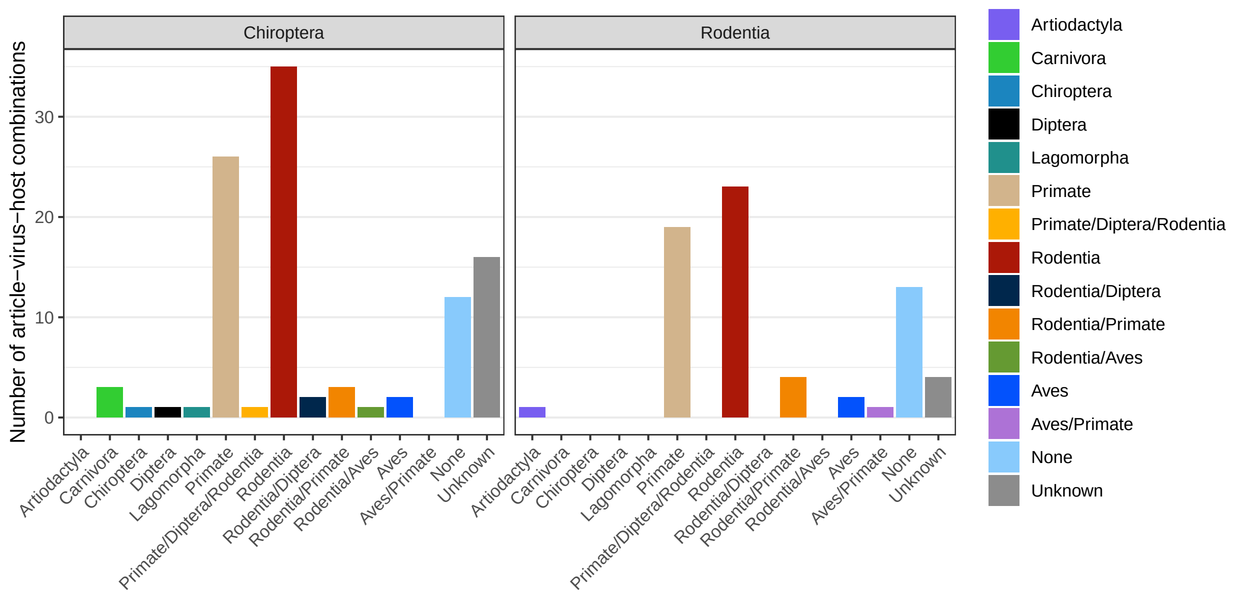

Figure S11: Distributions of host taxonomic orders for animals and cell lines used when passaging viruses prior to inoculation. Counts are numbers of article-host-virus combinations, split by the order of the inoculated host species. Viruses used in our dataset were most often passaged via either rodent or primate sources prior to inoculation, with only one study using a bat source to passage viruses (rabies virus).

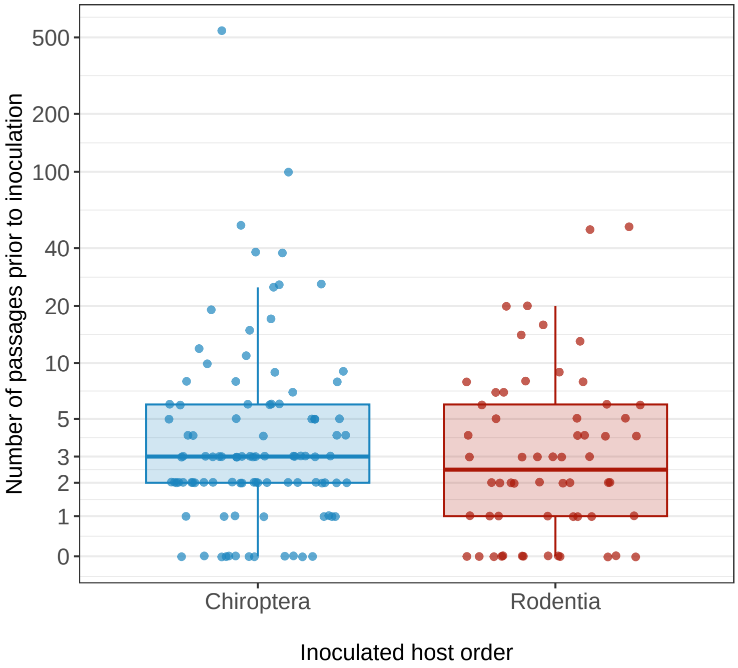

Figure S12: Numbers of viral passages prior to inoculation by the taxonomic order of the inoculated host. Each point represents a unique host-virus-article-passage number, with some articles including virus strains passaged differing numbers of times prior to inoculation. For studies where a range of passages were reported the highest number was used. For studies with passages reported in multiple host animals or cell lines the total number of passages was used. Studies for which the numbers of passages was not reported, or given a qualitative value (e.g. "low", or "many") were excluded. The mean number of viral passages prior to inoculation is similar among bat and rodent inoculations (bats: mean = 11.8, median = 3 range = 0–538; rodents: mean = 5.48, median = 2.5, range = 0–52; two-sided t-test p=0.264, t=1.12, df=109).

##### S1.3.7 Heatmap of mean severity per host-virus combination

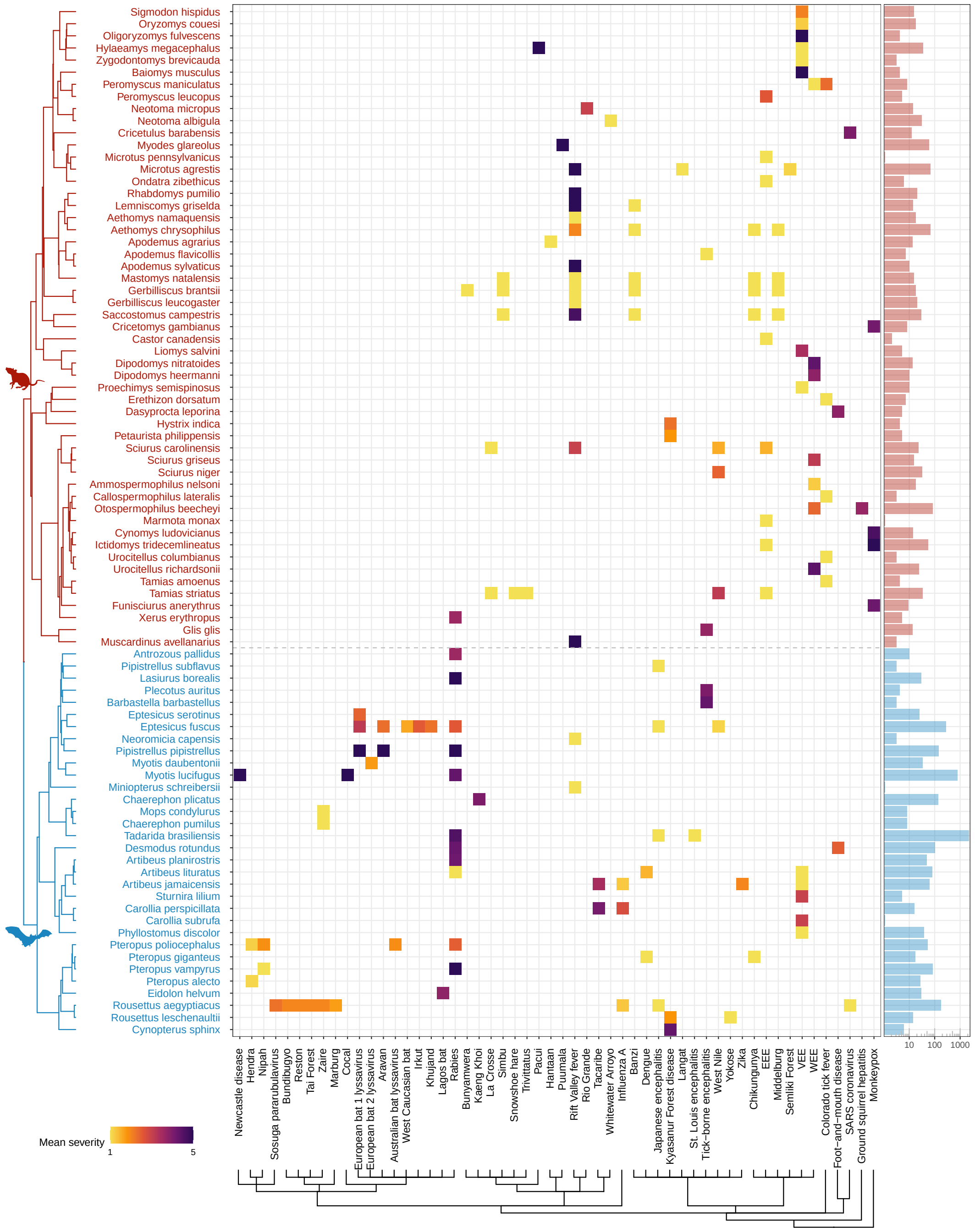

Figure S13: As an alternative to Fig. 1 we plot mean disease severity observed in experimental infections for each host-virus combination. The severity scale is calculated based on author reports of infection-induced disease and death (see S3 and S1.2.6 for details). Severity ranges from 1 (no disease) to 5 (mortality). Blank spaces indicate untested interactions. Hosts (rows) are grouped according to the Upham et al. (2019) mammal supertree. Viruses (columns) are grouped following the ICTV taxonomy. Virus names displayed are ICTV species names. If virus names ended with “ virus” this was removed to make the labels easier to read. The barchart on the right shows the total number of host individuals per species included in experimental infections.

#### S1.4 Statistical analyses

##### S1.4.1 General modelling procedure

Hierarchical Bayesian regressions were used to model three disease responses: presence/absence of any detectable disease (severity score $>1$; Bernoulli response with logit link), mortality (Bernoulli response with logit link), and disease severity (ordinal response with a cumulative probit link). To account for phylogenetic non-independence and species-level effects, all models included hierarchical effects of host and virus species, separated into phylogenetic (assuming a Brownian motion model of evolution) and non-phylogenetic effects, following the additive quantitative genetic model adapted to interspecific data as in Schmidt et al. (2021) and later recommended by Cinar et al. (2022) as best practice for meta-analyses in ecology and evolution. The correlation matrix for determining host species-level Brownian phylogenetic effects was calculated from the Upham et al. (2019) mammal supertree. Because a single phylogeny for all viruses in the study is not available, we instead used a taxonomic tree representation for viruses, based on ICTV taxonomy as reported by NCBI on February 23, 2024, with branch lengths of unit 1 per taxonomic level, subsetting the taxonomy to include only Superkingdom, Realm, Kingdom, Phylum, Class, Order, Family, Genus, Species, following Mollentze and Streicker (2020). For species-level and individual-level models we additionally include a hierarchical effect of article to adjust for potential variation in study design unaccounted for by continuous predictors (e.g., numbers of individuals, doses and inoculation routes).

To aid in the comparison of effect sizes across continuous and binary predictors, all continuous predictors were centered and scaled to a standard deviation of 0.5 (Gelman, 2008), with number of individuals (species-level models) and dose/mass (individual-level models) log-transformed prior to scaling and centering. Models were fit in Stan version 2.21.0 (Stan Development Team, 2023) using the R package *brms* version 2.16.3 (Bürkner, 2017). All models used Normal(0, 1.5) priors for regression coefficients and Normal(0, 1) priors for hierarchical variance parameters, as recommended by McElreath (2020). Full models were run across four independent chains, with a minimum of 4000 iterations per chain. For each chain, the first 50% of iterations were used as burn-in and discarded, resulting in a minimum of 8000 post-warmup posterior draws per model. We diagnosed convergence by observation of Rhat values equal to 1.0 for all estimated parameters, and ensured models fit well using posterior predictive checks.

##### S1.4.2 Dose and inoculation route

As the reported inoculation doses varied substantially across and within studies (Fig. S4), we included an effect of dose divided by the average body mass of the infected species in individual-level models. To include doses measured through different dose units (e.g., Mouse LD50, TCID50, PFU), dose/g was log-transformed, then standardized and centered relative to the mean and standard deviation per dose unit. Models including dose included this continuous dose effect as well as a hierarchical effect of dose unit. In addition to dose, studies varied the inoculation routes used during experimental infection (Fig. S7). Since some inoculation routes, such as intracerebral infection, might lead to more severe disease compared to others, such as oronasal exposure, inoculation route was included as a hierarchical effect. In all models, dose/g showed a positive effect, indicating that increasing dose relative to body mass led to a higher probability of showing signs of disease, mortality, and disease severity.

##### S1.4.3 Eco-evolutionary metrics

To test whether virus host specificity and the evolutionary position of inoculated hosts influence the presence and severity of disease we include three eco-ecoevolutionary predictors. Hypothesizing that costs of generalism (Leggett et al., 2013; Antonovics et al., 2013) leads specialist viruses to cause more severe disease, we included a measure of host phylogenetic diversity (more specifically the standardized effect size of phylogenetic distances among hosts – *ses.pd*; section S1.2.4), with larger values indicating viruses with greater phylogenetic host ranges. Across all models host phylogenetic diversity showed a mean negative effect.

We also wanted to test the hypothesis that viruses inoculated into host species that are distantly related to their co-adapted hosts may cause more severe disease (Farrell and Davies, 2019). One measure of potential host adaptation is whether the inoculated host is related to known reservoir hosts. However, the reservoir hosts of many viruses included in our dataset were unknown (Babayan et al., 2018). We therefore used two proxy variables. The first, host evolutionary isolation (Farrell and Davies, 2019), is calculated as the mean phylogenetic distance from all documented hosts to the inoculated host. As pathogens commonly infect closely related host species (Antonovics et al., 2013; Davies and Pedersen, 2008; Park et al., 2018), this metric assumes that the phylogenetic centroid of known susceptible species reflects the most likely position of hosts to which a virus is best adapted, and that the distance from an inoculated host to this centroid provides a reasonable proxy for the relative extent of co-adaptation (Farrell and Davies, 2019). As this metric provides a continuous proxy of adaptation and can be calculated for all viruses in our dataset, we included it in all models. While estimated effect sizes showed high uncertainty with 95% credible intervals crossing zero, the positive effect of host evolutionary isolation is consistent with previous models of infection-induced mortality for domesticated-animal pathogens (Farrell and Davies, 2019), indicating that inoculating hosts distantly related to other documented hosts may result in overt disease and more severe disease, potentially reflecting maladaptive virulence (Farrell and Davies, 2019; Leggett et al., 2013; Longdon et al., 2015).

As host evolutionary isolation does not take into account information on the relative roles of host species in transmission (e.g., reservoir versus spillover host), which may reflect a different type of host adaptation, we fit a set of sensitivity models adding a binary effect of "reservoir match". Using the taxonomic orders of documented reservoirs from Mollentze and Streicker (2020) available for 36 viruses in our dataset, we constructed a binary metric which is "True" if the inoculated host belongs to any of the known reservoir orders, and "False" if not. As this metric was not available for all viruses in our dataset (Babayan et al., 2018), and captures a different aspect of host adaptation, we explored it's effect through sensitivity models with it included as an additional predictor, and as a replacement for host evolutionary isolation (S2.2.2). The binary "reservoir match" also allowed us to run a further set of sensitivity models restricting our analysis to heterologous inoculations (i.e., viruses inoculated into species from non-reservoir groups) and homologous inoculations (i.e., viruses inoculated into species from presumed reservoir groups; S2.2.3). These analyses offer an opportunity to evaluate whether bats' capacity to tolerate viruses extends to viruses for which they are unlikely to have evolved defenses. We find that while homologous infections include smaller distances of host evolutionary isolation, there is considerable overlap in host evolutionary isolation for homologous and heterologous inoculations (Fig. S14). Further, as some viruses have presumed reservoirs in multiple host orders, plotting host evolutionary isolation across the presumed reservoir groups shows that inoculations associated with low host evolutionary isolation interactions tend to be homologous infections, but that some homologous infections include interactions marked by high evolutionary isolation.

Previous research found that increasing host evolutionary isolation is associated with increased mortality among domesticated animal pathogens, but this comes at the cost of decreased transmission (Farrell and Davies, 2019), suggesting that similar trends in our data may represent maladaptive virulence (Leggett et al., 2013; Longdon et al., 2015). Sensitivity models including a binary reservoir match effect (with or without host evolutionary isolation) showed some positive effects of reservoir match (S2.2.2). While this may be unexpected if long-term co-evolution leads to increased disease tolerance, these results may instead reflect adaptive virulence (Cressler et al., 2016) in which some level of virulence is necessary for transmission and maintenance in nature. Alternatively, as our metric of reservoir match does not capture whether the exact host species being tested is a known reservoir (see S1.4.3, Fig. S15), positive effects on disease outcomes may be explained by viruses being well adapted to the physiology of the hosts in the same taxonomic order, but not to the specific host species used in experimental inoculations, which may have experienced little to no selection to tolerate disease caused by these viruses.

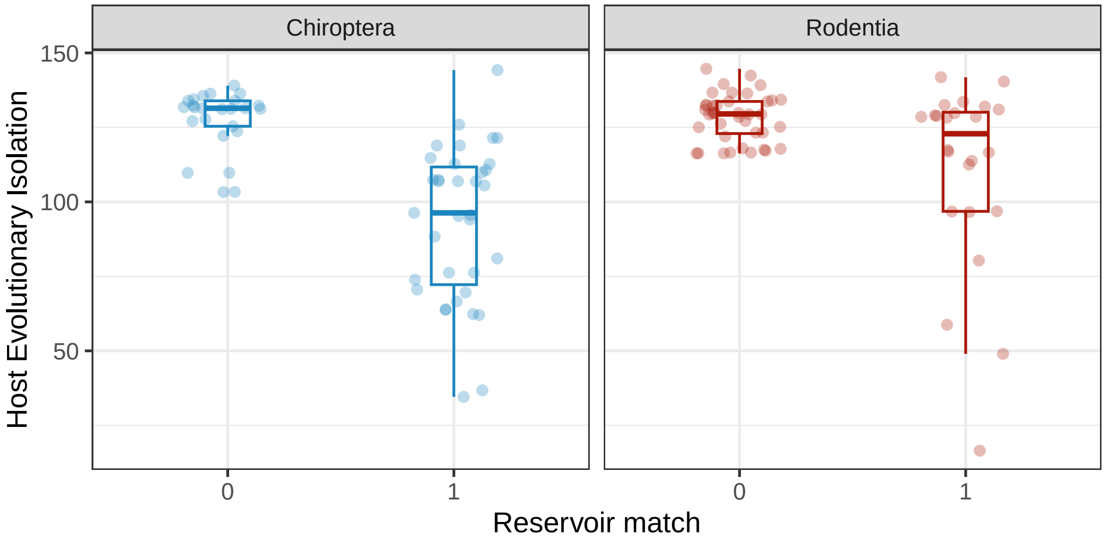

Figure S14: Distributions of host evolutionary isolation per host-virus interaction, separated by the order of inoculated host, and whether or not the inoculation was into a species belonging to the same order as a presumed reservoir species (i.e., a homologous infection).

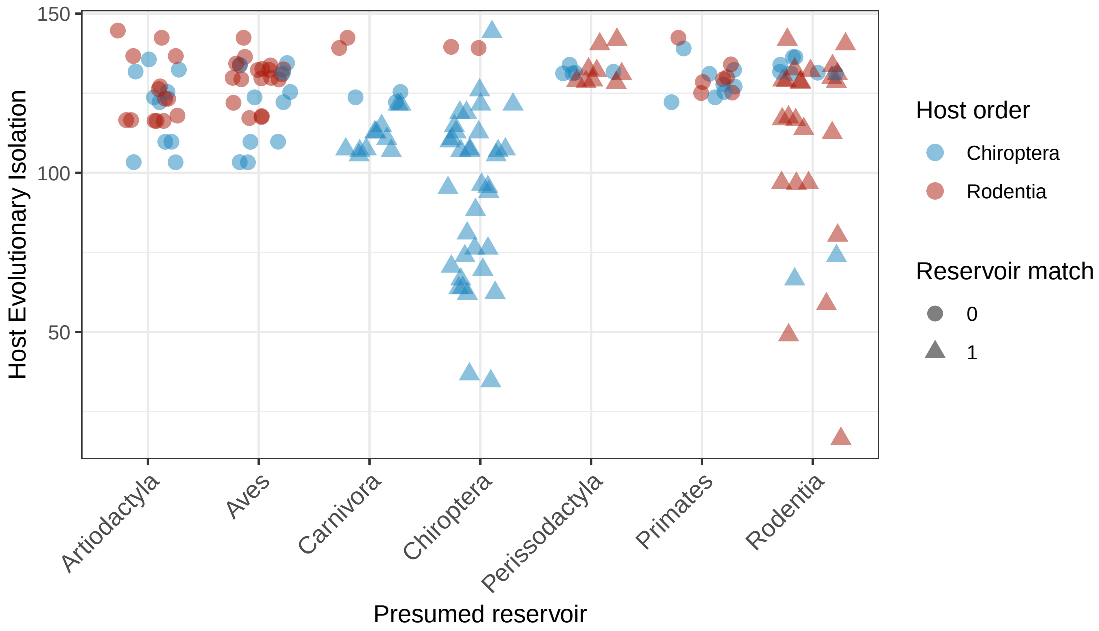

Figure S15: Distributions of host evolutionary isolation by presumed reservoir group. As some viruses are associated with multiple reservoir groups, each point represents a unique host-virus-reservoir group. Colours indicate the order of the inoculated host. Shapes indicate whether the inoculated host belonged to at least one of the presumed reservoir groups.

### S2 Supplementary Results

#### S2.1 Estimated hierarchical effects for full models

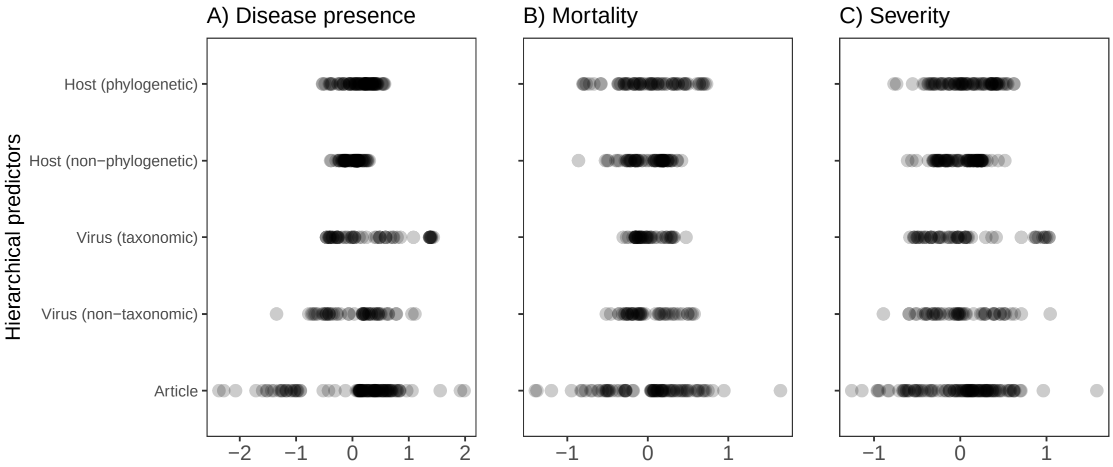

Figure S16: Estimated hierarchical effects for species-level full models (Fig. 2) of A) disease presence (n=193), B) mortality (n=154), and C) disease severity (n=193). Points represent mean estimated intercept per level of hierarchical predictor. Points are drawn at 30% transparency to help visualise overlap.

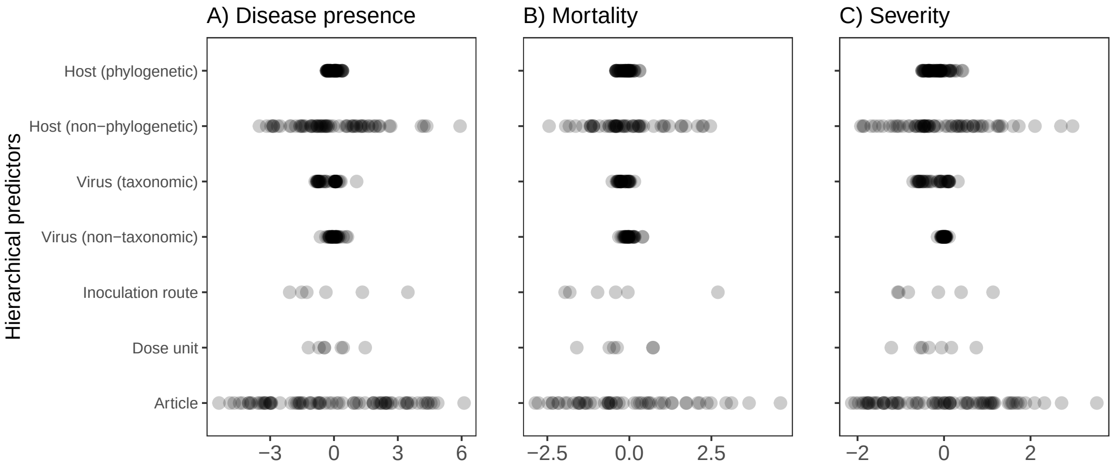

Figure S17: Estimated hierarchical effects for individual-level full models (Fig. 2) of A) disease presence (n=1900), B) mortality (n=1134), and C) disease severity (n=1999). Points represent mean estimated intercept per level of hierarchical predictor. Points are drawn at 30% transparency to help visualise overlap.

#### S2.2 Sensitivity models

##### S2.2.1 Models with only host order

Because covariates related to experimental design and reservoir group were not available for all individuals, disease presence was first modeled with a lone binary effect of host clade (Chiroperta / Rodentia), including hierarchical effects for host and virus species. Across models, there was no clear effect of host order as the effect of host order had 80% credible intervals crossing zero. To adjust for study-level variation, species-level models included an additional hierarchical effect of study ID, and the number of individuals included in the study as a continuous predictor.

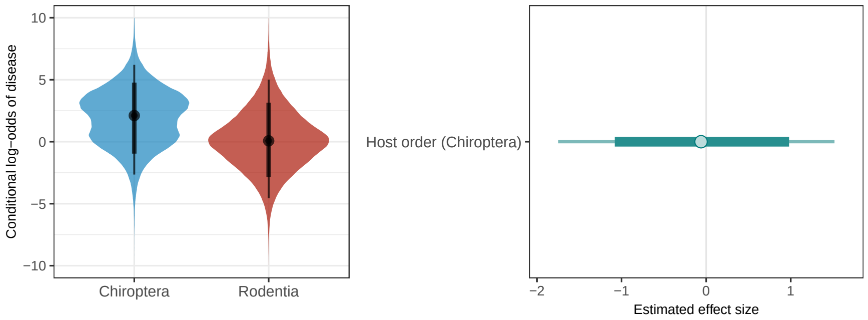

Figure S18: The species-level base model including the effect of host order and hierarchical effects for host and virus species showed no effect of host order on disease presence (N = 193). Points represent mean estimates, thick bars represent 80% credible intervals, and thin bars represent 95% credible intervals.

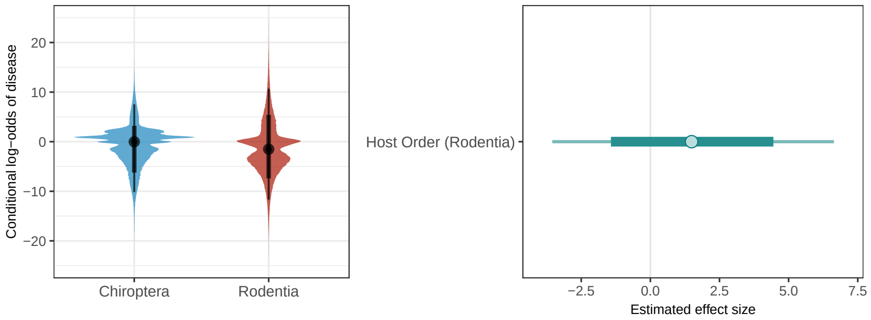

Figure S19: The individual-level base model including the effect of host order and hierarchical effects for hosts and viruses showed no effect of host order on disease presence (N = 1674). Points represent mean estimates, thick bars represent 80% credible intervals, and thin bars represent 95% credible intervals.

##### S2.2.2 Models including a binary reservoir match effect

Our main models used host evolutionary isolation as a proxy for the degree of host-virus co-evolution (S1.4.3). Using a smaller dataset of viruses for which reservoir hosts were known at least to the order level, we investigated the effect of viral "reservoir match" - a binary factor indicating whether the inoculated host belonged to the same order as one of the presumed reservoir hosts as reported by Mollentze and Streicker (2020). For bats, 58.3% (35/60) of experimental inoculations used viruses with bat reservoirs, while significantly fewer – 37.1% (23/62) – of rodent-virus interactions involved viruses with rodent reservoirs (chi-square test: p=0.019). Sensitivity analyses including reservoir match as an additional predictor showed that reservoir match had no clear effect on disease outcomes in the species level models (Fig. S20). However, in the individual level models, disease outcomes were potentially worse for reservoir-matched infections (i.e., bat viruses inoculated into bats and rodent viruses inoculated into rodents), with all three models showing mean estimated effects intervals above zero (although 80% credible intervals included zero).

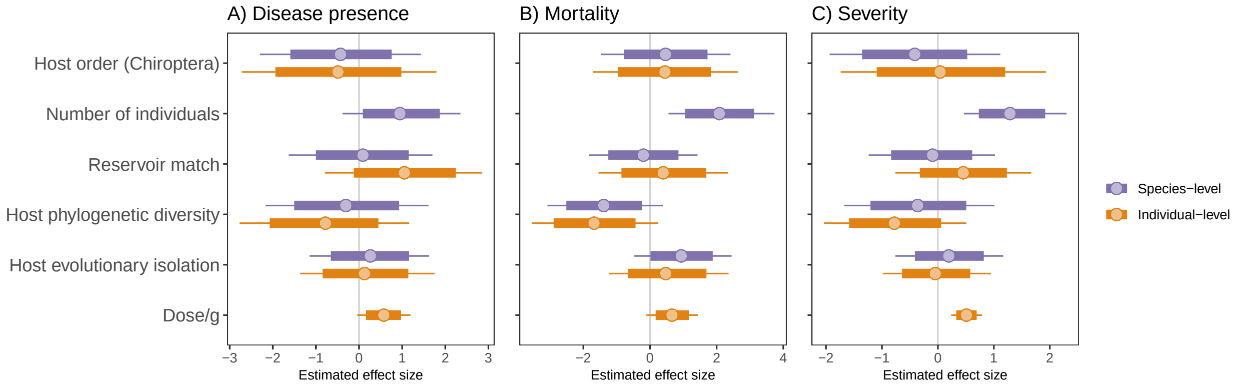

Figure S20: Estimated effect sizes for predictors for species-level and individual-level models of A) disease presence (species-level n=176; individual-level n=1920), B) mortality (species-level n=139; individual-level n=1088), and C) disease severity (species-level n=176; individual-level n=1920). Points represent mean estimates, thick bars represent 80% credible intervals, thin bars represent 95% credible intervals. Models are comparable to Fig. 2 but add a binary metric of reservoir order match.

To investigate whether reservoir match effects are influenced by the inclusion of host evolutionary isolation, we fit condensed versions of our reservoir match models (Fig. S20) removing host evolutionary isolation (Fig. S21). These models estimated reservoir matching as having similar effects to the full models with host evolutionary isolation (Fig. S20). This indicates that the estimated effects of reservoir match in these models is unlikely to be caused by confounding effects of including host evolutionary isolation, suggesting that while correlated, these metrics may capture different aspects of host-adaptation (see S1.4.3).

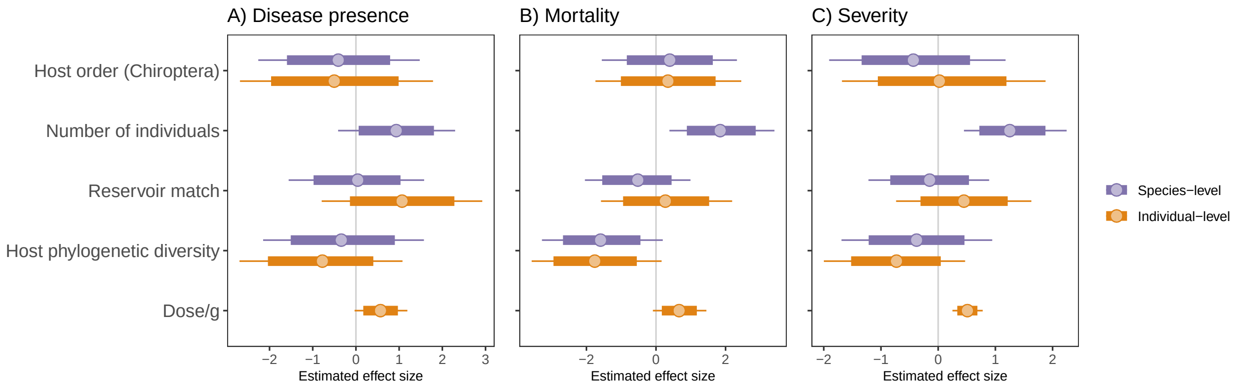

Figure S21: Estimated effect sizes for predictors for species-level and individual-level models of A) disease presence (species-level n=176; individual-level n=1920), B) mortality (species-level n=139; individual-level n=1088), and C) disease severity (species-level n=176; individual-level n=1920). Points represent mean estimates, thick bars represent 80% credible intervals, thin bars represent 95% credible intervals. Models are comparable to Fig. 2 but add a binary metric of reservoir order match and remove the host evolutionary isolation effect.

To investigate whether reservoir match effects vary by host order, we fit expanded versions of our individual-level reservoir match models (Fig. S20) adding an interaction between reservoir match and host order (Fig. S22). These models estimated reservoir matching as having similar effects to models without the interaction (Fig. S20). Plotting conditional effects of the interaction (Fig. S23 & S24) indicates that despite high uncertainties, inoculations into homologous hosts (reservoir match = 1) tend to result in more observable and more severe disease, but there is no clear interaction with host order.

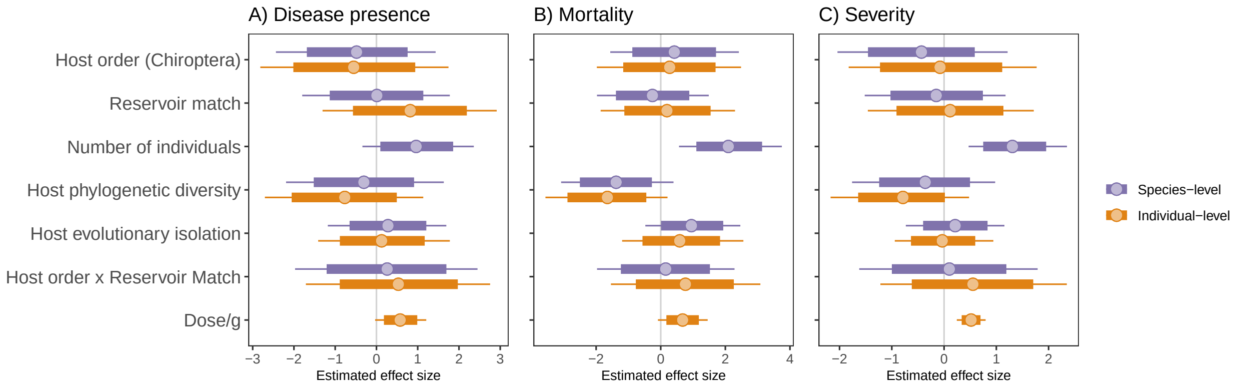

Figure S22: Estimated effect sizes for predictors for species-level and individual-level models of A) disease presence (species-level n=176; individual-level n=1920), B) mortality (species-level n=139; individual-level n=1088), and C) disease severity (species-level n=176; individual-level n=1920). Points represent mean estimates, thick bars represent 80% credible intervals, thin bars represent 95% credible intervals. Models are comparable to Fig. S20 but add an interaction between reservoir match and host order.

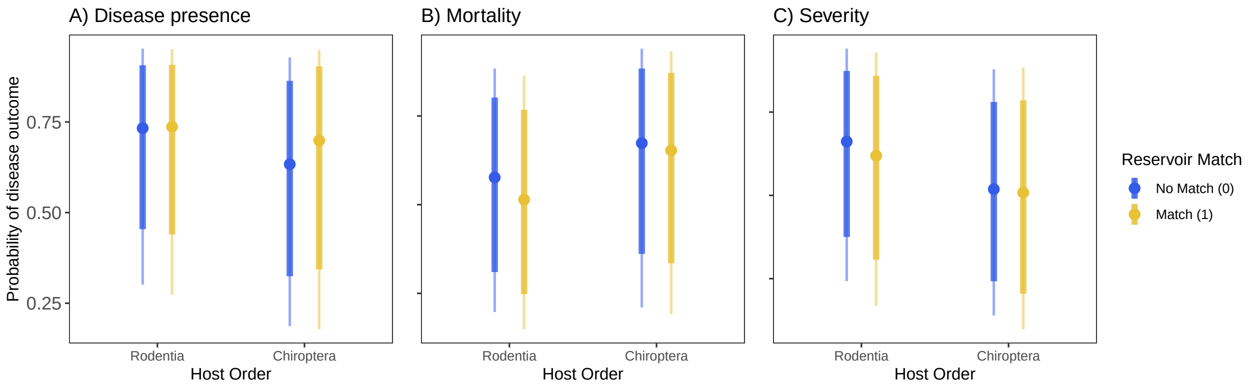

Figure S23: Estimated conditional effects of the reservoir match by host order interaction for species-level models of A) disease presence (n=176), B) mortality (n=139), and C) disease severity (n=176). Points represent mean estimates, thick bars represent 80% credible intervals, thin bars represent 95% credible intervals. Models are comparable to Fig. S20 but add an interaction between reservoir match and host order.

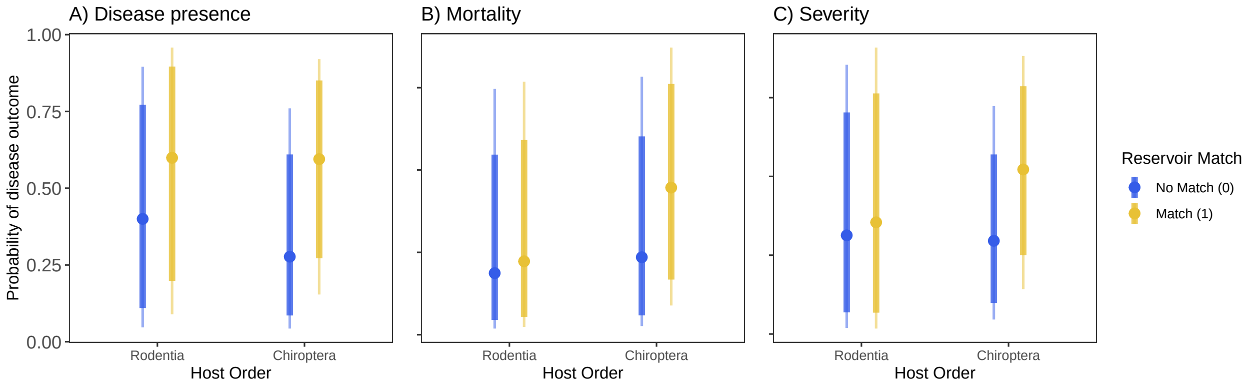

Figure S24: Estimated conditional effects of the reservoir match by host order interaction for individual-level models of A) disease presence (n=2007), B) mortality (n=1152), and C) disease severity (n=2007). Points represent mean estimates, thick bars represent 80% credible intervals, thin bars represent 95% credible intervals. Models are comparable to Fig. S20 but add an interaction between reservoir match and host order.

##### S2.2.3 Models restricted to homologous and heterologous hosts

###### Models restricted to homologous hosts

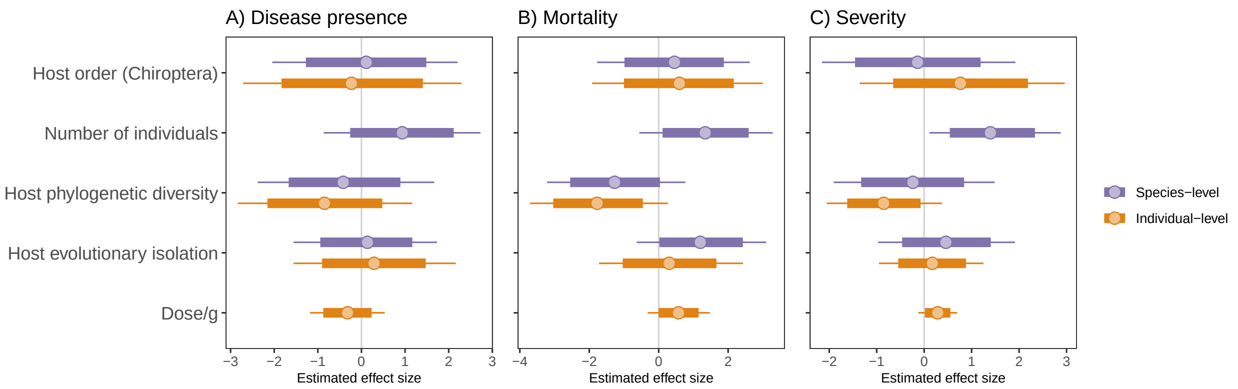

Figure S25: Estimated effect sizes for predictors for species-level and individual-level models of A) disease presence (species-level n=95; individual-level n=1114), B) mortality (species-level n=78; individual-level n=773), and C) disease severity (species-level n=95; individual-level n=1114). Points represent mean estimates, thick bars represent 80% credible intervals, thin bars represent 95% credible intervals. Models are comparable to Fig. 2 but restricted to homologous inoculations.

###### Models restricted to heterologous hosts

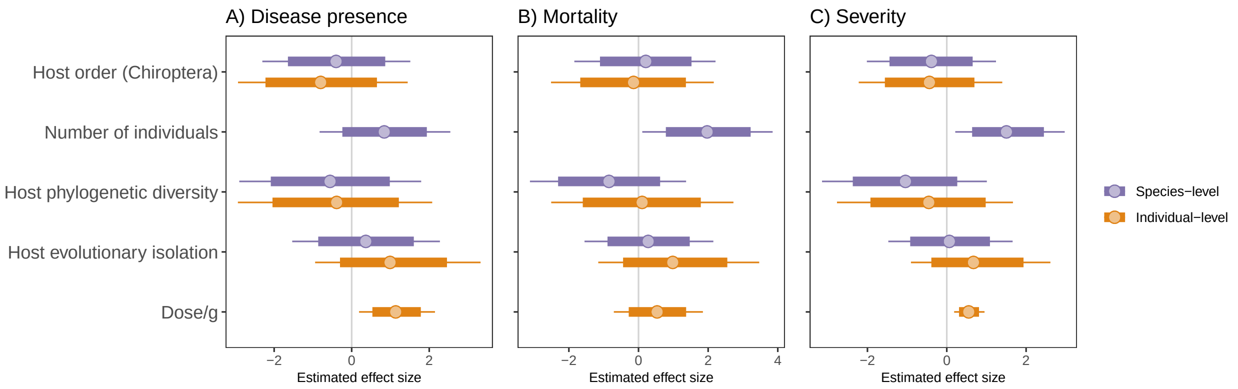

Figure S26: Estimated effect sizes for predictors for species-level and individual-level models of A) disease presence (species-level n=81; individual-level n=806), B) mortality (species-level n=61; individual-level n=315), and C) disease severity (species-level n=81; individual-level n=806). Points represent mean estimates, thick bars represent 80% credible intervals, thin bars represent 95% credible intervals. Models are comparable to Fig. 2 but restricted to heterologous inoculations.

##### S2.2.4 Models removing lyssaviruses

As lyssaviruses are known to cause mortality in their bat hosts, they are sometimes considered the "exception to the rule" regarding viral tolerance in bats (Hayman, 2019). To explore whether the lack of difference among host order in the full models may be due to the inclusion of lyssaviruses, we re-fit the full models excluding lyssaviruses (Fig. S27). We find that absence of host order effects in the full models was robust to removing lyssaviruses, which represented 23% (45/193) of host-virus-study combinations in our dataset.

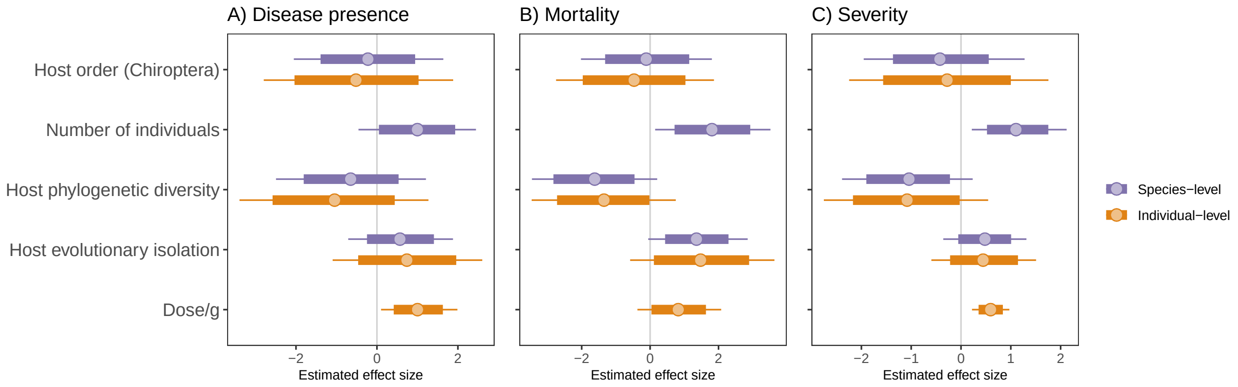

Figure S27: Estimated effect sizes for predictors for species-level and individual-level models of A) disease presence (species-level n=148; individual-level n=1308), B) mortality (species-level n=113; individual-level n=648), and C) disease severity (species-level n=148; individual-level n=1308). Points represent mean estimates, thick bars represent 80% credible intervals, thin bars represent 95% credible intervals. Models are comparable to Fig. 2 but exclude lyssaviruses.

##### S2.2.5 Models with alternative dose adjustments

To determine the sensitivity of our results to the choice of normalizing dose per average adult body mass per species, we fit alternative individual-level models swapping dose/g for the dose uncorrected for body mass (Fig. S28), and adding a separate body mass effect (Fig. S29). Both approaches to adjust for inoculation dose yielded qualitatively similar results to our main models using dose divided by average body mass.

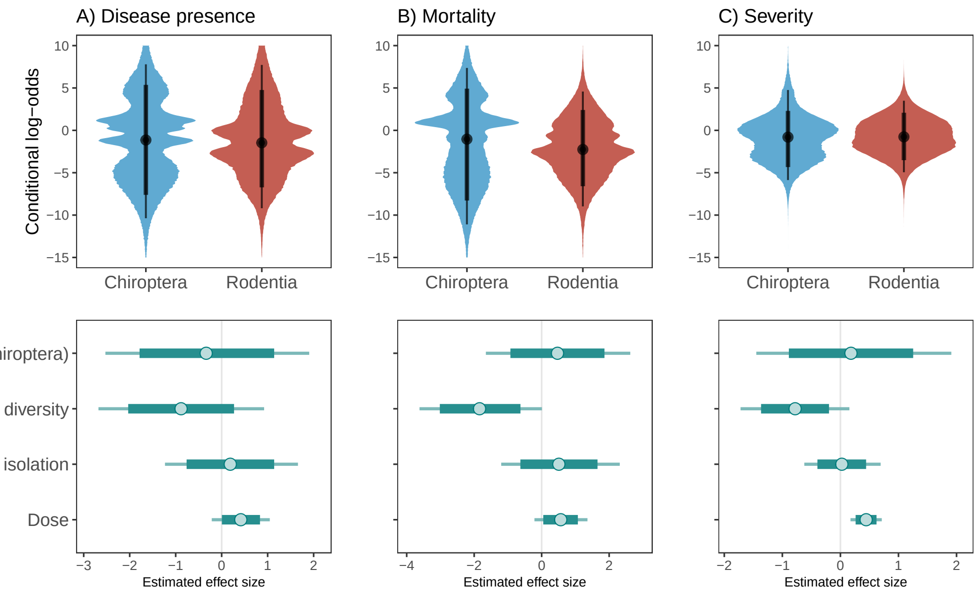

Figure S28: Conditional effect of host order (top row) and estimated effect sizes for predictors (bottom row) for species-level models of A) disease presence (n=1999), B) mortality (n=1134), and C) disease severity (n=1999). Points represent mean estimates, thick bars represent 80% credible intervals, thin bars represent 95% credible intervals. Models are comparable to individual-level main models (Fig. 2) but using the reported dose uncorrected by average body mass per species.

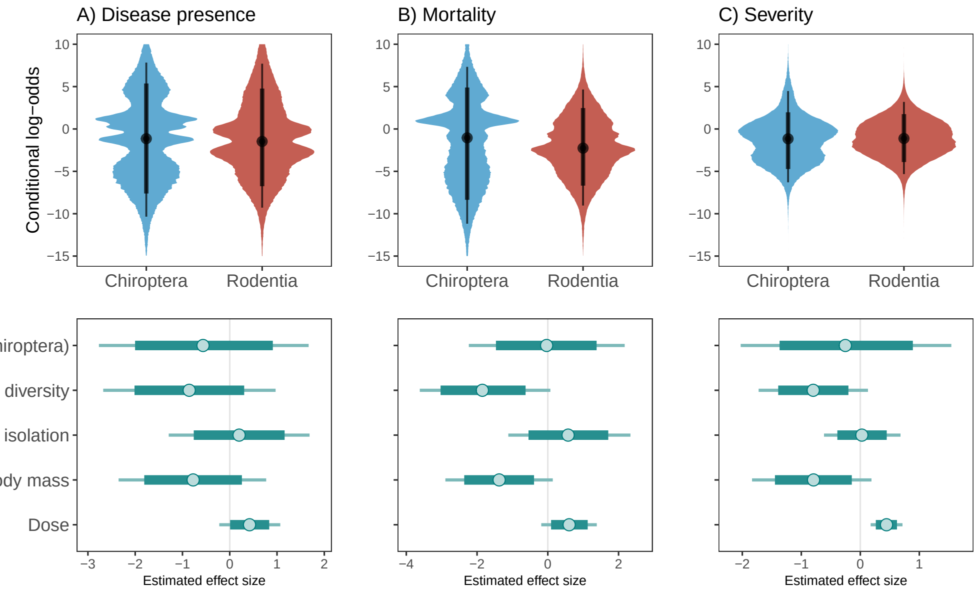

Figure S29: Conditional effect of host order (top row) and estimated effect sizes for predictors (bottom row) for individual-level models of A) disease presence (n=1999), B) mortality (n=1134), and C) disease severity (n=1999). Points represent mean estimates, thick bars represent 80% credible intervals, thin bars represent 95% credible intervals. Models are comparable to individual-level main models (Fig. 2) but using the reported dose uncorrected by average body mass per species and including a separate effect of average body mass per species.

##### S2.2.6 Models excluding inoculations into the brain

As inoculations directly into the brain or cranium are associated with increased severity, and are more commonly performed in bats compared to rodents (Fig. S7), we re-fit the individual level full model removing inoculations via direct injection into the brain or crania (Fig. S30). The results show host order effects consistent with main individual-level models including all inoculation routes (Fig. 2) indicating that the tendency for these inoculations to be associated with bats does not impact conclusions drawn from our main models.

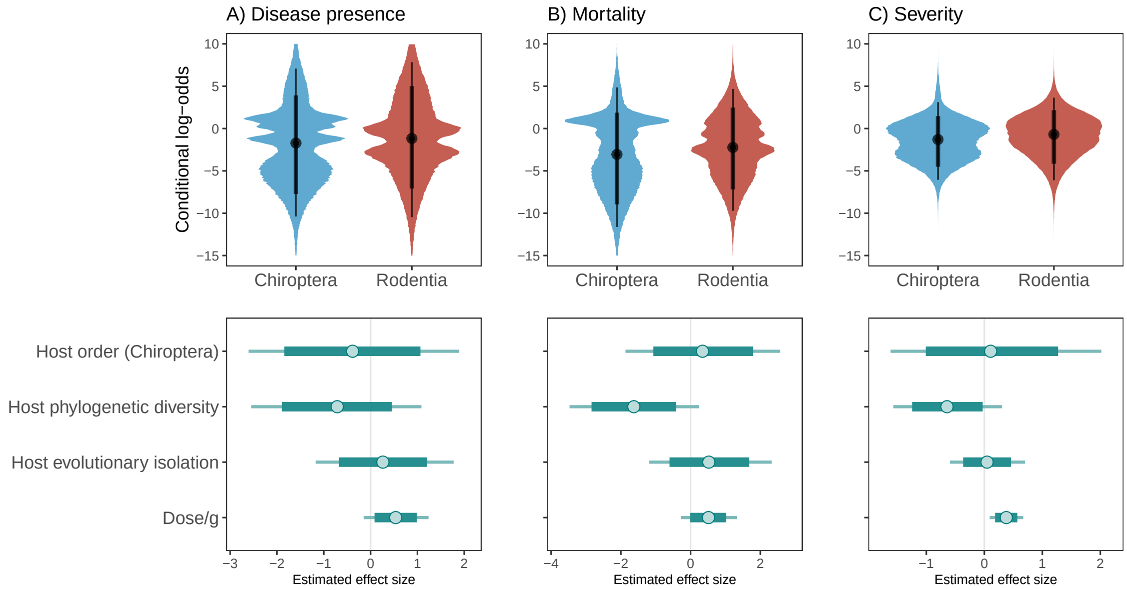

Figure S30: Conditional effect of host order (top row) and estimated effect sizes for predictors (bottom row) for individual-level models of A) disease presence (n=1795), B) mortality (n=972), and C) disease severity (n=1795). Points represent mean estimates, thick bars represent 80% credible intervals, thin bars represent 95% credible intervals. Models are comparable to individual-level full models (Fig. 2) but excluding inoculations via direct injection into the brain or cranium.

##### S2.2.7 Models including only viruses inoculated into both host orders

As a final sensitivity analysis, we refit our individual-level full model after restricting to viruses which were inoculated into both host orders (Fig. S31). This subset included eight viruses (Chikungunya virus, rabies virus, tick-borne encephalitis virus, Kyasanur Forest disease virus, Venezuelan equine encephalitis virus, Rift Valley fever virus, foot-and-mouth disease virus, West Nile virus) inoculated into 41 host species (15 bats; 26 rodents), representing 554 individuals across 47 host-virus combinations. Similar to all other models, there was no consistent effect of host order, however estimated effect sizes for our eco-evolutionary predictors showed increased uncertainty due to decreased effect sizes, while inoculation dose was positive for disease presence and severity.

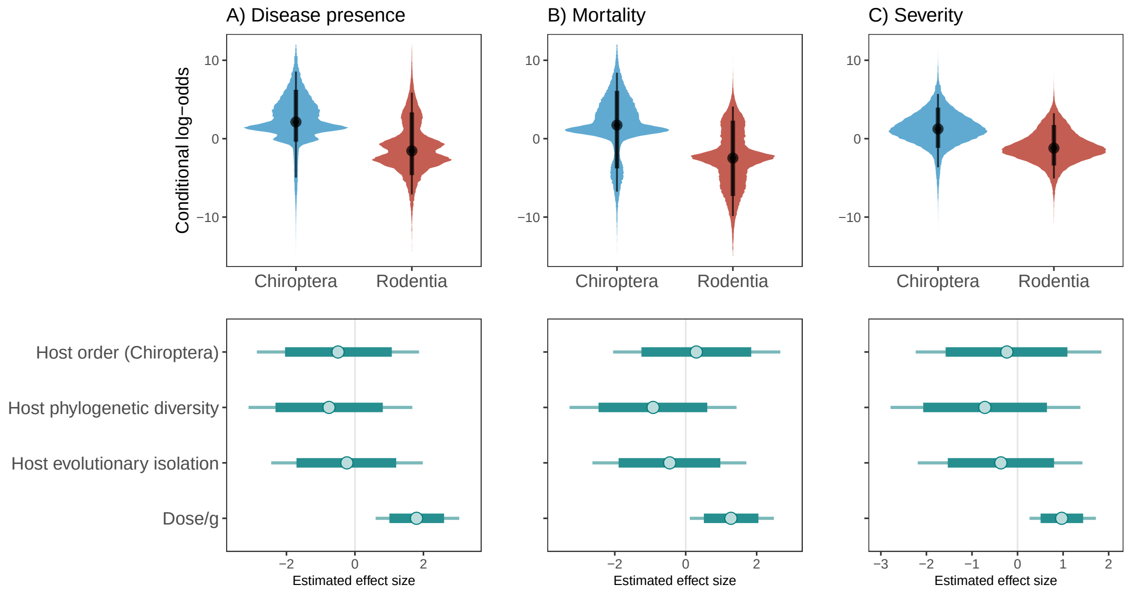

Figure S31: Conditional effect of host order (top row) and estimated effect sizes for predictors (bottom row) for individual-level models of A) disease presence (n=520), B) mortality (n=520), and C) disease severity (n=520). Points represent mean estimates, thick bars represent 80% credible intervals, thin bars represent 95% credible intervals. Models are comparable to individual-level full models (Fig. 2) but restricted to viruses which were inoculated into both host orders.

#### S2.3 Comparison to human case fatality rates

Using human case to fatality rates (CFR), reported by Guth et al., 2022) as a comparable measure of pathogenicity, we evaluated whether inoculation of bats with disproportionately pathogenic viruses could have disguised true differences in tolerance relative to rodents. We find that while bats are more often inoculated with viruses that cause high mortality in human infections (Fig. S32A), there is a large variation in the maximum severity observed in inoculated host species, including large variation in bats among viruses responsible for high human case fatality rates (Fig. S32B). Patterns are similar when considering infection-fatality rates per study combinations (Fig. S33).

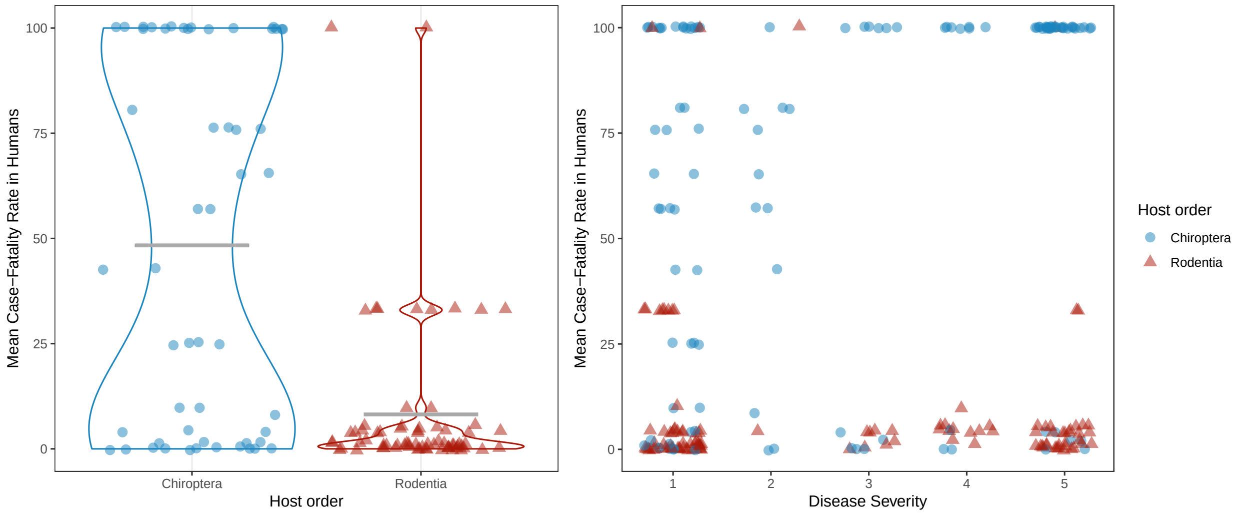

Figure S32: A) The distributions of mean human case-fatality rates (data from Guth et al., 2022) for viruses included in our experimental infection data relative to the inoculated host order, and B) mean human case-fatality rates versus maximum observed disease severity per host-virus-study combination. Colors and shapes indicate the taxonomic orders of inoculated host species. Gray bars indicate means. Note that all viruses with human case-fatality rates above 95% are either lyssaviruses or Whitewater Arroyo mammarenavirus, and those between 42-81% are paramyxoviruses and filoviruses.

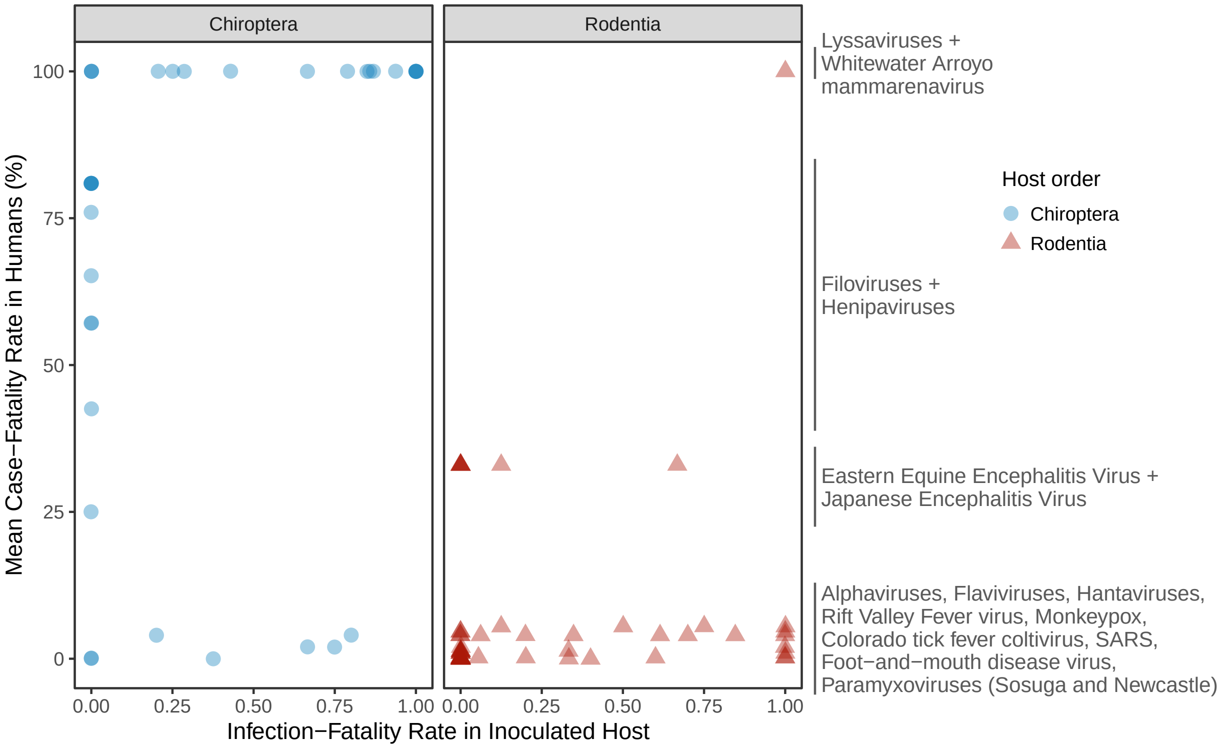

Figure S33: Mean human case-fatality rates (data from Guth et al., 2022) for viruses included in our experimental infection data relative to the inoculated host order versus observed infection-fatality rate per host-virus-study combination. Colors and shapes indicate the taxonomic orders of inoculated host species. Gray bars indicate means. Note that infection-fatality rates for experimental inoculations are available only for studies reporting individual-level disease outcomes.

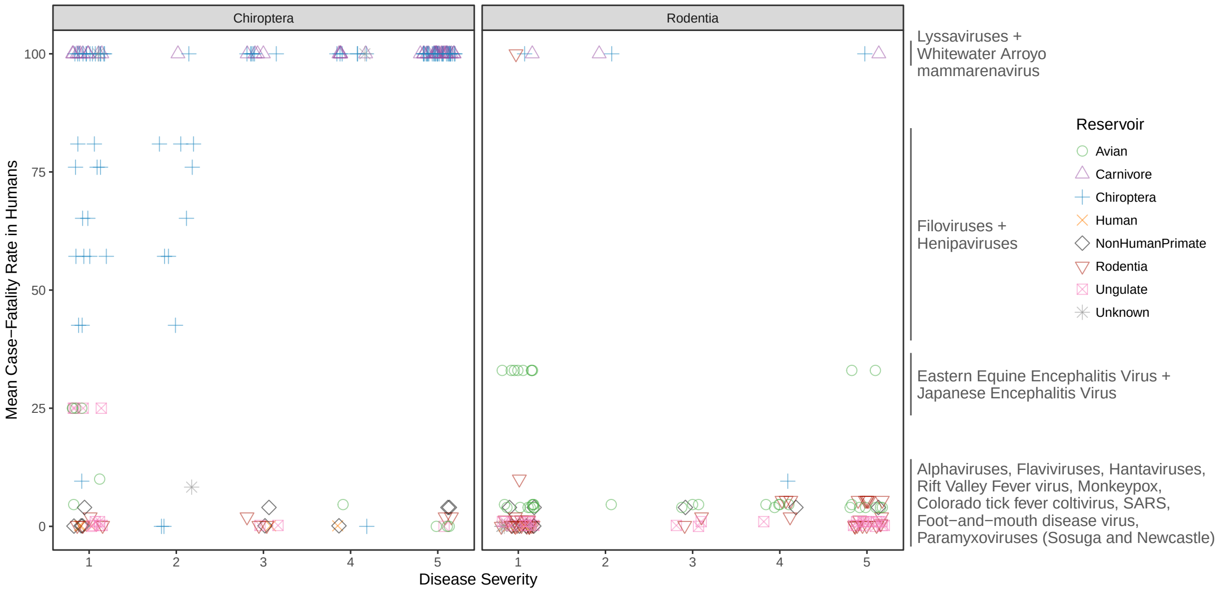

Figure S34: Mean human case fatality rates (data from Guth et al., 2022) versus maximum observed disease severity per experimental infection study. Colors and shapes indicate the taxonomic order of known host reservoir groups (data from Mollentze and Streicker, 2020). Points are separated by the taxonomic order of the experimentally inoculated host species (left box: Chiroptera; right box: Rodentia), and jittered horizontally to improve visualisation of overlapping points.

To investigate whether putatively heightened tolerance in bats may be obscured by bats being more often inoculated with highly pathogenic viruses, we fit a series of models predicting mean human case fatality rates (CFR) as a function of the order of inoculated hosts (Fig. S35). We treated mean human CFR as a proportion (0.0 - 1.0) rather than a percentage and modeled it using zero-one-inflated beta models in *brms*. We used *brms* default priors and assessed convergence of all models using R-hat=1.0 and visual inspection of model fits using posterior predictive checks. All models included a binary predictor of inoculated host order and a study-level hierarchical effect. The base model with no virus hierarchical effects and including all viruses in our study identified that bats are more often infected with high human CFR viruses, which aligns with the raw data presented in Fig S32. However, when we exclude bat-reservoired viruses, or model all viruses but include virus hierarchical effects, we find no difference between bats and rodents. Similarly, reducing models to include only heterologous host-virus pairs (viruses with known reservoirs inoculated into non-reservoirs), we find no difference between bats and rodents in mean CFR. As such, if genuinely heightened tolerance was undetectable in our dataset due to biases in highly pathogenic viruses being selected for experiments, it would mostly reflect bats experiencing less than expected disease from bat-associated viruses – consistent with the expectation that evolutionary association leads to virus-specific adaptations.

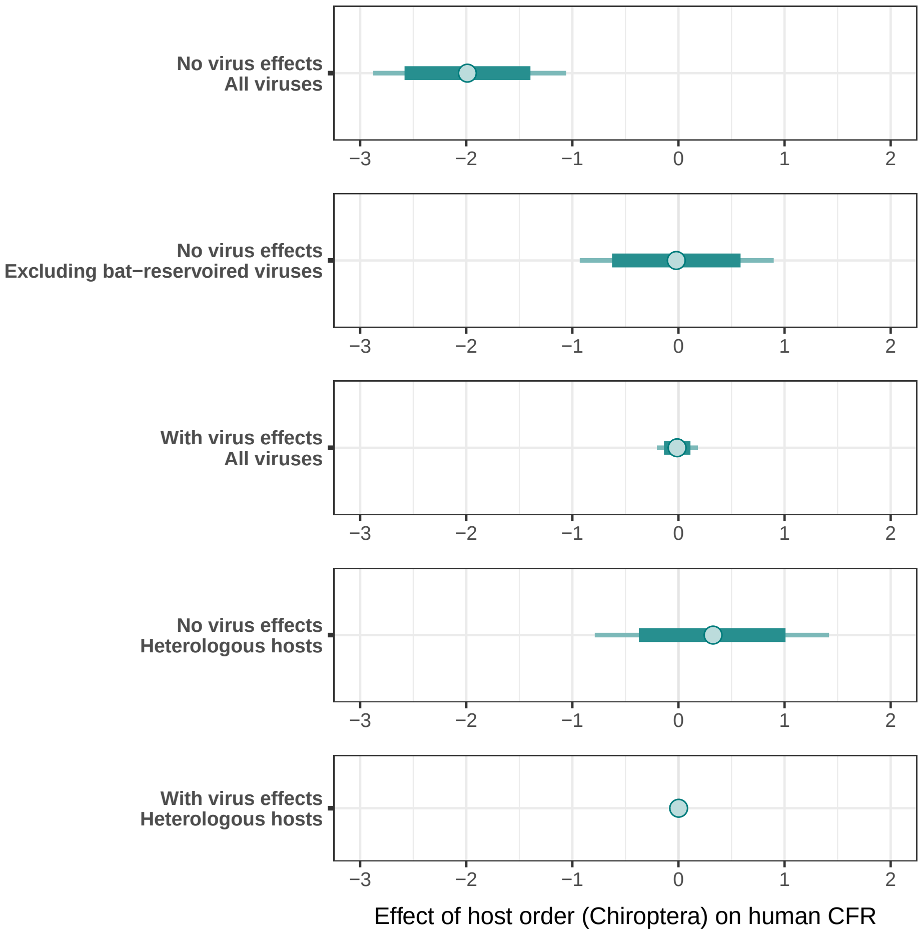

Figure S35: Estimated effect sizes for host order across models predicting mean human case fatality rates (CFR) per virus (data from Guth et al., 2022). Models test whether inoculations involving bat or rodent hosts differ in human CFR by predicting CFR using a binary effect of host order, but varying whether virus taxonomic and non-taxonomic hierarchical effects are included ("With virus effects" versus "No virus effects") and whether viruses known to be associated with bat reservoirs are excluded, or only heterologous hosts (i.e., viruses are inoculated into hosts belonging to a different order than that of putative virus reservoirs) are included.

#### S2.4 Taxa-level hierarchical effects on disease outcomes

It is likely that both certain viruses and certain host species are predisposed to be associated with severe disease or tolerance for reasons that cannot be anticipated *a priori*, and that these effects may be clustered on the respective host/virus phylogenies (Longdon et al., 2015; Imrie et al., 2024). To control for these effects while testing our main predictors, we included host and virus hierarchical effects (described above) into our models of disease occurrence and severity. These effects are visualized below for each measure of disease outcome, illustrating effects for both hosts and viruses for species-level and individual-level models. Interpretation of putative patterns should be made with caution given the small number of observations for some taxa (e.g., host species with only a single viral infection or virus species which were only inoculated into a single host). For each figure, we show the sum of additive phylogenetic and non-phylogenetic effects estimated from the full models containing all other covariates.

**Disease presence**

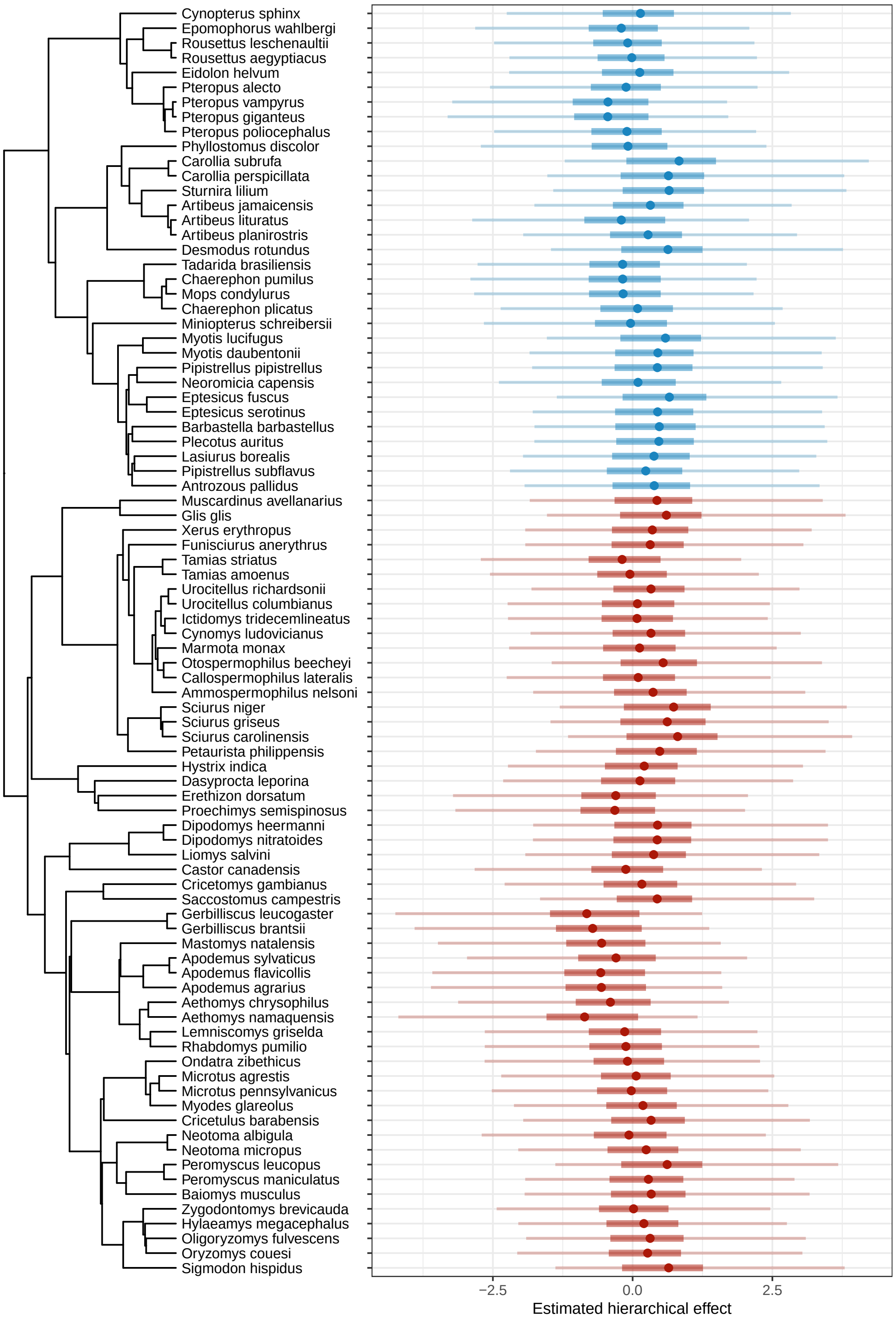

Figure S36: Estimated hierarchical effects for host species based on the species-level model of disease presence with all predictors (Fig. 2). Host-level effects are calculated by summing the estimated phylogenetic and non-phylogenetic hierarchical effects. Dots indicate mean estimates; thick lines are 50% credible intervals; thin lines are 90% credible intervals. Colors indicate host order.

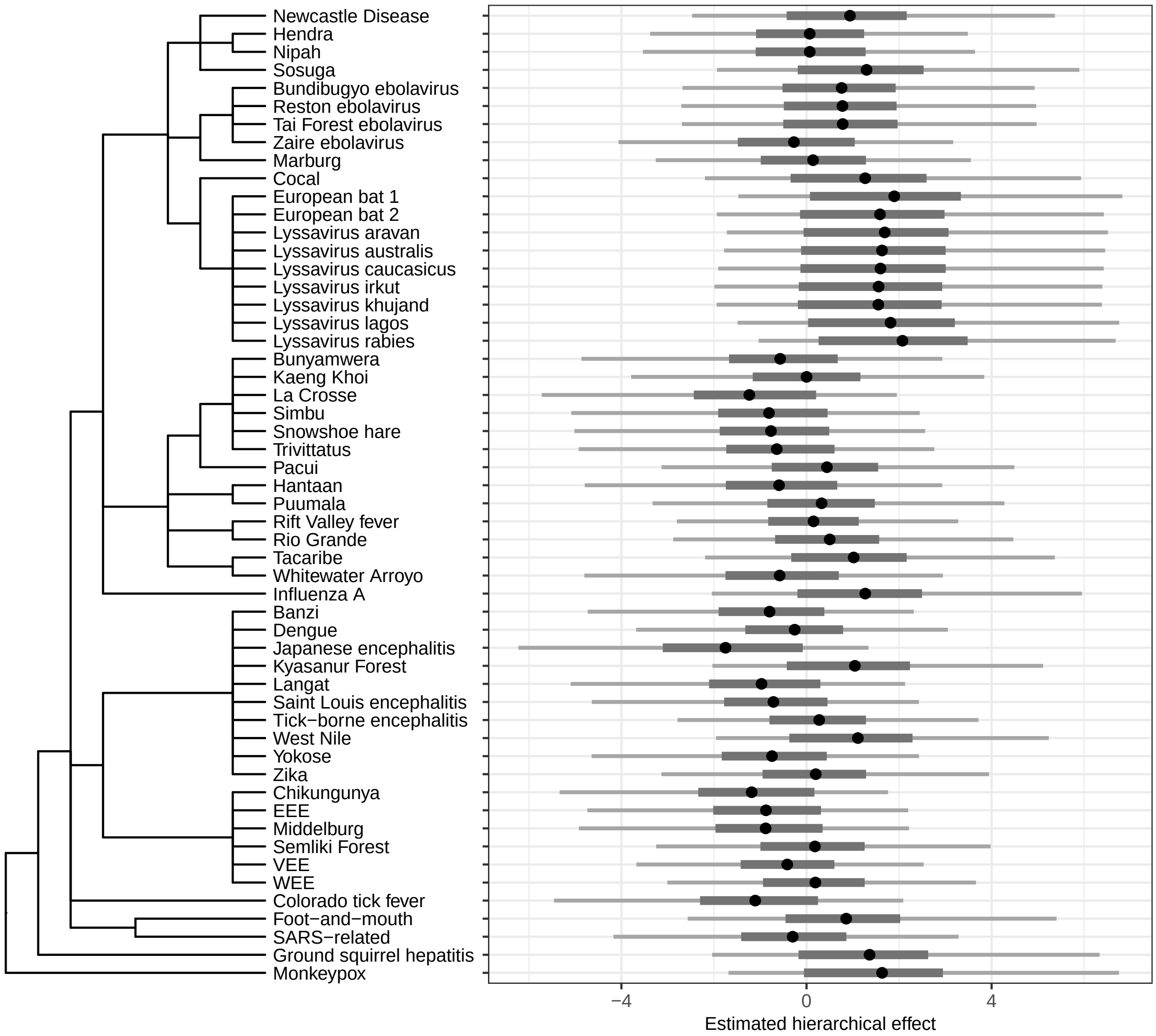

Figure S37: Estimated hierarchical effects for viral taxa based on the species-level model of disease presence with all predictors (Fig. 2). Virus-level effects are calculated by summing the estimated taxonomic and non-taxonomic effects. Dots indicate mean estimates; thick lines are 50% credible intervals; thin lines are 90% credible intervals.

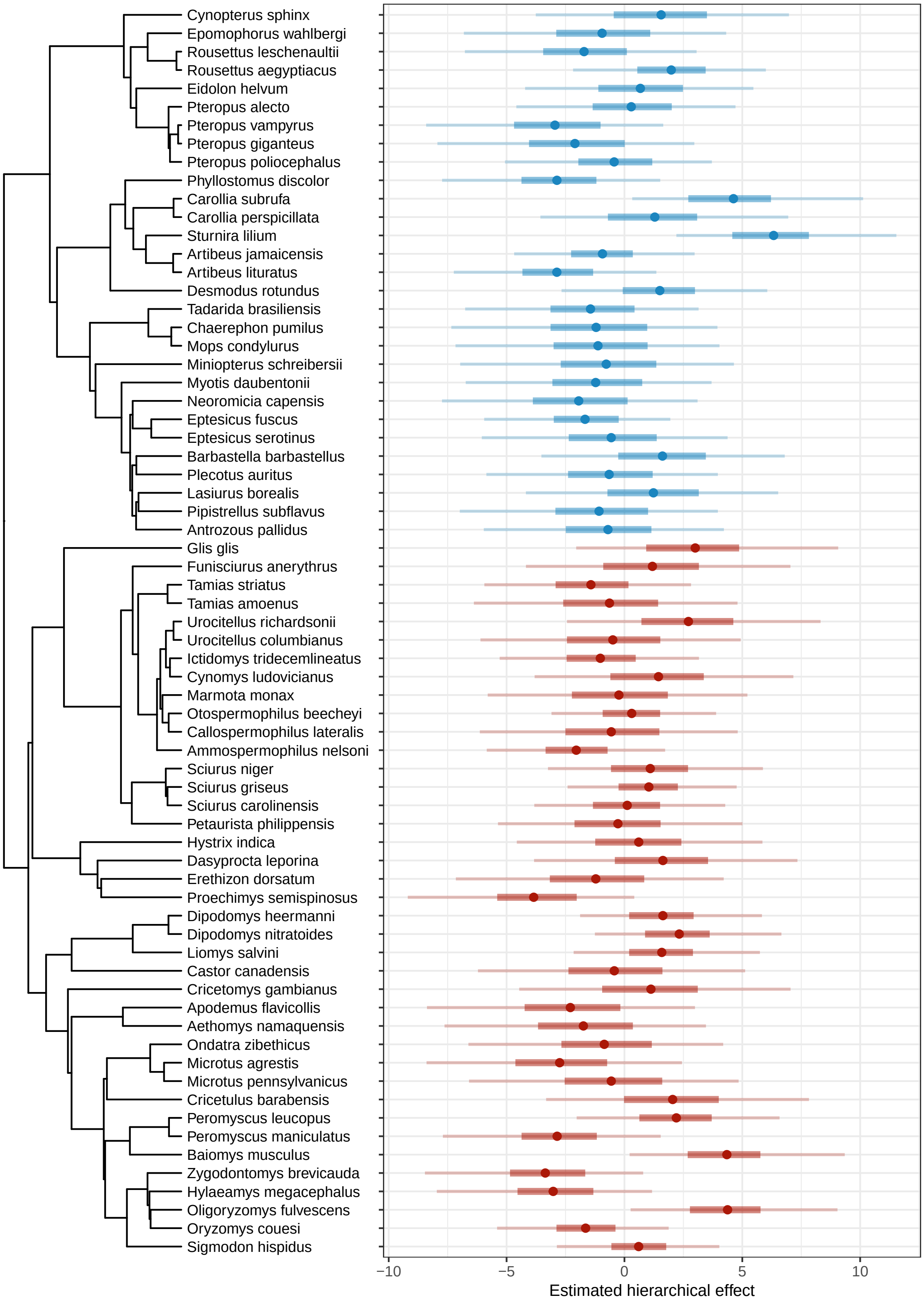

Figure S38: Estimated hierarchical effects for host species based on the individual-level model of disease presence accounting for dose (Fig. 2). Host-level effects are calculated by summing the estimated phylogenetic and non-phylogenetic hierarchical effects. Dots indicate mean estimates; thick lines are 50% credible intervals; thin lines are 90% credible intervals. Colors indicate host order.

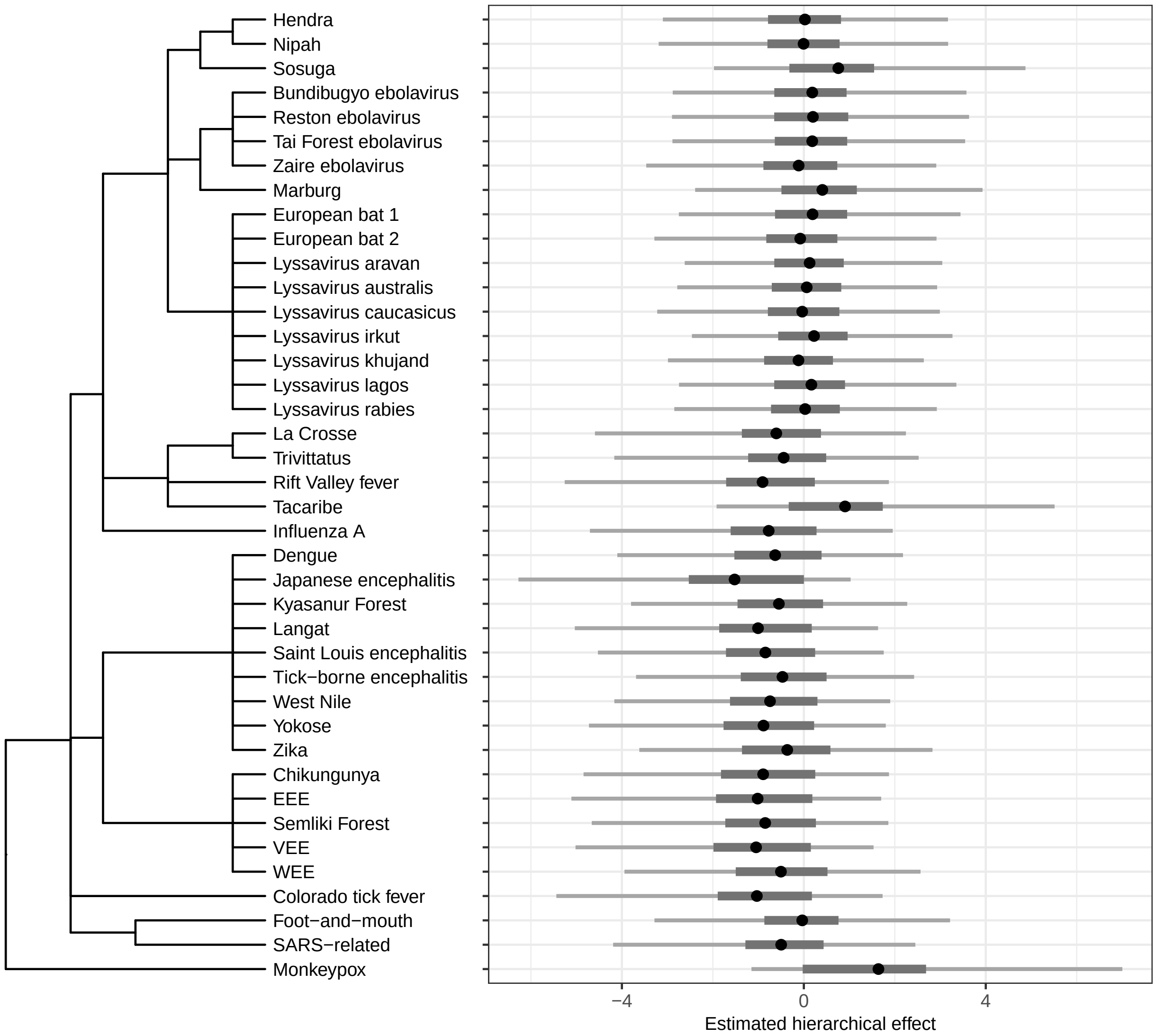

Figure S39: Estimated hierarchical effects for viral taxa based on the individual-level model of disease presence accounting for dose (Fig. 2). Virus-level effects are calculated by summing the estimated taxonomic and non-taxonomic effects. Dots indicate mean estimates; thick lines are 50% credible intervals; thin lines are 90% credible intervals.

#### Mortality

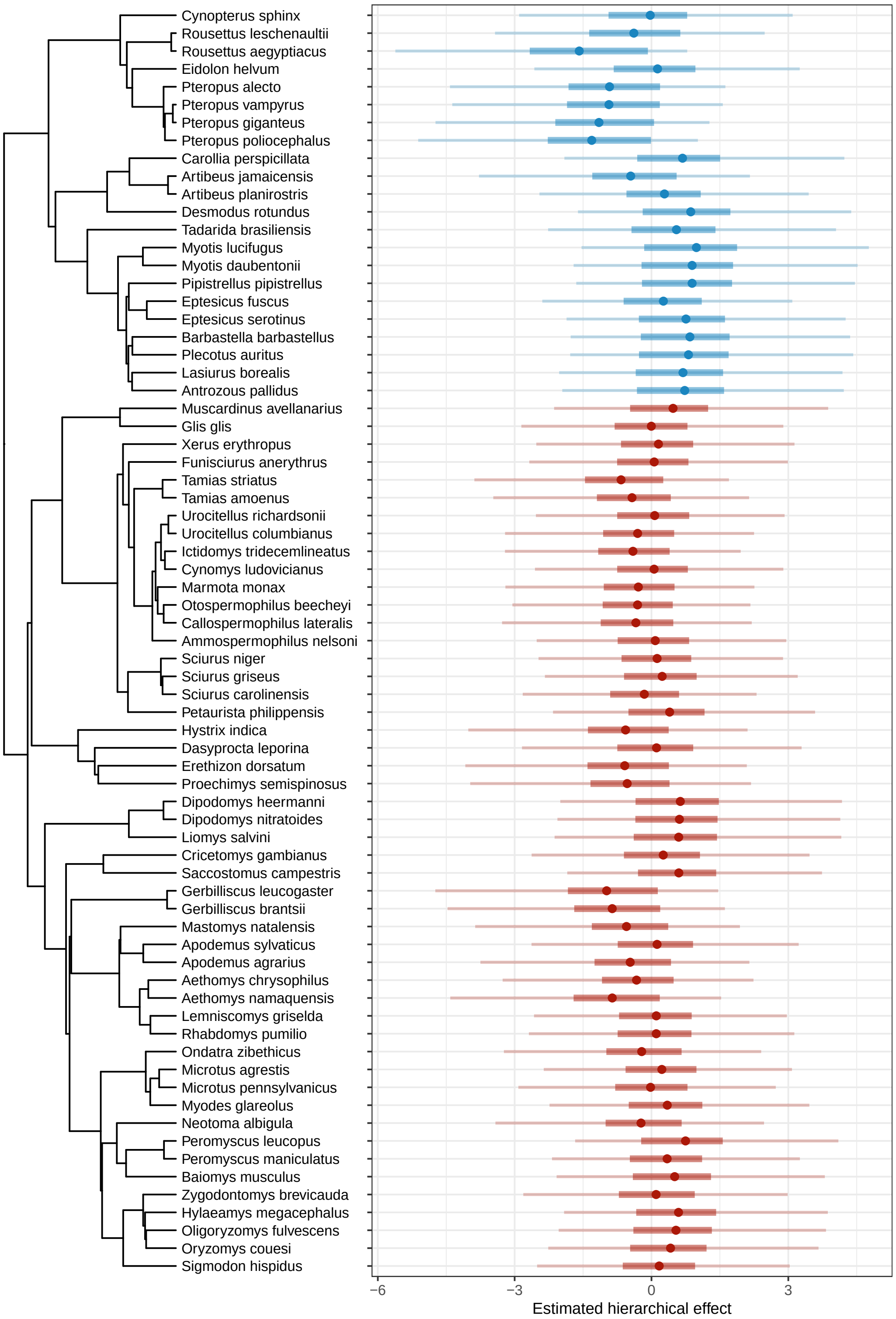

Figure S40: Estimated hierarchical effects for host species based on the species-level model of mortality with all predictors (Fig. 2). Host-level effects are calculated by summing the estimated phylogenetic and non-phylogenetic hierarchical effects. Dots indicate mean estimates; thick lines are 50% credible intervals; thin lines are 90% credible intervals. Colors indicate host order.

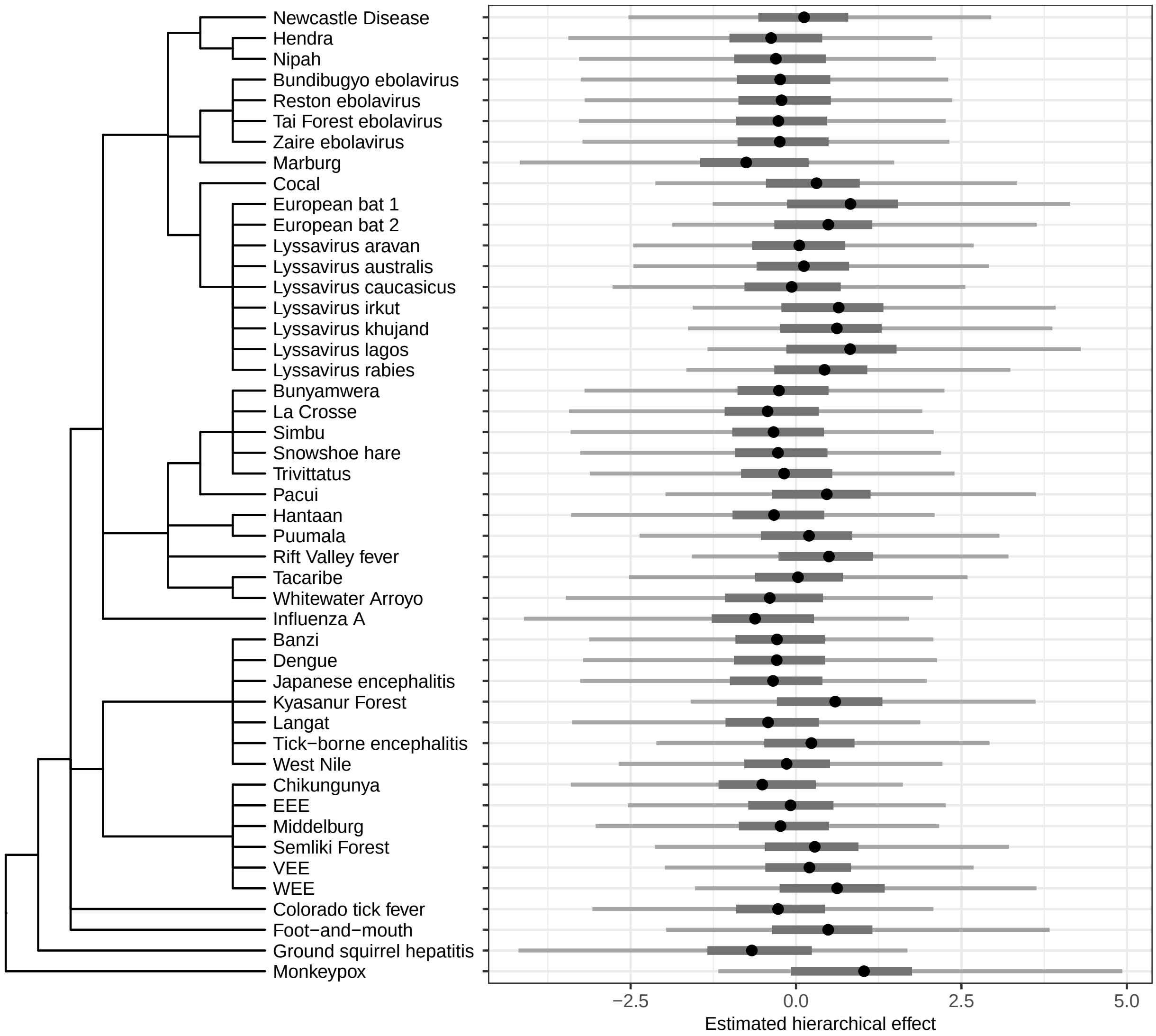

Figure S41: Estimated hierarchical effects for viral taxa based on the species-level model of mortality with all predictors (Fig. 2). Virus-level effects are calculated by summing the estimated taxonomic and non-taxonomic effects. Dots indicate mean estimates; thick lines are 50% credible intervals; thin lines are 90% credible intervals.

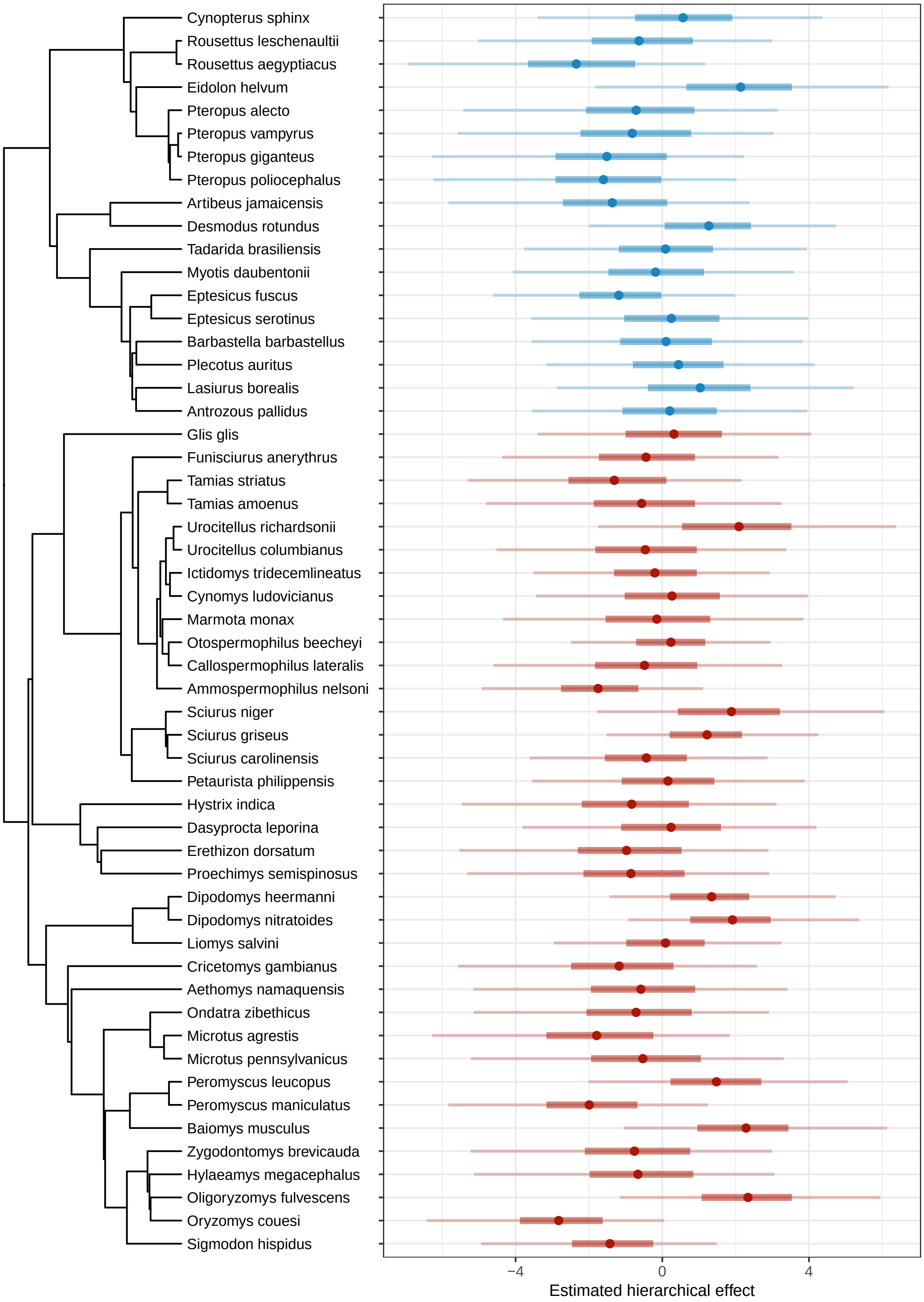

Figure S42: Estimated hierarchical effects for host species based on the individual-level model of mortality accounting for dose (Fig. 2). Host-level effects are calculated by summing the estimated phylogenetic and non-phylogenetic hierarchical effects. Dots indicate mean estimates; thick lines are 50% credible intervals; thin lines are 90% credible intervals. Colors indicate host order.

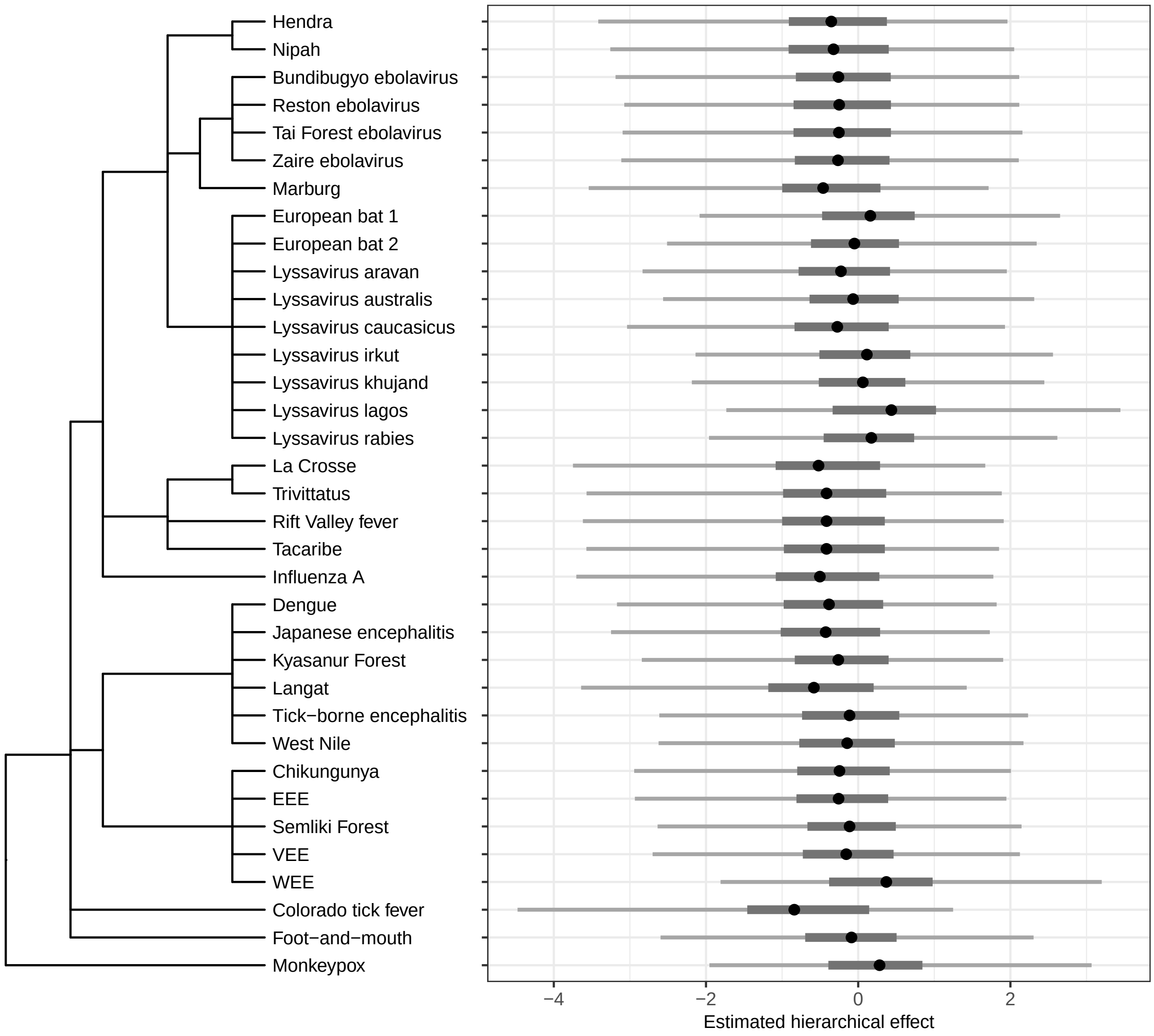

Figure S43: Estimated hierarchical effects for viral taxa based on the individual-level model of mortality accounting for dose (Fig. 2). Virus-level effects are calculated by summing the estimated taxonomic and non-taxonomic effects. Dots indicate mean estimates; thick lines are 50% credible intervals; thin lines are 90% credible intervals.

#### Disease severity

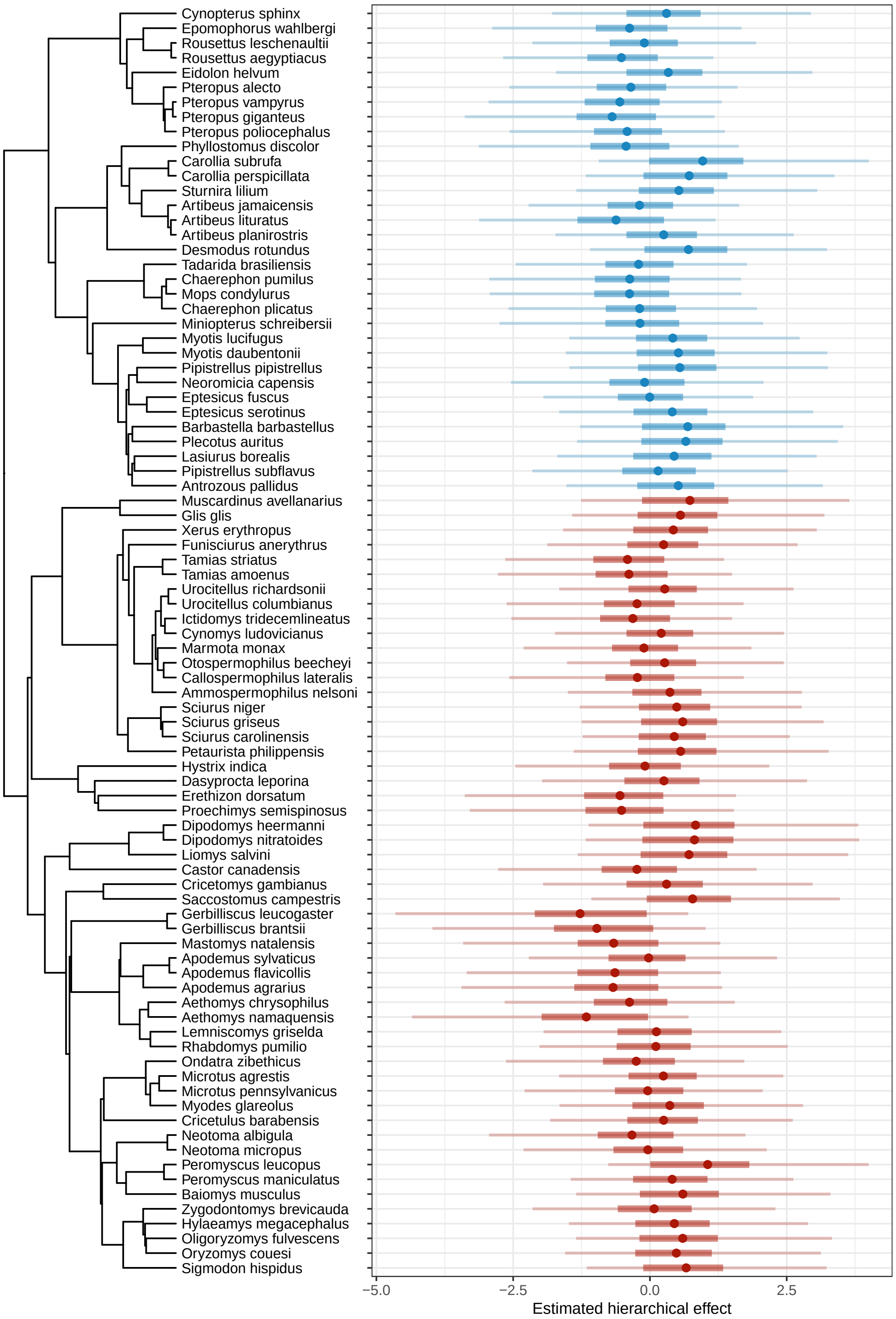

Figure S44: Estimated hierarchical effects for host species based on the species-level model of disease severity with all predictors (Fig. 2). Host-level effects are calculated by summing the estimated phylogenetic and non-phylogenetic hierarchical effects. Dots indicate mean estimates; thick lines are 50% credible intervals; thin lines are 90% credible intervals. Colors indicate host order.

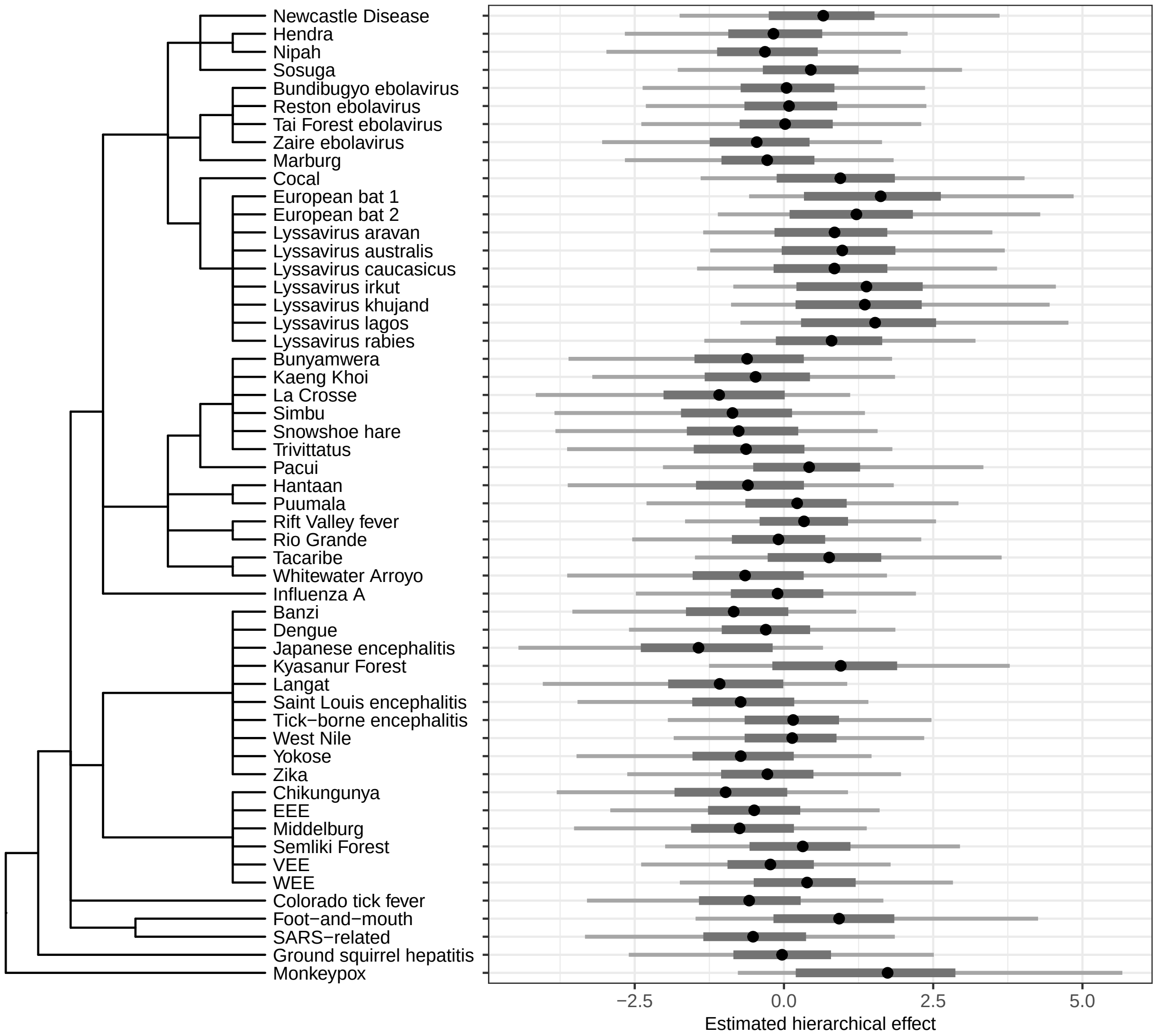

Figure S45: Estimated hierarchical effects for viral taxa based on the species-level model of disease severity with all predictors (Fig. 2). Virus-level effects are calculated by summing the estimated taxonomic and non-taxonomic effects. Dots indicate mean estimates; thick lines are 50% credible intervals; thin lines are 90% credible intervals.

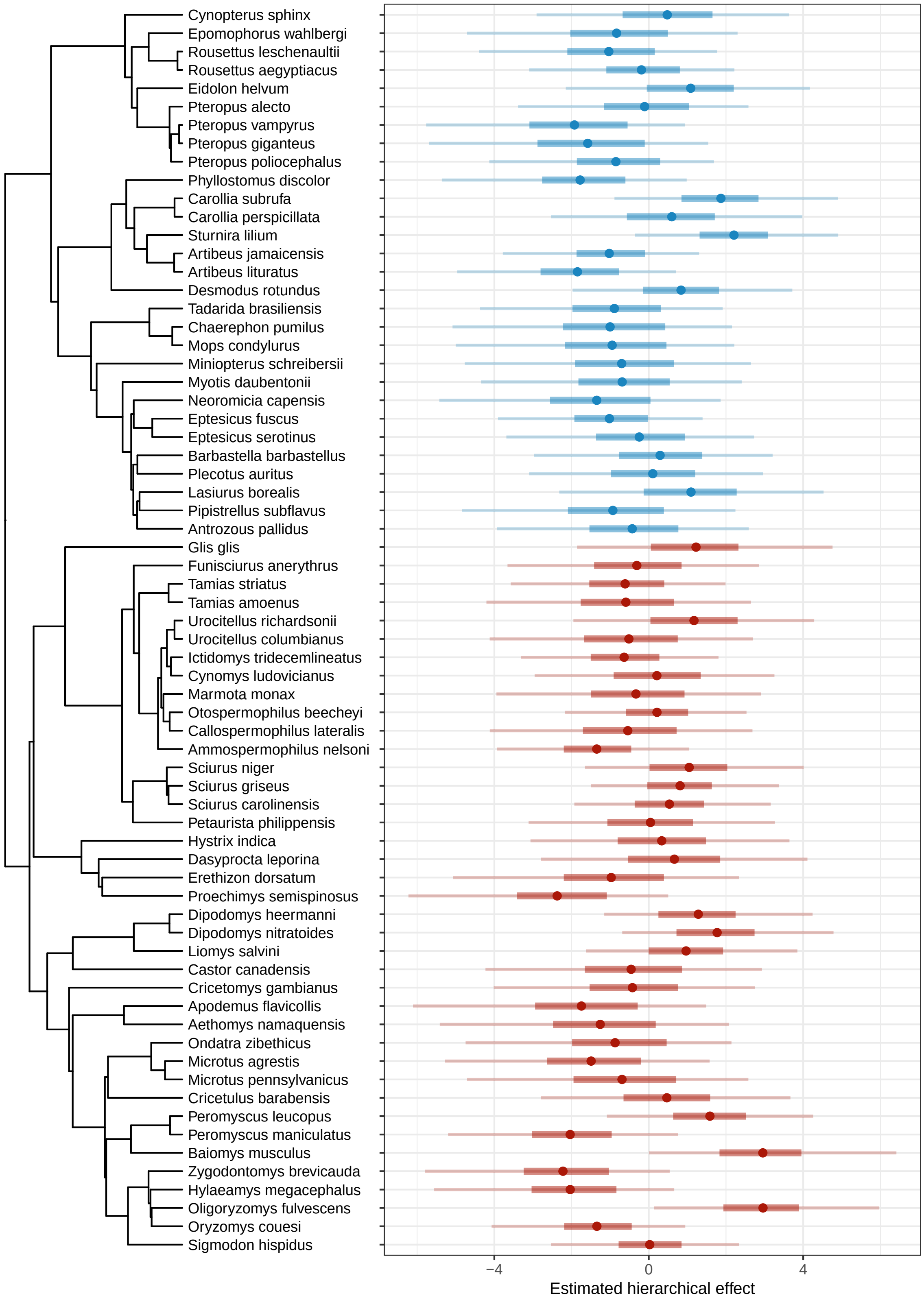

Figure S46: Estimated hierarchical effects for host species based on the individual-level model of disease severity accounting for dose (Fig. 2). Host-level effects are calculated by summing the estimated phylogenetic and non-phylogenetic hierarchical effects. Dots indicate mean estimates; thick lines are 50% credible intervals; thin lines are 90% credible intervals. Colors indicate host order.

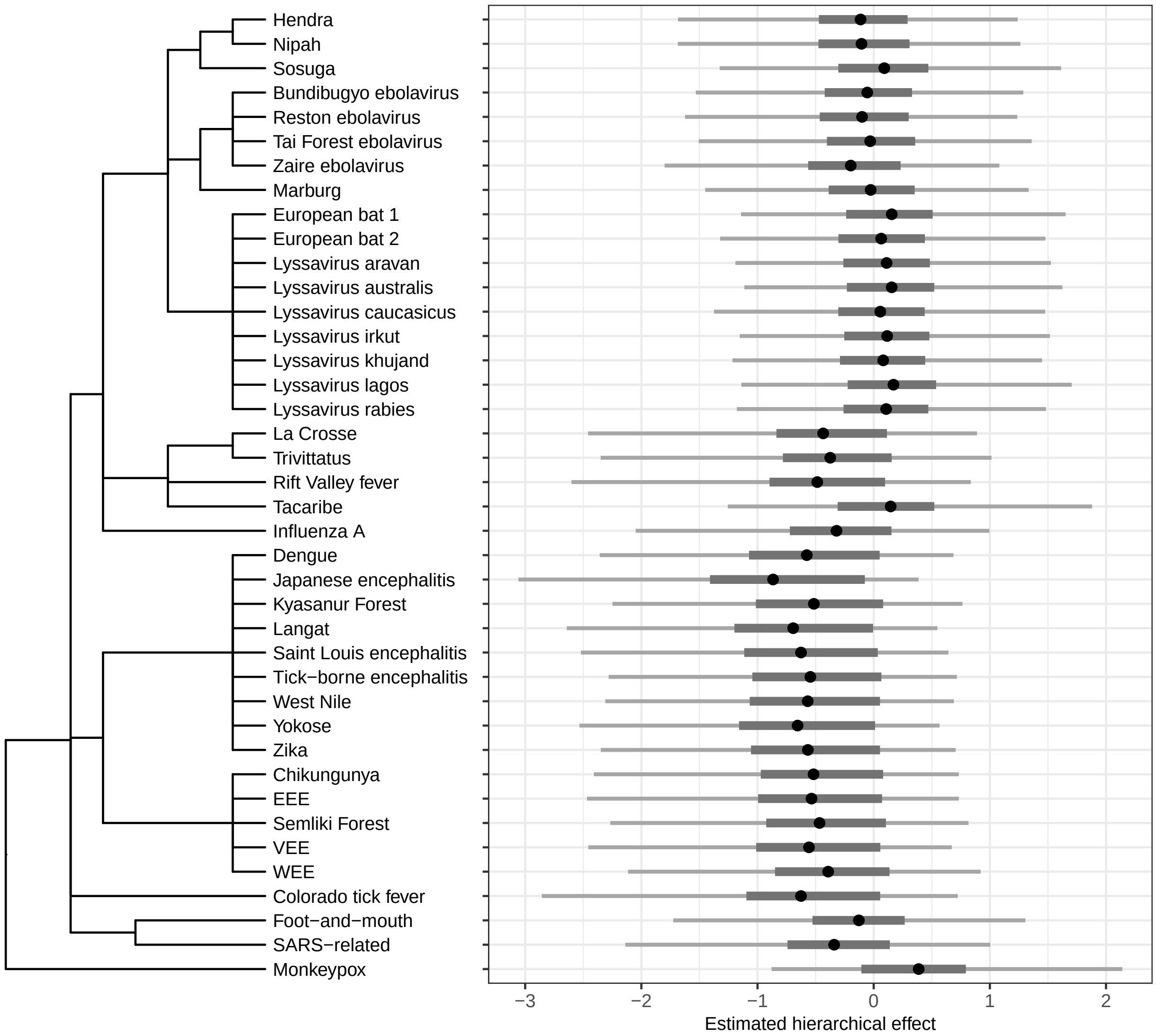

Figure S47: Estimated hierarchical effects for viral taxa based on the individual-level model of disease severity accounting for dose (Fig. 2). Virus-level effects are calculated by summing the estimated taxonomic and non-taxonomic effects. Dots indicate mean estimates; thick lines are 50% credible intervals; thin lines are 90% credible intervals.

#### S3 Articles reporting disease outcomes for susceptible host species

Addy, P. A. K., Lule, M., and Sekyalo, E. 1982. Experimental infection of Roussettus aegyptiacus E. geoffroy (chiroptera) with Japanese B encephalitis virus. *Annales de l'Institut Pasteur / Virologie*, 133(2): 133–137.

Aguilar-Setien, A., Loza-Rubio, E., Salas-Rojas, M., Brisseau, N., Cliquet, F., Pastoret, P.-P., Rojas-Dotor, S., Tesoro, E., and Kretschmer, R. 2005. Salivary excretion of rabies virus by healthy vampire bats. *Epidemiology and Infection*, 133(3):517–522.

Aitken, T. H. G., Woodall, J. P., De Andrade, A. H. P., Bensabath, G., and Shope, R. E. 1975. Pacui virus, phlebotomine flies, and small mammals in Brazil: An epidemiological case study. *The American Journal of Tropical Medicine and Hygiene*, 24(2):358–368.

Almeida, M. F., Martorelli, L. F. A., Aires, C. C., Sallum, P. C., Durigon, E. L., and Massad, E. 2005. Experimental rabies infection in haematophagous bats Desmodus rotundus. *Epidemiology and Infection*, 133(3):523–527.

Amman, B. R., Jones, M. E. B., Sealy, T. K., Uebelhoer, L. S., Schuh, A. J., Bird, B. H., Coleman-McCray, J. D., Martin, B. E., Nichol, S. T., and Towner, J. S. 2015. Oral shedding of Marburg virus in experimentally infected Egyptian fruit bats (Rousettus aegyptiacus). *Journal of Wildlife Diseases*, 51(1):113–124.

Amman, B. R., Schuh, A. J., Sealy, T. K., Spengler, J. R., Welch, S. R., Kirejczyk, S. G. M., Albariño, C. G., Nichol, S. T., and Towner, J. S. 2020. Experimental infection of Egyptian rousette bats (Rousettus aegyptiacus) with Sosuga virus demonstrates potential transmission routes for a bat-borne human pathogenic paramyxovirus. *PLOS Neglected Tropical Diseases*, 14(3):e0008092.

Aparecida, M. C., Souza, M., de, A. F., Nassar, C., Cortez, A., Sakai, T., Itou, T., and Sequetin, E. M. 2009. Experimental infection of vampire bats Desmodus rotundus (E. Geoffroy) maintained in captivity by feeding defibrinated blood added with rabies virus. *São Paulo*, 46(2):92–100.

Baer, G. M. and Bales, G. L. 1967. Experimental rabies infection in the Mexican freetail bat. *Journal of Infectious Diseases*, 117(1):82–90.

Begeman, L., Suu-Ire, R., Banyard, A. C., Drosten, C., Eggerbauer, E., Freuling, C. M., Gibson, L., Goharriz, H., Horton, D. L., Jennings, D., Marston, D. A., Ntiamoa-Baidu, Y., Riesle Sbarbaro, S., Selden, D., Wise, E. L., Kuiken, T., Fooks, A. R., Müller, T., Wood, J. L. N., and Cunningham, A. A. 2020. Experimental Lagos bat virus infection in straw-colored fruit bats: A suitable model for bat rabies in a natural reservoir species. *PLOS Neglected Tropical Diseases*, 14(12):e0008898.

Bertzbach, L. D., Vladimirova, D., Dietert, K., Abdelgawad, A., Gruber, A. D., Osterrieder, N., and Trimpert, J. 2021. SARS-CoV-2 infection of Chinese hamsters (Cricetulus griseus) reproduces COVID-19 pneumonia in a well-established small animal model. *Transboundary and Emerging Diseases*, 68(3): 1075–1079.

Bhat, H. R., Sreenivasan, M. A., Goverdhan, M. K., Naik, S. V., and Banerjee, K. 1976. Susceptibility of Hystrix indica Kerr, 1792, Indian crested porcupine (Rodentia Hystricidae) to KFD virus. *The Indian Journal of Medical Research*, 64(11):1566–1570.

Bhat, H. R., Sreenivasan, M. A., and Naik, S. V. 1979. Susceptibility of common giant flying squirrel to experimental infection with KFD virus. *The Indian Journal of Medical Research*, 69:697–700.

Burgdorfer, W. 1959. Colorado tick fever the behavior of CTF virus in the porcupine. *Journal of Infectious Diseases*, 104(1):101–104.

Burgdorfer, W. 1960. Colorado tick fever II. The behavior of Colorado tick fever virus in rodents. *Journal of Infectious Diseases*, 107(3):384–388.

Burns, K. F., Shelton, D. F., and Grogan, E. W. 1958. Bat rabies: Experimental host transmission studies. *Annals of the New York Academy of Sciences*, 70(3):452–466.

Ciminski, K., Ran, W., Gorka, M., Lee, J., Malmlov, A., Schinköthe, J., Eckley, M., Murrieta, R. A., Aboellail, T. A., Campbell, C. L., Ebel, G. D., Ma, J., Pohlmann, A., Franzke, K., Ulrich, R., Hoffmann, D., García-Sastre, A., Ma, W., Schountz, T., Beer, M., and Schwemmle, M. 2019. Bat influenza viruses transmit among bats but are poorly adapted to non-bat species. *Nature Microbiology*, 4(12):2298–2309.

Cogswell-Hawkinson, A., Bowen, R., James, S., Gardiner, D., Calisher, C. H., Adams, R., and Schountz, T. 2012. Tacaribe virus causes fatal infection of an ostensible reservoir host, the Jamaican fruit bat. *Journal of Virology*, 86(10):5791–5799.

Constantine, D. G. 1966. Transmission experiments with bat rabies isolates: Reactions of certain carnivora, opossum, rodents, and bats to rabies virus of red bat origin when exposed by bat bite or by intramuscular inoculation. *American Journal of Veterinary Research*, 27(116):24–32.

Davis, A., Bunning, M., Gordy, P., Panella, N., Blitvich, B., and Bowen, R. 2005. Experimental and natural infection of North American bats with West Nile virus. *The American Journal of Tropical Medicine and Hygiene*, 73(2):467–469.

Davis, A. D., Jarvis, J. A., Pouliott, C., and Rudd, R. J. 2013. Rabies virus infection in Eptesicus fuscus bats born in baptivity (naïve bats). *PLoS ONE*, 8(5):e64808.

Deardorff, E. R., Forrester, N. L., Travassos Da Rosa, A. P., Estrada-Franco, J. G., Navarro-Lopez, R., Tesh, R. B., and Weaver, S. C. 2009. Experimental infection of potential reservoir hosts with Venezuelan equine encephalitis virus, Mexico. *Emerging Infectious Diseases*, 15(4):519–525.

Donaldson, A. I. 1970. Bats as possible maintenance hosts for vesicular stomatitis virus. *American Journal of Epidemiology*, 92(2):132–136.

Ellison, J. A., Johnson, S. R., Kuzmina, N., Gilbert, A., Carson, W. C., VerCauteren, K. C., and Rupprecht, C. E. 2013. Multidisciplinary approach to epizootiology and pathogenesis of bat rabies viruses in the United States. *Zoonoses and Public Health*, 60(1):46–57.

Ernek, E., Kozuch, O., Lichard, M., Nosek, J., and Albrecht, P. 1963. Experimental infection of Clethrionomys glareolus and Apodemus flavicollis with tick-borne encephalitis virus. *Acta Virologica*, 7:434—436.

Falendysz, E. A., Lopera, J. G., Doty, J. B., Nakazawa, Y., Crill, C., Lorenzsonn, F., Kalemba, L. N., Ronderos, M. D., Mejia, A., Malekani, J. M., Karem, K., Carroll, D. S., Osorio, J. E., and Rocke, T. E. 2017. Characterization of Monkeypox virus infection in African rope squirrels (Funisciurus sp.). *PLOS Neglected Tropical Diseases*, 11(8):e0005809.

Federer, K. E. 1969. Susceptibility of the Agouti (Dasyprocta Aguti) to Foot-and-Mouth Disease Virus. *Zentralblatt für Veterinärmedizin Reihe B*, 16(9):847–854.

Findlay, G. 1932. Rift Valley fever or enzootic hepatitis. *Transactions of the Royal Society of Tropical Medicine and Hygiene*, 25(4):229–262.

Fooks, A. R., Johnson, N., Neubert, L., Kaipf, I., Denzinger, A., Franka, R., and Rupprecht, C. E. 2009. Detection of high levels of European bat lyssavirus type-1 viral RNA in the thyroid gland of experimentally-infected Eptesicus fuscus bats. *Zoonoses Public Health*, 56:270–277.

Franka, R., Johnson, N., Müller, T., Vos, A., Neubert, L., Freuling, C., Rupprecht, C. E., and Fooks, A. R. 2008. Susceptibility of North American big brown bats (Eptesicus fuscus) to infection with European bat lyssavirus type 1. *Journal of General Virology*, 89(8):1998–2010.

Freuling, C., Vos, A., Johnson, N., Kaipf, I., Denzinger, A., Neubert, L., Mansfield, K., Hicks, D., Nuñez, A., Tordo, N., Rupprecht, C. E., Fooks, A. R., and Müller, T. 2009. Experimental infection of serotine bats (Eptesicus serotinus) with European bat lyssavirus type 1a. *Journal of General Virology*, 90(10): 2493–2502.

Fulhorst, C. F., Milazzo, M. L., Bradley, R. D., and Peppers, L. L. 2001. Experimental infection of Neotoma albigula (Muridae) with Whitewater Arroyo virus (Arenaviridae). *The American journal of tropical medicine and hygiene*, 65(2):147–151.

Ganem, D., Weiser, B., Barchuk, A., Brown, R. J., and Varmus, H. E. 1982. Biological characterization of acute infection with ground squirrel hepatitis virus. *Journal of Virology*, 44(1):366–373.

Gerrard, D. L., Hawkinson, A., Sherman, T., Modahl, C. M., Hume, G., Campbell, C. L., Schountz, T., and Frietze, S. 2017. Transcriptomic signatures of Tacaribe virus-infected Jamaican fruit bats. *mSphere*, 2 (5):e00245–17.

Gómez, A., Kramer, L. D., Dupuis, A. P., Kilpatrick, A. M., Davis, L. J., Jones, M. J., Daszak, P., and Aguirre, A. A. 2008. Experimental infection of eastern gray squirrels (Sciurus carolinensis) with West Nile virus. *The American Journal of Tropical Medicine and Hygiene*, 79(3):447–451.

Gorka, M., Schinköthe, J., Ulrich, R., Ciminski, K., Schwemmle, M., Beer, M., and Hoffmann, D. 2020. Characterization of experimental oro-nasal inoculation of Seba's short-tailed bats (Carollia perspicillata) with bat influenza A virus H18N11. *Viruses*, 12(2):232.

Halpin, K., Hyatt, A. D., Fogarty, R., Middleton, D., Bingham, J., Epstein, J. H., Rahman, S. A., Hughes, T., Smith, C., Field, H. E., and Daszak, P. 2011. Pteropid bats are confirmed as the reservoir hosts of henipaviruses: A comprehensive experimental study of virus transmission. *The American Society of Tropical Medicine and Hygiene*, 85(5):946–951.

Halwe, N. J., Gorka, M., Hoffmann, B., Rissmann, M., Breithaupt, A., Schwemmle, M., Beer, M., Kandeil, A., Ali, M. A., Kayali, G., Hoffmann, D., and Balkema-Buschmann, A. 2021. Egyptian fruit bats (Rousettus aegyptiacus) were resistant to experimental inoculation with avian-origin influenza A virus of subtype H9N2, but are susceptible to experimental infection with bat-borne H9N2 virus. *Viruses*, 13(4): 672.

Hardy, J. L., Reeves, W. C., Rush, W. A., and Nir, Y. D. 1974. Experimental infection with western equine encephalomyelitis virus in wild rodents indigenous to Kern county, California. *Infection and Immunity*, 10(3):553–564.

Hughes, G. J., Kuzmin, I. V., Schmitz, A., Blanton, J., Manangan, J., Murphy, S., and Rupprecht, C. E. 2006. Experimental infection of big brown bats (Eptesicus fuscus) with Eurasian bat lyssaviruses Aravan, Khujand, and Irkut virus. *Archives of Virology*, 151(10):2021–2035.

Hutson, C. L., Nakazawa, Y. J., Self, J., Olson, V. A., Regnery, R. L., Braden, Z., Weiss, S., Malekani, J., Jackson, E., Tate, M., Karem, K. L., Rocke, T. E., Osorio, J. E., Damon, I. K., and Carroll, D. S. 2015. Laboratory investigations of African pouched rats (Cricetomys gambianus) as a potential reservoir host species for Monkeypox virus. *PLOS Neglected Tropical Diseases*, 9(10):e0004013.

Jackson, F. R., Turmelle, A. S., Farino, D. M., Franka, R., McCracken, G. F., and Rupprecht, C. E. 2008. Experimental rabies virus infection of big brown bats. *Journal of Wildlife Diseases*, 44(3):612–621.

Johnson, N., Vos, A., Neubert, L., Freuling, C., Mansfield, K. L., Kaipf, I., Denzinger, A., Hicks, D., Núñez, A., Franka, R., Rupprecht, C. E., Müller, T., and Fooks, A. R. 2008. Experimental study of European bat lyssavirus type-2 infection in Daubenton's bats (Myotis daubentonii). *Journal of General Virology*, 89 (11):2662–2672.

Jones, M., Schuh, A., Amman, B., Sealy, T., Zaki, S., Nichol, S., and Towner, J. 2015. Experimental inoculation of Egyptian rousette bats (Rousettus aegyptiacus) with viruses of the ebolavirus and marburgvirus genera. *Viruses*, 7(7):3420–3442.

Karstad, L., Spalatin, J., and Hanson, R. 1961. Natural and experimental infections with the virus of eastern encephalitis in wild rodents from Wisconsin, Minnesota, Michigan and Georgia. *Zoonoses Research*, 1 (5):87–96.

Keckler, M. S., Salzer, J. S., Patel, N., Townsend, M. B., Nakazawa, Y. J., Doty, J. B., Gallardo-Romero, N. F., Satheshkumar, P. S., Carroll, D. S., Karem, K. L., and Damon, I. K. 2020. IMVAMUNE® and ACAM2000® provide different protection against disease when administered postexposure in an intranasal monkeypox challenge prairie dog model. *Vaccines*, 8(3):396.

Kirejczyk, S. G. M., Amman, B. R., Schuh, A. J., Sealy, T. K., Albariño, C. G., Zhang, J., Brown, C. C., and Towner, J. S. 2022. Histopathologic and immunohistochemical evaluation of induced lesions, tissue tropism and host responses following experimental infection of Egyptian rousette bats (Rousettus aegyptiacus) with the zoonotic paramyxovirus, Sosuga virus. *Viruses*, 14(6):1278.

Kozuch, O., Nosek, J., Ernek, E., Lichard, M., and Albrecht, P. 1963. Persistence of tick-borne encephalitis virus in hibernating hedgehogs and dormice. *Acta Virologica*, 7:430–433.

Kuzmin, I. V. and Botvinkin, A. D. 1996. The behaviour of bats Pipistrellus pipistrellus after experimental inoculation with rabies and rabies-like viruses, and some aspects of pathogenesis. *Myotis*, 34:93–99.

Kuzmin, I. V., Franka, R., and Rupprecht, C. E. 2008. Experimental infection of big brown bats (Eptesicus fuscus) with West Caucasian bat virus (WCBV). *Developments in Biologicals*, 131:327–337.

La Motte, L. C. 1958. Japanese B encephalitis in bats during simulated hibernation. *American Journal of Epidemiology*, 67(1):101–108.

Lee, H. W., Lee, P. W., and Johnson, K. M. 1978. Isolation of the etiologic agent of Korean hemorrhagic fever. *The Journal of Infectious Diseases*, 137(3):298–308.

Leung, M. K., McLintock, J., and Iversen, J. 1978. Intranasal exposure of the Richardson's ground squirrel to western equine encephalomyelitis virus. *Can. J. comp. Med.*, 42.

Lord, R. D., De Chaurell, A. O., and Elsy, C. A. 1986. Experimental infection of vampire bats with foot and mouth disease virus. *Journal of Wildlife Diseases*, 22(3):413–414.

Madrières, S., Tatard, C., Murri, S., Vulin, J., Galan, M., Piry, S., Pulido, C., Loiseau, A., Artige, E., Benoit, L., Leménager, N., Lakhdar, L., Charbonnel, N., Marianneau, P., and Castel, G. 2020. How bank vole-PUUV interactions influence the eco-evolutionary processes driving nephropathia epidemica epidemiology—an experimental and genomic approach. *Pathogens*, 9(10):789.

Malmlov, A., Bantle, C., Aboellail, T., Wagner, K., Campbell, C. L., Eckley, M., Chotiwan, N., Gullberg, R. C., Perera, R., Tjalkens, R., and Schountz, T. 2019. Experimental Zika virus infection of Jamaican fruit bats (Artibeus jamaicensis) and possible entry of virus into brain via activated microglial cells. *PLOS Neglected Tropical Diseases*, 13(2):e0007071.

Marion, P. L., Knight, S. S., Salazar, F. H., Popper, H., and Robinson, W. S. 1983. Ground squirrel hepatitis virus infection. *Hepatology*, 3(4):519–527.

Mccoll, K., Chamberlain, T., Lunt, R., Newberry, K., Middleton, D., and Westbury, H. 2002. Pathogenesis studies with Australian bat lyssavirus in grey-headed flying foxes (Pteropus poliocephalus). *Australian Veterinary Journal*, 80(10):636–641.

McIntosh, B. 1961. Susceptibility of some African wild rodents to infection with various arthropod-borne viruses. *Transactions of the Royal Society of Tropical Medicine and Hygiene*, 55(1):63–68.

McLean, R. G., Szmyd, D. M., and Calisher, C. H. 1982. Experimental studies of Rio Grande virus in rodent hosts. *The American Journal of Tropical Medicine and Hygiene*, 31(3):569–573.

McMillan, B. and Boulger, L. R. 1960. The susceptibility of the ground-squirrel Xerus (Euxerus) Erythropus geoffroy, 1803, to rabies street virus and its potentiality as a reservoir of rabies in northern Nigeria. *Annals of Tropical Medicine & Parasitology*, 54(2):165–171.

Middleton, D., Morrissy, C., Van Der Heide, B., Russell, G., Braun, M., Westbury, H., Halpin, K., and Daniels, P. 2007. Experimental Nipah virus infection in pteropid bats (Pteropus poliocephalus). *Journal of Comparative Pathology*, 136(4):266–272.

Moreno, J. A. and Baer, G. M. 1980. Experimental babies in the vampire bat. *The American Journal of Tropical Medicine and Hygiene*, 29(2):254–259.

Neill, W. A. and Kading, R. C. 2021. Viral ecology and natural infection dynamics of Kaeng Khoi virus in cave-dwelling wrinkle-lipped free-tailed bats (Chaerephon plicatus) in Thailand. *Diseases*, 9(4):73.

Nosek, J., Gresikova, M., and Rehacek, J. 1961. Persistence of tick-borne encephalitis virus in hibernating bats. *Acta Virologica*, 2:112–116.

Nosek, J., Kozuch, O., Lichard, M., Ernek, E., and Albrecht, P. 1963. Experimental infection of the great dormouse (Glis glis) with tick-borne encephalitis virus. *Acta Virologica*, 7:374–376.

Obregón-Morales, C., Aguilar-Setién, Á., Perea Martínez, L., Galvez-Romero, G., Martínez-Martínez, F. O., and Aréchiga-Ceballos, N. 2017. Experimental infection of Artibeus intermedius with a vampire bat rabies virus. *Comparative Immunology, Microbiology and Infectious Diseases*, 52:43–47.

Oelofsen, M. J. and Van Der Ryst, E. 1999. Could bats act as reservoir hosts for Rift Valley fever virus? *Onderstepoort Journal of Veterinary Research*, 66:51–54.

Pantuwatana, S., Thompson, W. H., Watts, D. M., and Hanson, R. P. 1972. Experimental infection of chipmunks and squirrels with la crosse and trivittatus viruses and biological transmission of la crosse virus by aedes triseriatus. *The American Journal of Tropical Medicine and Hygiene*, 21(4):476–481.

Pavri, K. M. and Singh, K. R. 1965. Demonstration of antibodies against the virus of Kyasanur Forest disease (KFD) in the frugivorous bat Rousettus leschenaulti, near Poona, India. *The Indian Journal of Medical Research*, 53(10):956–961.

Pavri, K. M. and Singh, K. R. 1968. Kyasanur forest disease virus infection in the frugivorous bat, Cynopterus sphinx. *The Indian Journal of Medical Research*, 56(8):1202–1204.

Pawan, J. L. 1936. Rabies in the vampire bat of Trinidad, with special reference to the clinical course and the latency of infection. *Annals of Tropical Medicine & Parasitology*, 30(4):401–422.

Pawan, J. L. 1948. Fruit-eating bats and paralytic rabies in Trinidad. *Annals of Tropical Medicine & Parasitology*, 42(2):173–177.

Paweska, J. T., Jansen Van Vuren, P., Masumu, J., Leman, P. A., Grobbelaar, A. A., Birkhead, M., Clift, S., Swanepoel, R., and Kemp, A. 2012. Virological and serological findings in Rousettus aegyptiacus experimentally inoculated with Vero cells-adapted Hogan strain of marburg virus. *PLoS ONE*, 7(9): e45479.

Paweska, J. T., Jansen Van Vuren, P., Fenton, K. A., Graves, K., Grobbelaar, A. A., Moolla, N., Leman, P., Weyer, J., Storm, N., McCulloch, S. D., Scott, T. P., Markotter, W., Odendaal, L., Clift, S. J., Geisbert, T. W., Hale, M. J., and Kemp, A. 2015. Lack of marburg virus transmission From experimentally infected to susceptible in-contact Egyptian fruit bats. *Journal of Infectious Diseases*, 212(suppl 2):S109–S118.

Perea-Martínez, L., Moreno-Sandoval, H., Moreno-Altamirano, M., Salas-Rojas, M., García-Flores, M., Aréchiga-Ceballos, N., Tordo, N., Marianneau, Ph., and Aguilar-Setién, A. 2013. Experimental infection of Artibeus intermedius bats with serotype-2 dengue virus. *Comparative Immunology, Microbiology and Infectious Diseases*, 36(2):193–198.

Platt, K. B., Tucker, B. J., Halbur, P. G., Tiawsirisup, S., Blitvich, B. J., Fabiosa, F. G., Bartholomay, L. C., and Rowley, W. A. 2007. West Nile virus viremia in eastern chipmunks (Tamias striatus) sufficient for infecting different mosquitoes. *Emerging Infectious Diseases*, 13(6):831–837.

Platt, K. B., Tucker, B. J., Halbur, P. G., Blitvich, B. J., Fabiosa, F. G., Mullin, K., Parikh, G. R., Kitikoon, P., Bartholomay, L. C., and Rowley, W. A. 2008. Fox squirrels (Sciurus niger) develop West Nile virus viremias sufficient for infecting select mosquito species. *Vector-Borne and Zoonotic Diseases*, 8(2):225–234.

Pretorius, A., Oelofsen, M. J., Smith, M. S., and Van Der Ryst, E. 1997. Rift Valley fever virus: A seroepidemiologic study of small terrestrial vertebrates in South Africa. *The American Journal of Tropical Medicine and Hygiene*, 57(6):693–698.

Reagan, R. L., Yancey, F. S., and Brueckner, A. L. 1957. Transmission of rabies from artificially infected bats to Syrian hamsters. *A.M.A. Archives of Pathology*, 63(3):278–280.

Reagan, R. and Brueckner, A. L. 1951. Response of the cave bat (Myotus lucifugus) to Newcastle disease virus by vairous methods of exposure. *The Cornell Veterinarian*, pages 56–57.

Ressang, A., Gwan, S. I., and Hardjosworo, S. 1963. Bats, rodents and rabies in Indonesia. *Communicationes veterinariae*, 7(22):33–41.

Rodrigues, Y. J. L. and Tamayo, J. G. 2000. Pathology of experimental infection with the rabies virus in hematophagous bat (Desmodus rotundus). *Revista de la Facultad de Ciencias Veterinarias*, 41:71–72.

Root, J. J., Oesterle, P. T., Nemeth, N. M., Klenk, K., Gould, D. H., Mclean, R. G., Clark, L., and Hall, J. S. 2006. Experimental infection of fox squirrels (Sciurus niger) with West Nile virus. *The American Journal of Tropical Medicine and Hygiene*, 75(4):697–701.

Sbrana, E., Xiao, S.-Y., Newman, P. C., and Tesh, R. B. 2007. Comparative pathology of North American and Central African strains of monkeypox virus in a ground squirrel model of the disease. *The American Journal of Tropical Medicine and Hygiene*, 76(1):155–164.

Schlottau, K., Rissmann, M., Graaf, A., Schön, J., Sehl, J., Wylezich, C., Höper, D., Mettenleiter, T. C., Balkema-Buschmann, A., Harder, T., Grund, C., Hoffmann, D., Breithaupt, A., and Beer, M. 2020. SARS-CoV-2 in fruit bats, ferrets, pigs, and chickens: An experimental transmission study. *The Lancet Microbe*, 1(5):e218–e225.

Seamer, J., Fitzgeorge, R., and Smith, C. E. G. 1967. Resistance of the short-tailed vole Microtus agrestis (L.) to infection with two arboviruses. *The British Journal of Experimental Pathology*, 48(5):463–467.

Sétien, A. A., Brochier, B., Tordo, N., De Paz, O., Desmettre, P., Péharpré, D., and Pastoret, P.-P. 1998. Experimental rabies infection and oral vaccination in vampire bats (Desmodus rotundus). *Vaccine*, 16 (11-12):1122–1126.

Seymour, C., Dickerman, R. W., and Martin, M. S. 1978. Venezuelan encephalitis virus infection in neotropical bats: I. Natural infection in a guatemalan enzootic focus. *The American Journal of Tropical Medicine and Hygiene*, 27(2):290–296.

Seymour, C., Amundson, T. E., Yuill, T. M., and Bishop, D. H. L. 1983. Experimental infection of chipmunks and snowshoe hares with La Crosse and snowshoe hare viruses and four of their reassortants. *The American Journal of Tropical Medicine and Hygiene*, 32(5):1147–1153.

Shah, K. V. and Daniel, R. W. 1966. Attempts at experimental infection of the Indian fruit-bat Pteropus giganteus with chikungunya and dengue 2 viruses and antibody survey of bat sera for some viruses. *The Indian Journal of Medical Research*, 54(8):714–22.

Sims, R. A., Allen, R., and Sulkin, S. E. 1963. Studies on the pathogenesis of rabies in insectivorous Bats*: III. Influence of the gravid state. *The Journal of Infectious Diseases*, 112(1):17–27.

Storm, N., Jansen Van Vuren, P., Markotter, W., and Paweska, J. 2018. Antibody responses to marburg virus in Egyptian rousette bats and their role in protection against infection. *Viruses*, 10(2):73.

Sulkin, S. E., Krutzsch, P. H., Wallis, C., and Allen, R. 1957. Role of brown fat in pathogenesis of rabies in insectivorous bats (Tadarida b. mexicana). *Experimental Biology and Medicine*, 96(2):461–464.

Sulkin, S. E., Sims, R., and Allen, R. 1964. Studies of arthropod-borne virus infections in chiroptera II. Experiments with Japanese B and St. Louis encephalitis viruses in the gravid bat. Evidence of transplacental transmission. *The American Journal of Tropical Medicine and Hygiene*, 13:475–481.

Sulkin, S. E. 1962. Bat rabies: Experimental demonstration of the ”reservoiring mechanism”. *American Journal of Public Health and the Nations Health*, 52(3):489–498.

Sulkin, S. E., Krutzsch, P. H., Allen, R., and Wallis, C. 1959. Studies on the pathogenesis of rabies in insectivorous bats: I. Role of brown adipose tissue. *Journal of Experimental Medicine*, 110(3):369–388.

Sulkin, S. E., Allen, R., Sims, R., Krutzsch, P. H., and Kim, C. 1960. Studies on the pathogenesis of rabies in insectivorous bats : II. Influence of environmental temperature. *Journal of Experimental Medicine*, 112 (4):595–617.

Suu-Ire, R., Begeman, L., Banyard, A. C., Breed, A. C., Drosten, C., Eggerbauer, E., Freuling, C. M., Gibson, L., Goharriz, H., Horton, D. L., Jennings, D., Kuzmin, I. V., Marston, D., Ntiamoa-Baidu, Y., Riesle Sbarbaro, S., Selden, D., Wise, E. L., Kuiken, T., Fooks, A. R., Müller, T., Wood, J. L. N., and Cunningham, A. A. 2018. Pathogenesis of bat rabies in a natural reservoir: Comparative susceptibility of the straw-colored fruit bat (Eidolon helvum) to three strains of Lagos bat virus. *PLOS Neglected Tropical Diseases*, 12(3):e0006311.

Swanepoel, R., Blackburn, N. K., Efstratiou, S., and Condy, J. B. 1978. Studies on Rift Valley fever in some African murids (Rodentia: Muridae). *Journal of Hygiene*, 80(2):183–196.

Swanepoel, R., Leman, P. A., Burt, F. J., Zachariades, N. A., Braack, L. E., Ksiazek, T. G., Rollin, P. E., Zaki, S. R., and Peters, C. J. 1996. Experimental inoculation of plants and animals with ebola virus. *Emerging Infectious Diseases*, 2(4):321–325.

Tesh, R. B., Watts, D. M., Sbrana, E., Siirin, M., Popov, V. L., and Xiao, S.-Y. 2004. Experimental infection of ground squirrels (Spermophilus tridecemlineatus) with monkeypox Virus. *Emerging Infectious Diseases*, 10(9):1563–1567.

Tiawsirisup, S., Blitvich, B. J., Tucker, B. J., Halbur, P. G., Bartholomay, L. C., Rowley, W. A., and Platt, K. B. 2010. Susceptibility of fox squirrels (Sciurus niger) to West Nile virus by oral exposure. *Vector-Borne and Zoonotic Diseases*, 10(2):207–209.

Turmelle, A. S., Jackson, F. R., Green, D., McCracken, G. F., and Rupprecht, C. E. 2010. Host immunity to repeated rabies virus infection in big brown bats. *Journal of General Virology*, 91(9):2360–2366.

Watanabe, S., Omatsu, T., Miranda, M. E., Masangkay, J. S., Ueda, N., Endo, M., Kato, K., Tohya, Y., Yoshikawa, Y., and Akashi, H. 2010. Epizootology and experimental infection of Yokose virus in bats. *Comparative Immunology, Microbiology and Infectious Diseases*, 33(1):25–36.

Williamson, M., Hooper, P., Selleck, P., Gleeson, L., Daniels, P., Westbury, H., and Murray, P. 1998. Transmission studies of Hendra virus (equine morbilli-virus) in fruit bats, horses and cats. *Australian Veterinary Journal*, 76(12):813–818.

Williamson, M., Hooper, P., Selleck, P., Westbury, H., and Slocombe, R. 1999. Experimental hendra virus infection in pregnant guinea-pigs and fruit bats (Pteropus poliocephalus). *Journal of Comparative Pathology*, 122:201–207.

Woon, A. P., Boyd, V., Todd, S., Smith, I., Klein, R., Woodhouse, I. B., Riddell, S., Crameri, G., Bingham, J., Wang, L.-F., Purcell, A. W., Middleton, D., and Baker, M. L. 2020. Acute experimental infection of bats and ferrets with Hendra virus: Insights into the early host response of the reservoir host and susceptible model species. *PLOS Pathogens*, 16(3):e1008412.

Xiao, S.-Y., Sbrana, E., Watts, D. M., Siirin, M., Travassos Da Rosa, A. P., and Tesh, R. B. 2005. Experimental infection of prairie dogs with monkeypox virus. *Emerging Infectious Diseases*, 11(4):539–545.

Young, N. A., Johnson, K. M., and Gauld, L. W. 1969. Viruses of the Venezuelan equine encephalomyelitis complex. Experimental infection of Panamanian rodents. *The American Journal of Tropical Medicine and Hygiene*, 18(2):290–296.
